# Structural basis for the assembly and electron transport mechanisms of the dimeric photosynthetic RC–LH1 supercomplex

**DOI:** 10.1101/2021.12.17.473239

**Authors:** Peng Cao, Laura Bracun, Atsushi Yamagata, Bern M. Christianson, Tatsuki Negami, Baohua Zou, Tohru Terada, Daniel P. Canniffe, Mikako Shirouzu, Mei Li, Lu-Ning Liu

**Author notes:** Correspondence to (M.S.), (M.L.), (L.-N.L.). The authors contribute equally to this work (Peng Cao, Laura Bracun, Atsushi Yamagata). Department of Chemistry and Biology, Faculty of Environment and Life Science, Beijing University of Technology, Beijing 100124, China (Peng Cao).

## Abstract

The reaction center (RC) and light-harvesting complex 1 (LH1) form a RC–LH1 core supercomplex that is vital for the primary reactions of photosynthesis in purple photosynthetic bacteria. Some species possess the dimeric RC–LH1 complex with an additional polypeptide PufX, representing the largest photosynthetic complex in anoxygenic phototrophs. However, the details of the architecture and assembly mechanism of the RC–LH1 dimer are unclear. Here we report seven cryo-electron microscopy (cryo-EM) structures of RC–LH1 supercomplexes from *Rhodobacter sphaeroides*. Our structures reveal that two PufX polypeptides are positioned in the center of the S-shaped RC–LH1 dimer, interlocking association between the components and mediating RC–LH1 dimerization. Moreover, we identify a new transmembrane peptide, designated PufY, which is located between the RC and LH1 subunits near the LH1 opening. PufY binds a quinone molecule and prevents LH1 subunits from completely encircling the RC, creating a channel for quinone/quinol exchange. Genetic mutagenesis, cryo-EM structures, and computational simulations enable a mechanistic understanding of the assembly and electron transport pathways of the RC–LH1 dimer and elucidate the roles of individual components in ensuring the structural and functional integrity of the photosynthetic supercomplex.

## INTRODUCTION

Photosynthesis converts solar radiation into chemical energy to power almost all life on Earth ^1^. The photosynthetic systems of purple bacteria represent a model for exploring the assembly and function of photosynthetic apparatus. In purple phototrophic bacteria, the early stage of photosynthesis occurs in the reaction center–light-harvesting 1 (RC–LH1) core supercomplexes that are accommodated within the intra-cytoplasmic membranes (termed chromatophores) ^1,2^. The RC–LH1 supercomplex comprises an RC surrounded by the LH1 ring that is made of multiple αβ-heterodimers ^3,4^. LH1 collects photons directly from sunlight or from the peripheral antenna, light-harvesting complex 2 (LH2), and transfers excitation energy to the RC. The subsequent photo-induced charge separation in the RC initiates a cyclic electron transfer between the RC and cytochrome *bc*_1_ (Cyt *bc*_1_) complex, which ultimately creates proton gradients across the chromatophore membrane to generate ATP ^5,6^.

The compositions and architectures of RC–LH1 supercomplexes exhibit a diversity among various bacterial species ^2^. Many RC–LH1 complexes occur as monomers, in which the RC is encircled by a closed LH1 ring of 16 αβ-heterodimers ^7–11^, or an open LH1 ring of 15-16 αβ-polypeptides with a gap formed by specific transmembrane (TM) polypeptides ^12–17^. For instance, the PufX polypeptide in the *Rhodobacter* (*Rba*.) *veldkampii* RC–LH1 monomer functions as a molecular “cross brace” to stabilize the RC–LH1 core complex and mediate a large opening in the LH1 ring for quinone/quinol exchange ^17^. In contrast, most of the *Rhodobacter* species, exemplified by *Rba*. *sphaeroides*, possess the dimeric RC–LH1 core structures, representing the largest RC–LH1 supercomplexes in bacterial photosynthesis. Previous electron microscopy and X-ray crystallography studies have shown that the *Rba. sphaeroides* RC–LH1 dimer contains two RCs surrounded by an S-shaped LH1 ring with 28 LH1 αβ-heterodimers plus 2 PufX polypeptides ^18–23^. However, detailed information about the modular structures of the dimeric RC–LH1 complexes that mediate their stepwise assembly and electron flow remain uncharacterized.

Here, we report cryo-electron microscopy (cryo-EM) structures of RC–LH1 complexes from the model purple phototrophic bacterium *Rba*. *sphaeroides* DSM 158 (strain 2.4.1). These high-resolution structures reveal in detail how PufX and a new TM polypeptide, PufY, are associated with the RC–LH1 complex and how they mediate dimerization, LH1 encirclement, and quinone transport through the RC–LH1 core supercomplex.

## RESULTS AND DISCUSSION

### Overall structures of the RC–LH1 supercomplexes

Wild-type (WT) *Rba*. *sphaeroides* DSM 158 contains both RC–LH1 monomers and dimers (Fig. S1), corroborated by atomic force microscopy (AFM) imaging of isolated photosynthetic membranes ^24,25^. The cryo-EM structure of the RC–LH1 monomer (33 polypeptides, 76 cofactors) at 2.79 Å resolution shows that the RC is surrounded by an open LH1 ring of 14 αβ-subunits, with a large gap (46 Å) formed by a PufX TM polypeptide that has a tilted angle to the membrane plane (Fig. 1a, Figs. S2-S4, Tables S1, S2). The RC comprises the H, L and M subunits, a bacteriochlorophyll (BChl) *a* dimer as the special pair to act as the primary electron donor, two BChl *a* monomers, two bacteriopheophytins (BPhes), one spheroidene (SPO) carotenoid, and a Fe^2+^ ion, similar to the reported crystal structure ^26^. The monomeric RC–LH1 structure resembles the architecture of the RC–LH1–PufX monomer from *Rba*. *veldkampii* ^17^, except for a missing LH1 subunit and newly identified densities (assigned as PufY, see details below) that are located between the RC and LH1, on the opposite side of PufX near the LH1 opening (Fig. S5).

**Fig. 1.**
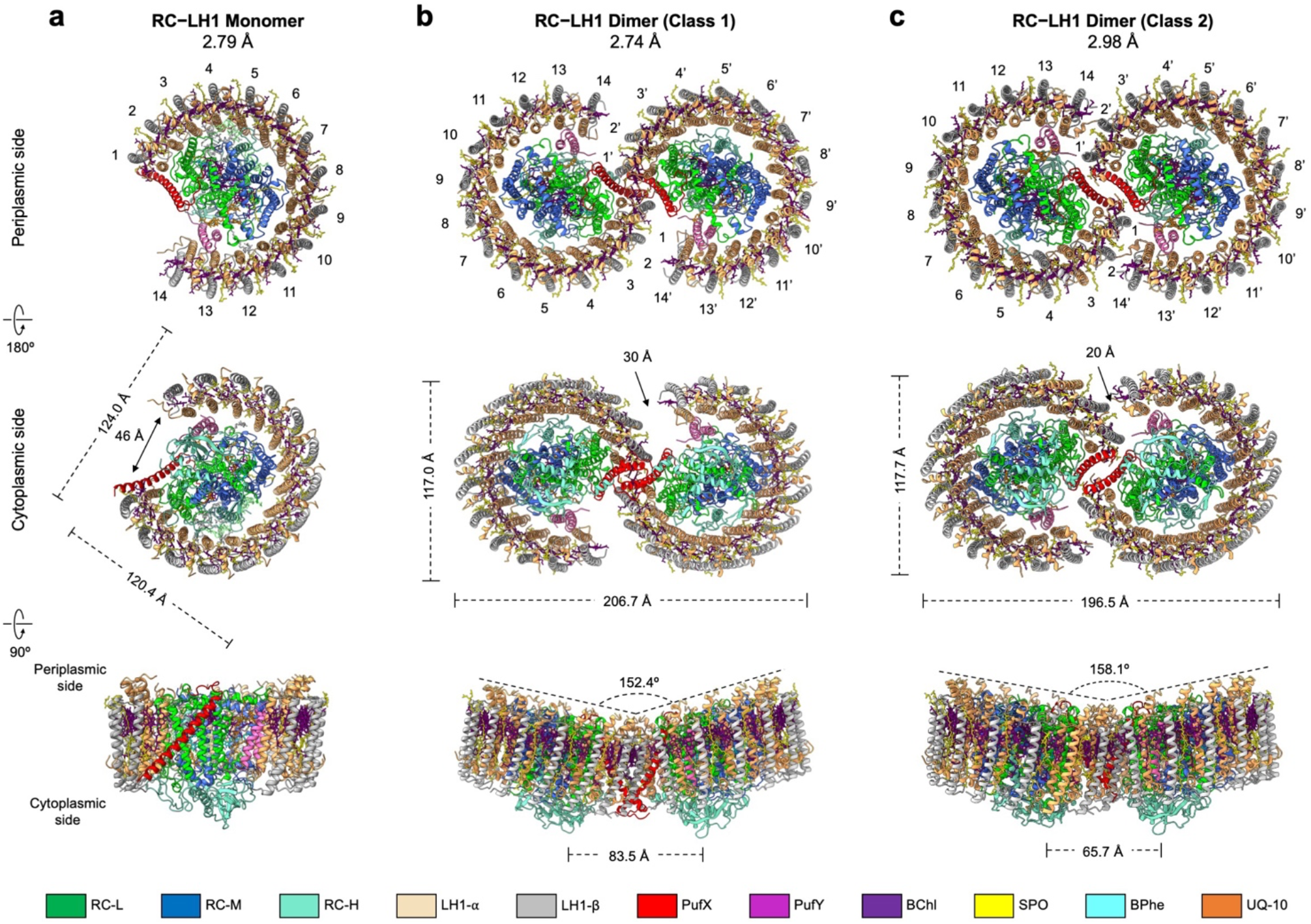
Cryo-EM structures of the RC–LH1 core supercomplexes from *Rba*. *sphaeroides*. (**a**) The RC–LH1 monomer. Top, view from the periplasmic side; middle, view from the periplasmic side; bottom, side view in the membrane plane (facing the PufX-mediated gap region). (**b**) The Class-1 RC–LH1 dimer. Top, view from the periplasmic side; middle, a view from the periplasmic side; bottom, side view in the membrane plane. (**c**) The Class-2 RC–LH1 dimer. Top, a view from the periplasmic side; middle, view from the periplasmic side; bottom, side view in the membrane plane. Color scheme is presented at the bottom as follows: LH1-α, wheat; LH1-β, grey; PufX, red; PufY, purple; RC-L, green; RC-M, marine; RC-H, teal; BChls, purple; BPhes, cyan; carotenoids, yellow; quinones, orange.

Cryo-EM single-particle analysis determined two distinct classes of RC–LH1 dimers (Class-1 and Class-2) at 2.74 Å and 2.98 Å resolution, respectively (Fig. 1b, 1c, Figs. S6, S7, Table S1). The dimers of both classes adopt a two-fold symmetry composed of two identical monomers. The structures of the monomers within the RC–LH1 dimers of both classes and the WT RC–LH1 monomer are closely comparable (Fig. S8).

The Class-1 dimer comprises 66 polypeptides and 148 cofactors, with a molecular mass of 558.4 kDa; the long and short dimensions are 206.7 Å and 117.0 Å, respectively (Fig. 1b, Table S2). The LH1 subunits, composed of 28 αβ-polypeptide pairs, 56 BChls *a* and 50 SPOs, form an S-shaped structure made up of two C-shaped LH1 rings, with two openings of ~30 Å in distance, encircling the two RCs. In addition, two PufX polypeptides are located at the dimerization interface in the center of the RC–LH1 complex, and two PufY polypeptides are located near the two LH1 openings. The two monomers exhibit a tilted angle of 152° measured at the periplasmic face of the complex. The intrinsic bent architecture of the RC–LH1 dimer could create the local membrane curvature, supported by all-atom molecular dynamics (AAMD) simulations (Fig. S9).

Compared with the Class-1 dimer, the Class-2 dimer (66 polypeptides, 152 cofactors) displays a slightly reduced length spanning the dimer (196.5 Å) and a tilted angle of 158° (Fig. 1c, Table S2); rotational and lateral shifts at the monomer-monomer interface were observed (Fig. S10). Consequently, the continuous S-shaped LH1 array “breaks” in the middle, resulting in two separated C-shaped LH1 rings with a narrower opening (~20 Å) in the LH1 array than that of the Class-1 dimer. Three ubiquinone molecules (Q_A_, Q_B_, Q_3_) were identified in each monomer of the RC-LH1 supercomplexes, tentatively assigned as ubiquinone-10 (UQ-10) according to the HPLC analysis (Figs. S1, S11). In addition, an extra putative UQ-10 (named as Q_Y_) that is strongly associated with PufY was identified in both the Class-2 dimer and WT monomer (Fig. S11).

### Structure of the LH1 ring

Both N- and C-termini of LH1-α form short helices, which are hydrogen bonded with LH1-β_n+1_ and LH1-β_n-1_, respectively (Fig. S12). LH1-α_n_ and LH1-α_n-1_ also form hydrogen bonds through Tyr41-Arg53 residues at the periplasmic side. The RC–LH1 interactions within the dimeric and monomeric structures (Fig. S13, Table S3) resemble those identified in the *Rba*. *veldkampii* RC–LH1–PufX ^17^.

A total of 28 BChls *a* and 25 SPOs were resolved in the LH1 ring of each monomer. The LH1 αβ-heterodimer sandwiches 2 BChls and 2 SPOs, consistent with previous estimation ^18,27^, except for the two LH1 pairs close to the gap: LH1-1 lacks one SPO and LH1-14 has no SPO (Fig. S14, S15). The two carotenoids, SPO-α and SPO-β, mediate the associations between LH1_n_ and LH1_n+1_ and between LH1_n_ and LH1_n-1_, respectively (Fig. S16). SPO-α projects into the TM region between α- and β-polypeptides and adopts a similar conformation to the carotenoids in the LH1 of other purple bacteria ^3,7–9,12–17^. SPO-β, which has not been observed in previously reported RC–LH1 structures, is located at the periplasmic layer between LH1-β_n_ and LH1-β_n+1_ (Fig. S16). The additional SPO provides a tightly arranged pigment network within the LH1 barrier (Fig. S16), reducing the possibility for traffic of quinones/quinols through the channels between LH1 subunits as proposed in other RC– LH1 complexes ^7,8,16,17^ and emphasizing the necessity of forming an open LH1 ring in the RC–LH1 dimer and monomer.

The 28 BChl molecules in each monomer encircle an open ring at the periplasmic side (Fig. 1). Comparable Mg–Mg distances were found between the BChls in each LH1 αβ-subunit (9.1–9.4 Å) and of adjacent pairs (~8.5 Å) (Fig. S10). BChls in the Class-1 dimer form a continuous S-shaped array, ensuring that solar energy absorbed by the LH1 BChls in one monomer can be transferred to the LH1 of the neighboring monomer. This arrangement may provide the foundation for equilibrated, efficient exciton coupling and energy trapping ^28^. In contrast, the continuity of the array of excitonically coupled BChls is interrupted at the monomer-monomer interface in the Class-2 dimer, resulting in a ~23-Å gap between adjacent BChls from the two monomers (Figs. S10, S11).

### Location of PufX and its interactions within the RC–LH1 dimer

PufX (82 amino acids) has been proposed to play roles in dimerization of RC–LH1 and exchange of quinones/quinols between the RC and Cyt *bc*_1_ ^22,23,29,30^. However, its precise structure and location within the RC–LH1 dimer have remained elusive. Our cryo-EM structures reveal that two PufX polypeptides are positioned at the dimerization interface of the RC–LH1 supercomplex (Fig. 2), in agreement with AFM and low-resolution EM structures ^23,31,32^ but distinct from the model inferred from an 8-Å crystal structure (PDB ID: 4V9G), which proposed that PufX is positioned in the gap next to the terminal LH1 subunit and adjacent to the Q_B_ site of the RC ^18^. The N-terminus of PufX is exposed on the cytoplasmic side of the complex and its C-terminal domain is situated on the periplasmic surface (Fig. 2a, 2b). The PufX peptide is tilted ~45° from the RC to the peripheral side of LH1 and adopts a slightly bent configuration in the Ala24–Gly36 region within the membrane layer, resembling the PufX structure from *Rba*. *veldkampii* ^17^ (Fig. S17). The crossing angle for the PufX dimer is ~73° (Fig. 2b).

**Fig. 2.**
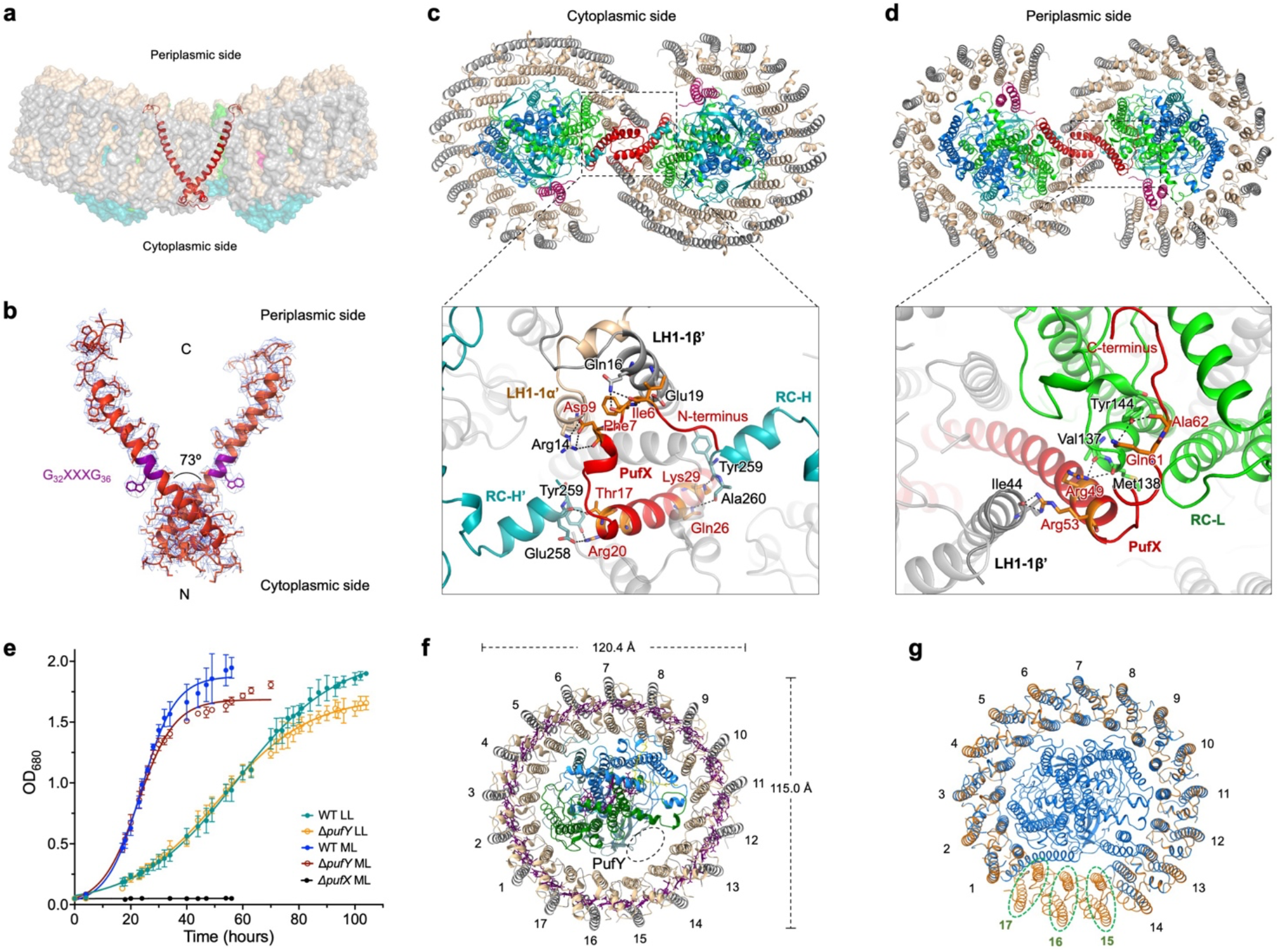
The location of PufX and interactions within the RC–LH1 dimer. (**a**) Two PufX polypeptides (red cartoon) are positioned at the monomer–monomer interface of the RC–LH1 dimer. (**b**) Cryo-EM densities of the PufX dimer. The G_32_XXXG_36_ motif within the TM helix of PufX, which is known to drive TM helix-helix association ^42^, does not feature the dimerization interface of the PufX dimer, and thus does not possibly mediate PufX–PufX dimerization. (**c**) Interactions between PufX and the RC–LH1 subunits (RC-H, LH1-1) at the cytoplasmic side. The residues of PufX, RC-H, LH1-1α and LH1-1β involved in the inter-subunit interactions are shown in sticks and colored in orange, teal, wheat, and grey, respectively. All the interacting residues involved in the association between PufX and the RC-L subunit are listed in Table S4. (**d**) Interactions between PufX and the RC–LH1 subunits (RC-L, LH1-1) at the periplasmic side. The residues of PufX, RC-L and LH1-1β involved in the inter-subunit interactions are shown in sticks and colored in orange, green, and grey, respectively. (**e**) Photosynthetic growth of the *Rba*. *sphaeroides* wild-type (WT), Δ*pufX*, and Δ*pufY* mutants under low light (LL, 5 μmol photons s^-1^ m^-2^) and moderate light (ML, 25 μmol photons s^-1^ m^-2^), monitored at OD_680_ and fitted with logistic growth using Prism. The growth rate is 0.16 h^-1^ for WT and 0.15 h^-1^ for Δ*pufY* under ML, and 0.06 h^-1^ for WT and 0.06 h^-1^ for Δ*pufY* under LL. (**f**) Cartoon representation of the Δ*pufX* RC– LH1 complex. Without PufX, LH1 forms a complete circle of 17 subunits, surrounding the RC. Mass spectrometry confirmed the presence of PufY in the Δ*pufX* RC–LH1 monomer but PufY exhibits weak densities in the Δ*pufX* RC–LH1 structure. Thus, the potential location of PufY is indicated by a dashed circle. (**g**) Comparison of the structures of the Δ*pufX* RC–LH1_17_ monomer (yellow) and the wild-type RC–LH1_14_ monomer (blue) from *Rba*. *sphaeroides*. To highlight the differences in the LH1 architectures, the RC subunits of Δ*pufX* RC–LH1_17_ were not shown. The three extra LH1 subunits within the Δ*pufX* RC–LH1_17_ monomer are circled.

The N-terminal region of PufX (Ala2-Pro15) is well resolved in the Class-1 dimeric structure, revealing that residues Ile6-Asp9 interact with LH1-1αβ of the other monomer via hydrogen bonds and salt bridges (Fig. 2c, Table S4). The N-terminal domain of the TM helix of PufX (Thr17-Lys29) forms hydrogen bonds with both RC-H subunits of the dimer at the cytoplasmic side. The N-termini of the two PufX polypeptides are in close proximity to each other at the monomer-monomer interface, albeit without detectable interactions. It is likely that water molecules, which were not identified at the present resolution, mediate PufX–PufX interactions at the N-terminal region. In contrast, the N-terminal domain of PufX (Ala2-Pro15) is not resolved in the structures of the monomer and Class-2 dimer (Fig. S8), presumably due to its disordered structure when not binding with RC-H and LH1-1 subunits of the neighboring monomer. In Class-2, differences in the RC–LH1 association occur at the dimerization interface, and the two N-terminal heads of PufX are further separated, due to the twist of two RC–LH1 monomers, compared with those in Class-1 (Figs. S18, S19).

At the periplasmic side, the C-terminal regions of the two PufX polypeptides are separated by LH1-1 from each monomer and have no close contacts with each other (Fig. 2d). The Arg49-Ala62 residues interacts with RC-L (Val137-Tyr144) within the same monomer and LH1-1β of the neighboring monomer through hydrogen bonds. Due to steric hindrance provided by PufX within the same monomer, and interactions with the PufX of the adjacent monomer, LH1-1α displays a conformational shift compared with the other LH1 subunits and lacks SPO-β; moreover, both SPO-α and the phytol tail of BChl-α kink at the periplasmic side to interact with PufX (Fig. S14). All these interactions formed by the N- and C-termini of PufX and LH1 interlock the RC–LH1 structures and mediate dimerization of RC–LH1, providing the structural evidence for previous observations ^33–36^.

To verify the roles of PufX in the RC–LH1 dimer, we genetically deleted the *pufX* gene, which resulted in the exclusive formation of RC–LH1 monomers and loss of capability to perform photoheterotrophic growth (Fig. 2e, Fig. S1), consistent with previous results ^34,37–39^. The cryo-EM structure of the Δ*pufX* RC–LH1 monomer at 4.20-Å resolution shows that the central RC is surrounded by a completely closed LH1 ring of 17 subunits, adopting a slightly elliptical shape (120 Å × 115 Å) (Figs. 2f, 2g, Figs. S20, S21, Table S1). The cryo-EM structure explains the previous observation that the absence of PufX resulted in an increase in the ratio of LH1 to RC ^40^. The closed LH1 ring and the tightly packed pigments within LH1s impede quinone/quinol exchange, thereby causing the incapability of this mutant strain to grow photosynthetically. We could not obtain a higher-resolution cryo-EM structure of the Δ*pufX* RC–LH1_17_ monomer, and the RC and PufY exhibit weaker densities than the peripheral LH1. Mass spectrometry confirmed that ~27% of PufY was retained in the *ΔpufX* RC–LH1 monomer compared to the WT monomer. It is probably because the lack of intensive interactions between PufX and the RC–LH1 subunits, which may confine the orientation of the RC inside the LH1 ring ^13^ and stabilize the association between RC, PufY, and LH1, could result in an unstable assembly of RC–LH1 complexes and formation of LH1-only rings, as suggested previously ^38,41^.

### A new transmembrane polypeptide PufY

Another distinctive feature of the *Rba*. *sphaeroides* RC–LH1 structures is the presence of extra densities at the interface between the RC and LH1-13 and LH-14 subunits, adjacent to the RC Q_B_ site (Fig. 3a). This position is close to the location of the previously assigned PufX polypeptide in the low-resolution crystal and cryo-EM structures ^18,20^. We identified this protein as RSP_7571 (termed PufY, equivalent to the Protein-U ^43^ or Protein-Y ^44^ reported recently), based on the perfect match of its sequence with the cryo-EM density map (Fig. 3b, Fig. S4) and mass spectrometry results. PufY comprises 53 residues with two parallel TM helices connected by a short loop (Fig. 3b, Fig. S22). It has a molecular mass of 5,553 Da and migrates at the same position as LH1-β (5,588 Da) in SDS-PAGE (Fig. S1), explaining why it has not been identified in previous studies. Unlike *pufX*, the *rsp_7571* gene encoding PufY is not in proximity to the photosynthesis gene clusters (Fig. S22). Bioinformatic analysis revealed both the protein sequence of PufY and genomic location are highly conserved in all *Rba. sphaeroides* subspecies and several additional *Rhodobacter* species (Fig. S22).

**Fig. 3.**
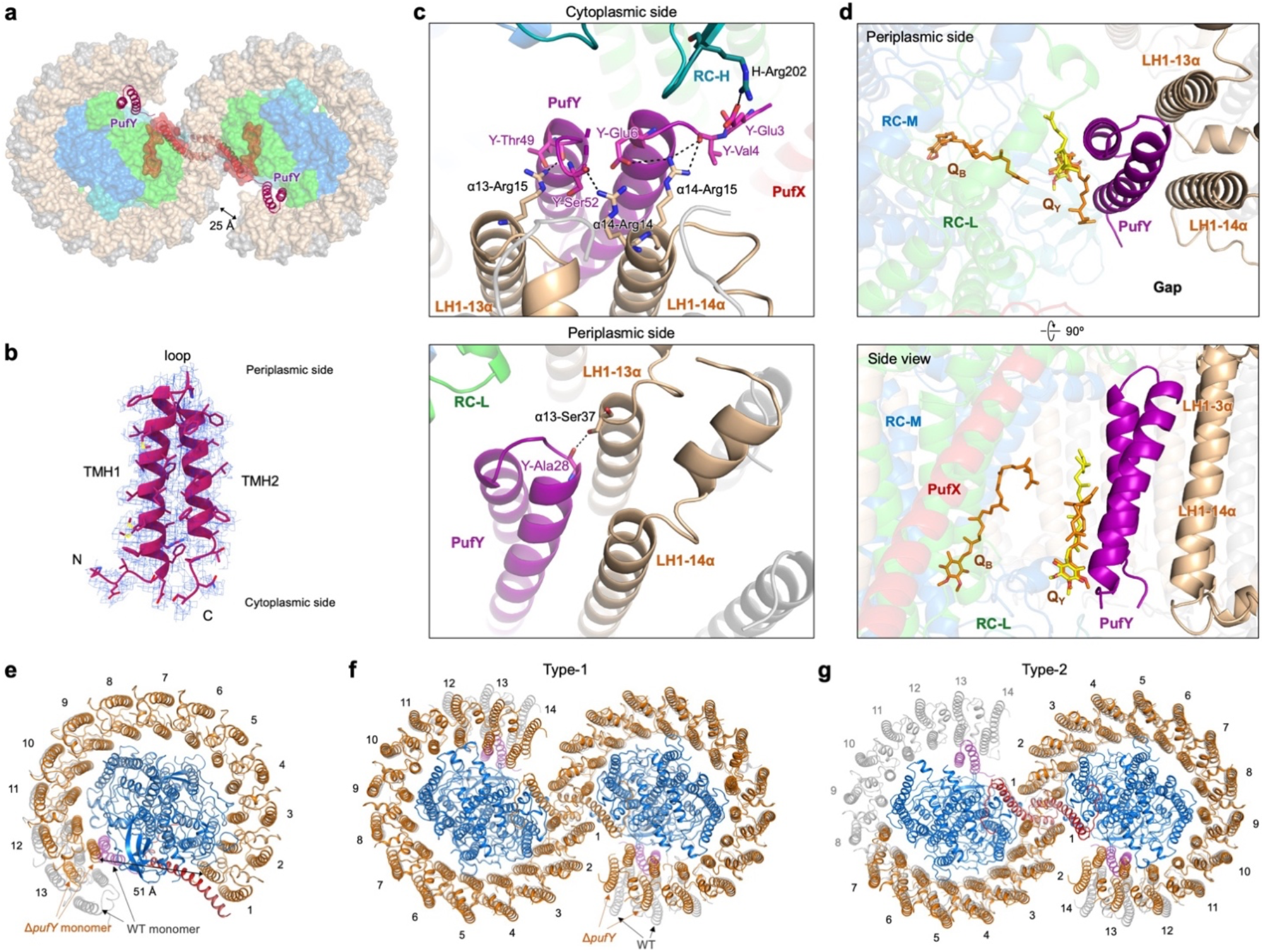
The PufY location and interactions within the RC–LH1 dimer. (**a**) Two PufY polypeptides (purple cartoon) are positioned between the RC and LH1, close to the gaps of the RC–LH1 dimer (periplasmic view). (**b**) Cryo-EM density of PufY that comprises two transmembrane helices connected by a short loop. (**c**) Interactions between PufY and the RC–LH1 subunits. At the cytoplasmic side (top), the N- and C-terminal residues of PufY form hydrogen bonds with RC-H, LH1-13α, and LH1-14α. At the periplasmic side (bottom), PufY is hydrogen bonded with LH1-13α. All the interacting residues involved in the association between PufY and RC–LH1 are listed in Table S5. (**d**) A quinone molecule (Q_Y_) is identified in the structures of the Class-2 dimer (yellow) and wild-type monomer (orange), which binds with PufY near the RC Q_B_ site. (**e**) Cryo-EM structure of the Δ*pufY* RC–LH1 monomer (cytoplasmic view, LH1 shown in orange cartoon) shows 13 LH1 subunits surrounding the RC, interlocked by PufX. Comparison of the structures of the Δ*pufY* RC–LH1 monomer and the wild-type RC–LH1_14_ monomer (LH1 shown in grey cartoon, PufY shown in purple cartoon) shows that lack of PufY induces the shift of terminal LH1 subunits LH1-12 and LH1-13 towards the RC, and missing of the 14^th^ LH1 subunit creates a larger opening in the LH1 ring of the Δ*pufY* RC–LH1 monomer. (**f**) Cryo-EM structure of the Type-1 Δ*pufY* RC–LH1 dimer (periplasmic view, LH1 shown in orange cartoon) shows a two-fold symmetric dimer with 13-14 LH1 subunits per monomer (the last pair exhibits poor density). Comparison of the structures of the Type-1 Δ*pufY* RC–LH1 dimer and the wild-type Class-1 RC–LH1 dimer (LH1 shown in grey cartoon, PufY shown in purple cartoon) shows that lack of PufY induces the shift of terminal LH1 subunits LH1-12, LH1-13, and LH1-14 towards the RC. LH1 subunits follow the contours of the RC and adopt an elliptical shape in the LH1 ring of the Δ*pufY* RC–LH1 dimer. (**g**) Cryo-EM structure of the Type-2 Δ*pufY* RC–LH1 dimer (periplasmic view, LH1 shown in orange cartoon) shows an asymmetric dimer with 13-14 LH1 subunits in one monomer and a partial LH1 ring (only 7-9 LH1 subunits, the last several pairs exhibit poor density) in the other monomer. The structure shows a wide opening in the LH1 ring that lacks LH1-8– LH1-14 subunits, compared with the structure of the wild-type Class-1 RC–LH1 dimer (LH1 shown in grey cartoon, PufY shown in purple cartoon). Note the LH1-9 pair still shows density but is considerably weak, thus was not modeled in the structure.

The integration of PufY creates extra space between LH1-13/14 and the RC, compared with the RC–LH1 structures from *Rba*. *veldkampii* ^17^ and *Rhodopseudomonas* (*Rps*.) *palustris* ^13^ (Fig. 3a, Fig. S23). Both N- and C-termini of PufY at the cytoplasmic side of the RC–LH1 complex form multiple hydrogen bonds (Glu3-Glu6, Thy49-Ser52) with RC-H (Arg202) and the N-terminal residues (Arg14, Arg15) of LH1-13α and LH1-14α (Fig. 3c, Table S5). At the periplasmic side, Ala28 at the short loop of PufY is hydrogen bonded with Ser37 of LH1-13α. Moreover, a putative UQ-10 (Q_Y_) that associates with PufY and is close to the RC Q_B_ site was identified in the cryo-EM maps of the Class 2 dimer and WT monomer, whereas it was not visible in the Class-1 dimer (Fig. 3d). The head group of Q_Y_ is located near the cytoplasmic side and is sandwiched by the aromatic rings of RC-M Trp41 and PufY Phe7 via π–π interactions, forming a quinone-binding pocket (Fig. S11).

The unique location of PufY and its interactions with the terminal pairs of LH1 and the RC suggest the possible roles of PufY in forming the opening in the LH1 ring and facilitating quinone exchange. Genetic deletion of *pufY* had no notable effect on the oligomerization states of the RC– LH1 complexes (Fig. S1), suggesting that PufY, which is not involved in monomer-monomer interactions, is not essential for dimerization of RC–LH1. Compared to the WT, the Δ*pufY* strain shows a comparable growth rate but a slightly early onset of the stationary phase under both moderate light (ML, 25 μmol photons s^-1^ m^-2^) and low light (LL, 5 μmol photons s^-1^ m^-2^) (Fig. 2e). The cryo-EM structure of the Δ*pufY* RC–LH1 monomer (2.86-Å resolution) shows that the RC is encircled by an LH1 ring of 13 subunits, forming an incomplete RC–LH1 complex with an enlarged LH1 opening of 51 Å adjacent to the Q_B_ site (Fig. 3e, Figs. S24, S25). Cryo-EM analysis reveals two classes of the Δ*pufY* RC–LH1 dimers, both possessing an S-shaped LH1 ring (Fig. 3f, 3g, Figs. S26-29). The Type-1 dimer (3.08-Å resolution) contains 13-14 LH1 subunits in each monomer (Fig. 3f, the last pair exhibits poor densities). The Type-2 dimer (3.45-Å resolution) adopts an asymmetric structure, with one monomer consisting of 13-14 LH1 subunits and the other showing a widely opened LH1 barrier with only 7-9 pairs of LH1 αβ-polypeptides (the last several pairs exhibit poor densities), completely exposing the RC Q_B_ site (Fig. 3g). In the absence of PufY, LH1-12, LH1-13, and LH1-14 shift inward to the RC and follow the contour of the RC, forming a relatively closed, elliptical LH1 ring compared with the monomers of the WT Class-1 dimer (Fig. S30). These results suggest that although PufY appears not essential in maintaining the RC–LH1 functionality like PufX (at least under the tested ML and LL conditions), it stabilizes LH1 and prevents closure of the LH1 ring, thereby facilitating quinone transport to the RC Q_B_ site. It may further intimate the importance of dimerization and modularity of the RC–LH1 complex: when the access of quinones to one of the RC Q_B_ sites is obstructed, the other RC–LH1 monomer with an incomplete LH1 barrier within the dimer could ensure sufficient quinone/quinol exchange of the whole dimeric complex.

### Assembly pathway of the RC–LH1 dimer

The cryo-EM structures provide evidence for the stepwise assembly of the dimeric RC–LH1 complex (Fig. 4). PufX interacts with RC-L, RC-H and the 1^st^ LH1 αβ-heterodimer within the same RC–LH1 monomer (Fig. 2), suggesting that PufX appears at the early stage of the RC–LH1 assembly, which is supported by biochemical results ^35,45^. The association of PufX with the RC provides an anchoring point for the 1^st^ LH1 subunit, initiating the sequential assembly of the LH1 ring around the RC and stabilizing the RC–LH1 association. Incorporation of PufY then leads to recruitment and repositioning of the LH1 terminal subunits near the Q_B_ site of the RC, creating a wide channel for efficient quinone/quinol exchange between the RC and Cyt *bc*_1_.

**Fig. 4.**
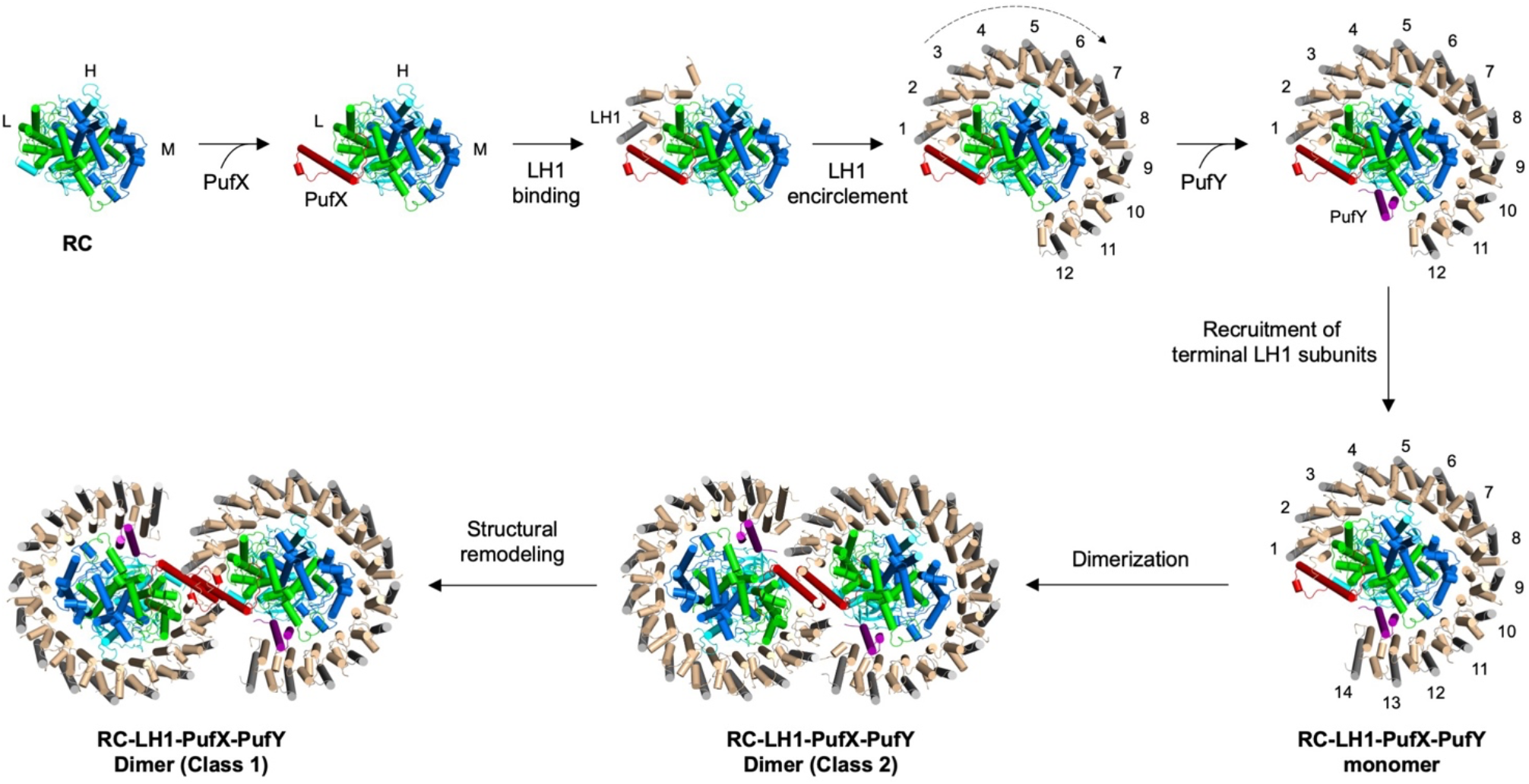
A proposed assembly pathway of the dimeric RC–LH1–PufX–PufY core complex. The sequential assembly of the RC–LH1 dimer starts from the binding of PufX to the RC complex, which provides an anchoring point for binding with the first LH1 αβ-subunits. This triggers the sequential assembly of LH1 subunits to ensure encirclement of the RC. Association of PufY prevents complete closure of the LH1 ring and creates extra space around the RC Q_B_ site to facilitate quinone/quinol binding and diffusion. This eventually generates the RC– LH1–PufX–PufY monomer. Subsequently, the interface of the monomer formed by PufX, RC-H, and LH1 subunits promotes dimerization, resulting in the formation of the Class-2 RC–LH1–PufX–PufY dimer as an intermediate structure. Extensive interactions formed at the monomer-monomer interface eventually create the stable and photosynthetically active Class-1 RC–LH1–PufX–PufY dimer.

The WT RC–LH1 monomer has a higher structural similarity with the Class-2 dimer than Class-1: The N-terminus of PufX in the WT monomer and Class-2 dimer is subject to conformational flexibility without interactions with the neighboring monomer; no interaction between PufX and RC-H is identified in the WT monomer and within the same monomer of the Class-2 dimer (Fig. 2, Figs. S8, S18). These results, together with the distinct BChl arrangements (Figs. S10, S11), suggest that the Class-2 dimer may serve as an intermediate structure in the formation of a mature RC–LH1 dimer from monomers. Hence, dimerization of the RC–LH1 core complex may occur after the formation of a complete RC–LH1–PufX–PufY monomer. Two RC–LH1–PufX–PufY monomers associate through the interactions between PufX, RC-H, and LH1-1, forming the Class-2 RC–LH1– PufX–PufY dimer as an intermediate structure. In the final step, the two monomers in the Class-2 dimer twist, driven by increasing inter-monomer interactions, leading to the eventual formation of the Class-1 RC–LH1–PufX–PufY dimer. This hierarchical assembly process signifies the necessity of a modular organization and a flexible dimerization interface of the RC–LH1 core complex. Consistent with this, some monomers and irregular structures are always seen together with dimers in chromatophores ^24,46^.

Recently, four “protein-Z” polypeptides have been identified in a dimeric RC–LH1 structure from a *Rba*. *sphaeroides* ΔLH2 mutant strain grown under semi-aerobic conditions in darkness ^47^. In our Class-1 and Class-2 dimer structures from the *Rba*. *sphaeroides* WT (DSM 158) strain that was grown phototrophically under anoxic conditions, the cryo-EM densities at the corresponding positions of “protein-Z” are too weak to determine their presence and organization (Fig. S31), albeit a high global resolution (2.74 Å for Class-1) and local resolution (Fig. S6). This discrepancy is presumably attributed to the distinctive strain growth conditions.

### Pathways for quinone/quinol exchange in the RC–LH1 dimer

In the Class-1 dimer, three UQ-10 molecules were identified in each monomer (Fig. 5, Fig. S11). Two are located at the Q_A_ and Q_B_ sites. The third one (termed Q_3_) is located near the periplasmic side and its head group projects towards the isoprenoid tail of Q_B_, suggesting its involvement in the exchange of Q_B_. In addition, an extra UQ-10 molecule (Q_Y_) was identified in the Class-2 dimer and WT monomer at the gap region between PufY and the RC (Fig. 3d, 5), suggesting that PufY might be involved in congregating quinones close to the Q_B_ site where quinones are photoreduced and protonated to quinols. The tail of Q_Y_ projects into the membrane and is adjacent to the head group of Q3 and the tail of Q_B_. The three UQ-10 molecules are clustered together, with head–head distances of ~23 Å (Q_B_–Q_3_), ~22 Å (Q_B_–Q_Y_) and ~25 Å (Q_3_–Q_Y_), delineating the routes for quinone/quinol entering or leaving the Q_B_ site. Coarse-grained molecular dynamics (CGMD) simulations indicate that Q_Y_ could move toward Q_3_, while the Q_3_ molecule shifts toward the Q_B_ site (Fig. S32), reminiscent of the “diving” of quinones in membranes ^6^. Q_Y_ was not identified in the cryo-EM structure of Class-1 dimer, likely due to its high dynamics and transient binding with PufY in the photosynthetically active RC–LH1 dimer (Fig. S32).

**Fig. 5.**
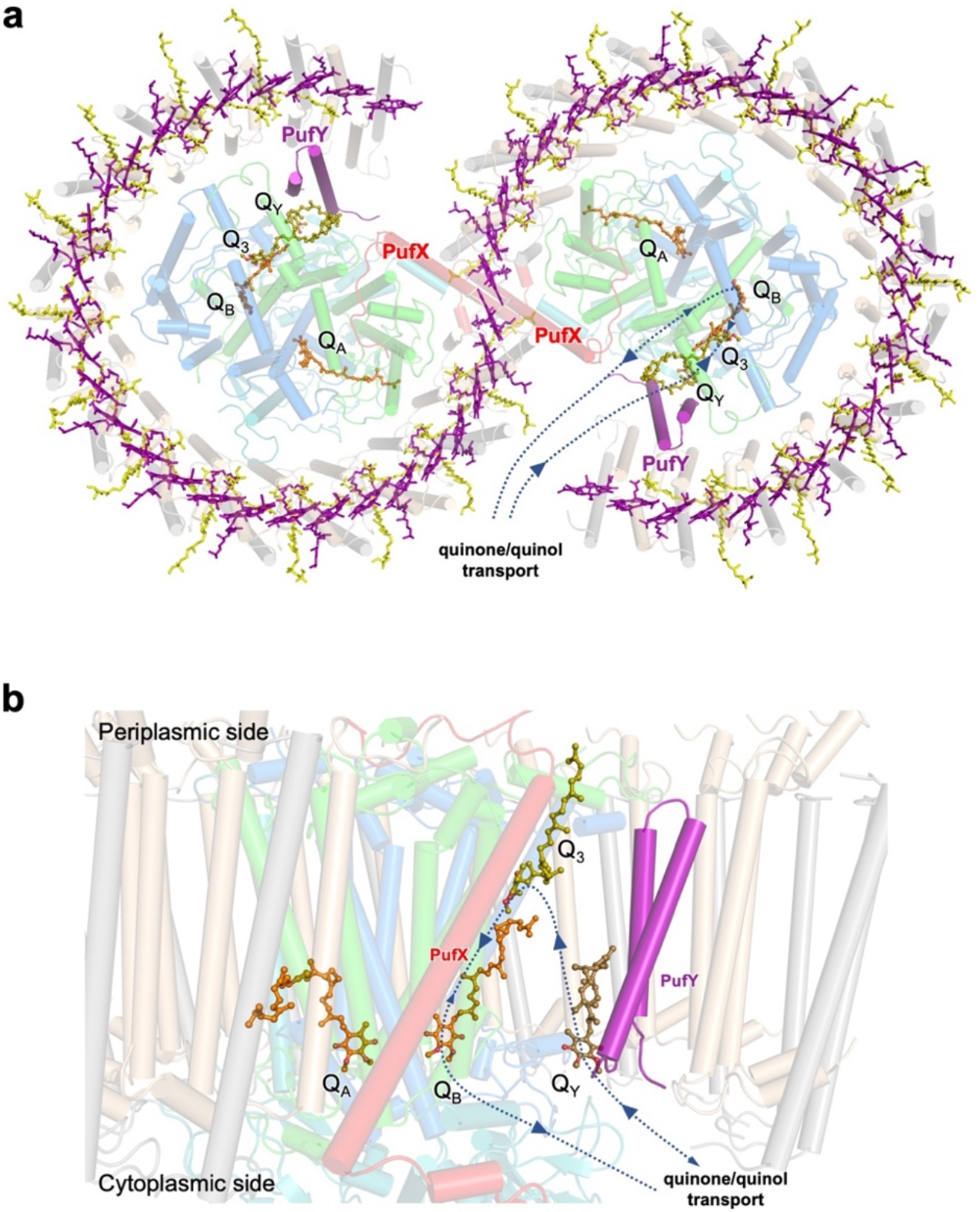
Quinone exchange pathways in the RC–LH1–PufX–PufY dimer as suggested by cryo-EM structures and computational simulations. (**a**) Periplasmic view of the locations of quinones (Q_A_, Q_B_, Q_3_, and Q_Y_) in the RC–LH1 dimer. The arrows indicate the quinone/quinol diffusion pathways through the large opening in the LH1 ring, which is formed by PufX and PufY. (**b**) Side view of the quinone transport route through the Q_Y_, Q_3_, and Q_B_ molecules. The pathways indicated by larger arrows are verified by CGMD simulations (Fig. S32).

The *Rba*. *sphaeroides* LH1 unit contains two carotenoids, distinct from other reported RC– LH1 complexes that have only one carotenoid per LH1 unit. The tightly arranged pigments within the LH1 array may block the channels between adjacent LH1 subunits for quinone/quinol passage (Fig. S15), consistent with previous results that photosynthetic growth was partially recovered when LH1 was deleted ^40^ or by decreasing the carotenoid content in the LH1 barrier ^27^. In contrast, cryo-EM structures and UQ-10-binding CGMD simulations indicate that the large openings in the S-shaped LH1 ring, formed by PufX and PufY, provide the dominant channels for quinol/quinone exchange of the RC Q_B_ site (Fig. 5, Fig. S32). The tilted conformation of PufX and its binding with the RC and LH1 subunits ensure an open LH1 ring. Removal of PufX resulted in an RC–LH117 complex with a completely closed LH1 ring, and thereby the loss of phototrophic viability (Fig. 2). Spectroscopic analysis confirmed that PufX could facilitate the quinone-mediated redox interaction between the RC and Cyt *bc*_1_ in *Rba*. *veldkampii* and *Rba*. *sphaeroides* ^48^. PufY interacts with the terminal LH1 subunits and provides steric hindrances to create additional space near the Q_B_ site. The opening adopts a similar location within the LH1 ring as the gaps with varying sizes observed in other reported RC–LH1 structures ^7–9,12–14,17^ (Fig. S23), implying the general mechanism of quinone transport to/from the Q_B_ site.

## CONCLUSIONS

Our cryo-EM structures describe in detail the hierarchical assembly of the dimeric photosynthetic RC–LH1–PufX–PufY supercomplex and the roles of PufY and PufX in mediating LH1 encirclement and dimerization of the core complex to facilitate rapid shuttle of quinones/quinols between the RC and Cyt *bc*_1_. Our results provide insights into the structural variability and modularity of bacterial photosynthetic complexes, which enable efficient light harvesting, excitation energy transfer, and quinone transport that underpin photosynthesis in phototrophic bacteria grown in changing environmental conditions.

## MATERIALS AND METHODS

### Strains, mutagenesis and growth methods

*Rba*. *sphaeroides* wild-type (DSM 158) and the Δ*pufY* mutant were grown phototrophically under anoxic conditions in liquid M22+ medium supplemented with vitamins (0.08 M nicotinic acid, 0.01 M thiamine, 7.3 mM 4-aminobenzoic acid, 0.4 mM d-biotin) and 0.1% casamino acids, at 29°C in sealed glass bottles under a light intensity of 25 μmol photons s^-1^ m^-2^ provided by Bellight 70 W halogen bulbs unless specified otherwise. Growth assays of the WT and Δ*pufY* strains were performed under low light (LL, 5 μmol photons s^-1^ m^-2^) and moderate light (ML, 25 μmol photons s^-1^ m^-2^). The non-phototrophic *Rba*. *sphaeroides* Δ*pufX* mutant was grown at 29°C in the dark in the same medium under microoxic conditions in an orbital shaker set at 150 rpm.

Unmarked genomic deletions were constructed in WT *Rba. sphaeroides* using the allelic exchange suicide vector pk18mobsacB using the previously described method ^49^. Deletion of *pufX (rsp_0255*) was conducted as follows: primer pairs XupF and XupR (AGTCTCTAGAGCACCTATCTCCGCGCTCAG and CTGCCCCGAGACTTGTCTCAGTGTGATCGCTCCTCAGTTCAG) and XdownF and XdownR (CTGAACTGAGGAGCGATCACACTGAGACAAGTCTCGGGGCAG and ATGCAAGCTTGTCGTAGGCGGATTCCGAGC) were used to amplify the regions flanking *rsp_0255*, fused via PCR, digested and cloned into the BamHI and HindIII sites of pk18mobsacB. For unknown reasons, the regions flanking *pufY* (*rsp_7571*) could not be fused by the same method, so a 1-kbp synthesized DNA fragment (Genewiz, Germany), comprised of two 500-bp regions identical to the upstream and downstream DNA sequences flanking *rsp_7571*, was digested and cloned into the same sites of the pK18mobsacB vector as detailed above. The resulting plasmids were transferred from *Escherichia (E.) coli* S17 cells to *Rba. sphaeroides* by conjugation. Selection of transconjugants was performed on M22 agar containing 30 μg·mL^-1^ kanamycin, and second recombinants were isolated on M22 medium containing 10% (w/v) sucrose. Successful generation of Δ*pufX* and *ΔpufY* strains was confirmed using PCR using Q5 High-Fidelity DNA Polymerase (New England Biolabs, UK) and DNA sequencing (Eurofins).

### Purification of RC-LH1 complexes

The cells were harvested by centrifugation at 5,000 g for 10 min at 4°C, washed twice with Tris-HCl (pH 8) and resuspended in working buffer (20 mM HEPES-Na, pH 8.0). Cells were disrupted by passage through a French press three times at 16,000 psi. Cell debris was removed by centrifugation at 20,000 g for 30 min. Membranes were collected by centrifuging the resulting supernatant at 125,000 g for 90 min and were solubilized by addition of β-DDM (n-dodecyl β-D-maltoside) to a final concentration of 3% (w/v) for 15 min to 1 hour in the dark at 4°C with gentle stirring. After the unsolubilized material was removed by centrifugation at 21,000 g for 30 min, the clarified supernatant containing solubilized photosynthetic complexes was applied onto the 10–30% or 10–25% (w/v) continuous sucrose gradients made with working buffer containing 0.01% (w/v) β-DDM. Gradients were centrifuged at 230,000 g for 18 h. For the WT dimers, a milder detergent α-DDM (n-dodecyl α-D-maltoside) was used in the isolation of RC-LH1 complexes, which includes a further purification step using a Superose 6 gel filtration column (GE). The purity of the RC-LH1 complex was characterized by SDS-polyacrylamide gel electrophoresis (SDS-PAGE). Bands of proteins smaller than 15 kD were cut and analyzed by LC-MS/MS as described below, to identify the new peptide PufY. For the WT monomers and mutants, the RC-LH1 complexes were collected and concentrated using Vivaspin 6 100,000 MWCO columns (Cytiva) for heavy complexes (dimers) and Vivaspin 6 50,000 MWCO columns for light complexes (monomers). Simultaneously, the buffer containing sucrose was exchanged to working buffer containing 0.01% (w/v) DDM.

### Cryo-EM data collection

A 3.0-μL aliquot of the RC-LH1 complex was applied onto the holey carbon grid (300M-Au-R1.2/1.3; Beijing EBO technology limited) glow-discharged for 45 s for WT dimers, or the copper grids (Quantifoil R1.2/1.3 Cu, 300 mesh) with a thin carbon-supported film glow-discharged for 70 s for the WT monomers, Δ*pufX* monomers, and Δ*pufY* RC-LH1. The grids were plunge-frozen into liquid ethane using a Vitrobot Mark IV (Thermo Fisher Scientific). Parameters for plunge freezing were set as follows: blotting time of 3 s (WT monomer, Δ*pufY* monomer, Δ*pufY* dimers), 4 s (WT dimers) or 6 s (Δ*pufX* monomers), waiting time of 30 s (WT dimers, WT monomer, Δ*pufY* monomer, Δ*pufY* dimer) or 3 s (Δ*pufX* monomer), blotting force level of 0 (Δ*pufY* monomer, Δ*pufY* dimers), 4 (WT dimers) or −10 (WT monomer, Δ*pufX* monomer), humidity 100%, chamber temperature 4°C.

For the *Rba. sphaeroides* WT dimers, the micrographs were collected at FACILITY on a 300 kV Titan Krios microscope (Thermo Fisher Scientific) equipped with a K2 Summit direct electron detector (Gatan) at the Center for Biological Imaging, Core Facilities for Protein Science at the Institute of Biophysics, Chinese Academy of Sciences. Automated data acquisition was facilitated at a 130,000 magnification in super-resolution counting mode by SerialEM software. Images were recorded by beam-image shift data collection methods ^50^. Each raw movie stack (32 frames) was captured using an exposure time of 6.4 s leading to a total dose of 60 e^-^/Å^2^ and a defocus value in the range from −1.8 to −2.2 μm. A total of 6,227 images were binned resulting in a pixel size of 1.04 Å for further data processing. Other RC-LH1 complexes were collected at the cryo-EM facility of RIKEN Center for Biosystems Dynamics Research (Yokohama) on a 300 kV Titan Krios equipped with a K3 Summit direct electron detector (Gatan) in counting mode, using EPU software. For the WT monomer, a total of 4,700 movies were recorded at an 81,000 magnification and a pixel size of 1.06 Å. Each raw movie stack (48 frames) was captured by applying a total dose of 45.026 e^-^/Å^2^. The defocus ranged from −0.7 to −1.8 μm. For the Δ*pufX* monomer, a total of 4,807 movies were recorded at an 81,000 magnification and a pixel size of 1.06 Å. Each raw movie stack (48 frames) was captured by applying a total dose of 46.549 e^-^/Å^2^. The defocus ranged from −0.8 to −1.8 μm. For the Δ*pufY* monomer, a total of 4,700 movies were recorded at a 105,000 magnification and a pixel size of 0.8285 Å. Each raw movie stack (48 frames) was captured by applying a total dose of 51.527 e^-^/Å^2^. The defocus ranged from −0.8 to −2 μm. For the Δ*pufY* dimer, a total of 12,209 movies were recorded at a 105,000 magnification and a pixel size of 0.8285 Å. Each raw movie stack (48 frames) was captured by applying a total dose of 50.868 e^-^/Å^2^. The defocus ranged from −0.8 to −2 μm.

### Data processing

Collected movies were imported into RELION 3.0 or 3.1 ^51,52^ and motion corrected using MotionCor2 ^53^. Contrast transfer function (CTF) parameters were determined by CTFFIND-4.1 ^54^ or Gctf ^55^. For the WT dimers, a subset of particles was manually picked and processed with reference-free 2D classification, and the five 2D-class averages were selected as references for further automatic particle picking of the complete dataset, resulting in a total of 957,176 particles. 765,091 particles were selected for 3D classification after two rounds of reference-free 2D classification, which were classified into four classes using the initial model from the 8-Å RC–LH1 complex of *Rba. sphaeroides* (PDB ID: 4V9G), which was lowpass filtered to a lower resolution of 20 Å. A total of 434,588 particles were kept for further 3D classification with C2 symmetry. Two distinct classes (145,392 particles for Class-1 and 115,388 particles for Class-2) were selected for 3D auto-refinement, CTF refinement and Bayesian polishing, which resulted in the 2.74-Å and 2.96-Å density maps for Class-1 and Class-2, respectively.

For the WT monomer, Δ*pufX* monomer, Δ*pufY* monomer and Δ*pufY* dimer datasets, particles were picked automatically using SPHIRE-crYOLO ^56^. The input box sizes were 250 × 250 pixels for WT monomers and Δ*pufX* monomers, 300 × 300 pixels for Δ*pufY* monomers, and 400 × 400 pixels for Δ*pufY* dimers. The total numbers of particles picked were 241,140 for WT monomers, 809,777 for Δ*pufX* monomers, 241,170 for Δ*pufY* monomers, and 373,025 for Δ*pufY* dimers. Particles were then extracted in RELION 3.1 and subjected to reference-free 2D classifications. After each classification step, only the particles sorted into well-defined classes were selected to continue being processed. A 3D initial model was calculated in RELION 3.1 based on those particles and used as a reference for subsequent 3D classifications of selected particles. Once a well-defined 3D class was obtained, it was refined into a high-resolution electron potential map. The final selected particle numbers were 68,554 for WT monomers, 80,701 for Δ*pufX* monomers, 56,391 for Δ*pufY* monomers, 71,027 for Δ*pufY* Type-1 dimers and 53,830 for Δ*pufY* Type-2 dimers. Each map then went through several iterations of CTF parameter refinement, particle polishing, and 3D auto-refinement before it was corrected for the modulation transfer function (MTF) of the Gatan K3 summit camera and sharpened by RELION’s post-processing job type. Based on the Fourier shell correlation (FSC) 0.143 criterion, the final global map resolutions were estimated to be 2.79 Å for WT monomers, 6.62 Å for Δ*pufX* monomers, 2.86 Å for Δ*pufY* monomers, 3.08 Å for Δ*pufY* Type-1 dimers, and 3.45 Å for Δ*pufY* Type-2 dimers. To improve the resolution of the Δ*pufX* monomer, particles that made up the final map were further used to train Topaz ^57^ to recognize particles of interest, and particle picking was repeated using the trained Topaz. A total of 3,420,815 particles were picked, classified into reference-free 2D classes and subsequently into 3D classes until a well resolved 3D class was obtained. The final number of particles was 66,058 and the final map resolution was 4.20 Å. Local map resolutions were calculated by RELION 3.1.

### Model building and refinement

Two copies of 1.9-Å crystal structures of RC-LH1 from *Tch. tepidum* (PDB ID: 5Y5S) were initially docked into the 2.74-Å resolution cryo-EM map of the WT Class-1 dimer using UCSF Chimera ^58^. The amino acid sequences were further mutated to its counterparts in *Rba. sphaeroides* 2.4.1. PufY was identified as the protein RSP_7571 through mass spectrometry, which was further confirmed based on the perfect match of the specific sequence with the cryo-EM densities. The model of Class-1 was rebuilt manually basing on the cryo-EM density with COOT ^59^ and then real-space refined using Phenix 1.19.2 ^60^ using C2 symmetry. The model of Class-2 was rebuilt and refined following the same procedure using the structure of Class-1 as the initial model. MolProbity ^61^ was used to evaluate the geometries of the structures. The atomic model of the WT monomer was built using the WT Class-2 dimer as a reference starting model. All amino acid sequences making up the models of WT monomer and dimers are listed in Fig. S33. The orientation of the SPO-β molecules in the RC-LH1 complexes was tentatively assigned based on the resolutions of the WT monomer and Class-1 dimer structures. The models of Δ*pufX* and Δ*pufY* monomers were built using the WT monomer as a starting model and both Δ*pufY* dimers were built using the WT Class-1 dimer as a starting model. The starting model structures were fit into experimental electron density maps as rigid bodies using UCSF Chimera. Three LH1 subunits were added to the starting structure of the Δ*pufX* monomer and fitted into map, and PufX was removed. PufY was removed in all the Δ*pufY* structures, and several LH1 subunits were removed in the Δ*pufY* monomer and Type-2 dimer. These modifications were performed by using UCSF Chimera. Each structure was then adjusted manually using COOT. The final model was refined by Phenix 1.19.2, and the stereochemistry was assessed by MolProbity. Images were prepared with Chimera, ChimeraX ^62^ and PyMOL ^63^ (Molecular Graphics System).

### Growth assays

Three replicates of *Rba*. *sphaeroides* cultures were grown phototrophically in 8 ml glass tubes as detailed above. The tubes were constantly rotated using the RM-4D Rotator mixer (Premiere). Optical density at 680 nm was measured using a Colorimeter (WPA Colourwave CO7000).

### Absorption spectra

Purified RC-LH1 complexes from sucrose gradient was collected and absorbance was measured from 300 to 900 nm in 1-nm intervals using a Libra S22 spectrophotometer (Biochrom).

### Analysis of pigments and quinones

The quinones from the dimeric RC–LH1 samples were extracted using 1:1 methanol:chloroform (v:v) containing 0.02% (w/v) FeCl3, and were injected into a Shimadzu 20A HPLC system employing a Shim-pack GIST C-18 reversed-phase column (4.6 mm × 250 mm). The column was pre-equilibrated and was then eluted with 8:2 methanol:isopropanol (v:v) at a flow rate of 1 mL min^-1^ for 1 hour at 40°C. The elution was analyzed by an LC-20AT detector (Shimadzu), monitored at 275 nm. Ubiquinone-10 (UQ-10) was identified by comparing the retention time of the quinones from the RC-LH1 complexes with that of the UQ-10 standard (TargetMol, China, Cat No.: T2796).

### Liquid chromatography-mass spectrometry (LC-MS/MS) protein identification

Gel bands containing the protein sample were manually excised. Each of the protein bands was then digested individually as below. The protein bands were cut into small plugs, washed twice in distilled water, and were then dehydrated in 100% acetonitrile for 10 min and dried in a SpeedVac (Labconco) for 15 min. Reduction (10 mM DTT in 25 mM NH_4_HCO_3_ for 45 min at 56°C) and alkylation (40 mM iodoacetamide in 25 mM NH_4_HCO_3_ for 45 min at room temperature in the dark) were performed, followed by washing with 50% acetonitrile in 25 mM NH_4_HCO_3_ twice. The gel plugs were then dried using a SpeedVac and digested with sequence-grade modified trypsin (40 ng for each band) in 25 mM NH4HCO3 overnight at 37°C. The enzymatic reaction was stopped by adding formic acid to a 1% final concentration. The solution was transferred for LC-MS/MS analysis.

LC-MS/MS analysis was performed using a nanoLC-LTQ-Orbitrap XL mass spectrometer (Thermo Scientific, San Jose, CA) in line with an Easy-nLC 1200 HPLC system. Tryptic peptides generated above were loaded onto a self-packed trap column (ReproSil-Pur C18-AQ, 150 μm i.d. × 2 mm, 5 μm particle) (Dr. Maisch GmbH, Ammerbuch) which was connected to the self-packed analytical column (ReproSil-Pur C18-AQ, 75 μm i.d × 150 mm, 3 μm particle). The peptides were then eluted over a gradient (0-36% B in 78 mins, 36-80% B in 12 mins, where A = H_2_O with 0.1% formic acid and B = 80% acetonitrile with 0.1% formic acid) at a flow rate of 300 nL min^-1^ and introduced online into the linear ion trap mass spectrometer using nano electrospray ionization (ESI). Data-dependent scanning was incorporated to select the 10 most abundant ions (one microscan per spectra; precursor isolation width ± 1.0 m/z, 35% collision energy, 30 ms ion activation, exclusion duration: 120 s; repeat count: 1) from a full-scan mass spectrum (300 to 1600 m/z at res = 60,000) for fragmentation by collision induced dissociation (CID). Lock mass option was enabled for the 462.14658 m/z.

MS data were analyzed using Proteome Discoverer (version 1.4.0.288, Thermo Scientific). MS2 spectra were searched against the UniProt proteome database of *Rhodobacter sphaeroides* 2.4.1 (UniProt ID: UP000002703) using the SEQUEST search engine. Peptides with and above +2 charge states were accepted if they were fully enzymatic. The following residue modifications were allowed in the search: carbamidomethylation on cysteine and oxidation on methionine. Peptide tolerance of 20 ppm and fragment mass tolerance of 0.6 Da were applied. Peptide spectral matches (PSM) were validated by a targeted decoy database search at a 1% false discovery rate (FDR). Peptide identifications were grouped into proteins according to the law of parsimony.

### All-atom molecular dynamics (AAMD) simulations

The dimeric structure was divided into two monomers and the position of the lipid bilayer of each monomer was predicted by the Positioning of Proteins in Membrane (PPM) server ^64^. The monomer structures were superposed on the dimeric structure to calculate the positions of the lipid bilayers in the dimer. The average vector of the normal vectors of the two lipid-bilayer surfaces was calculated. The dimeric structure was translated and rotated, so that the geometric center was at the origin and the average vector was aligned to the Z axis. The dimeric structure was then embedded in a solvated lipid bilayer using the “Membrane Builder” function ^65^ of the CHARMM-GUI server ^66^. As some atoms of BChls *a*, BPhes and UQ-10 are missing in the cryo-EM structure, their structures were modeled manually. The generated system was composed of 66 protein chains, 64 BChls *a*, 4 BPhes, 2 tetrastearoyl cardiolipin (TSCL), 52 SPOs, 8 UQ-10, 16 1,2-distearoyl-*sn*-glycero-3-phosphocholine (DSPC) and 1,389 1-palmitoyl-2-oleoyl-*sn*-glycero-3-phosphocholine (POPC) molecules, as well as 2 Fe^2+^, 545 K^+^, 576 Cl^-^ ions, and 196,050 water molecules. The total number of atoms was 868,794 and the size of the initial system was 25.6 nm × 25.6 nm × 14.1 nm. The same topologies and parameters as those used in the previous study ^17^ were used for BChls *a*, BPhes, and SPOs. The topology and the parameters of TSCL were created by modifying those of tetraoleoyl cardiolipin provided by the CHARMM-GUI server. Distances between the metal ions (Mg^2+^ and Fe^2+^) and their coordinating atoms were restrained with a force constant of 1.0 × 10^5^ kJ nm^-2^. The TIP3P model ^67^ was used for the water molecules. The CHARMM36m force field ^6^ was used for the protein chains and the CHARMM36 force field ^68–70^ was used for the other molecules.

After the system was energy-minimized, the system was equilibrated in nine steps. In the first two steps, the system was equilibrated for 0.25 ns in the *NVT* ensemble, and in successive three steps, it was equilibrated for 1.125 ns in the *NPT* ensembles. In these steps, the positions of the protein and the ligand non-hydrogen atoms were restrained to their initial positions. The force constants were gradually decreased from 4,000 to 1,000 kJ nm^-2^ for the protein backbone and the ligand atoms and from 2,000 to 500 kJ nm^-2^ for the protein sidechain atoms. The Z coordinates of the phosphorus atoms of the POPC molecules were also restrained with the force constants from 1,000 to 40 kJ nm^-2^. In the sixth step, a 40-ns MD simulation was performed with the position restraints of 1000 kJ nm^-2^ for the protein backbone and the ligand atoms and of 500 kJ mol^-2^ for the protein sidechain atoms. No restraints were imposed on the lipid atoms. In the last three steps, an MD simulation was performed for 1.5 ns in total. The force constants were gradually decreased to 50 kJ nm^-2^ for the protein backbone and the ligand atoms and 0 kJ nm^-2^ for the protein sidechain atoms. Finally, a 500-ns production MD simulation was performed without restraints. The temperature was kept at 303.15 K throughout the MD simulations and the pressure was kept at 1.0 × 10^5^ Pa except for the first two equilibration steps. In the equilibration steps, the Berendsen weak coupling method ^71^ was used to control the temperature and the pressure. In the production run, the Nosé-Hoover method ^72,73^ and the Parrinello-Rahman method ^74,75^ were used to control them, respectively. Bond lengths involving hydrogen atoms were constrained using the LINCS algorithm ^76,77^ to allow the use of a large time step (2 fs). Electrostatic interactions were calculated with the particle mesh Ewald method ^78,79^. All AAMD simulations were performed using GROMACS 2021 ^80^, with coordinates recorded every 10 ps.

### Coarse-grained molecular dynamics (CGMD) simulations

CGMD simulations were performed for the whole dimer structure and the UQ-10-free structure. The former is referred as to “UQ-10-bound simulation”. In the latter simulation, all the UQ-10 molecules were removed from the dimer structure and 60 UQ-10 molecules were randomly placed around the dimer as described below to investigate the UQ-10-binding process. Therefore, the latter simulation is referred as to “UQ-10-binding simulation”. The coarse-grained (CG) models were constructed using the MARTINI 2.2 force field ^81^. The CG parameters of UQ-10, DSPC, and POPC were taken from literatures ^82,83^. The parameters of the coarse-grained models of BChls, BPhes, and TSCL were created by modifying those of chlorophyll *a* ^82^, pheophytin *a* ^82^, and tetraoleoyl cardiolipin ^84^, respectively (for detail, see Supplementary Text). The parameters of SPO were determined so that the CG model reproduced the dynamics observed in AAMD and the water-octanol partition coefficient predicted by the XLOGP3 program ^85^. The ElNeDyn model ^86^ was used to construct the CG models of protein chains and to maintain the quaternary structure of the dimer with an elastic network. All the backbone beads of the protein, the SP3 beads of BChls and BPhes and all the beads of SPOs were mutually connected by springs when the distance between a bead pair was within 9 Å. The force constants were 500 kJ mol^-1^ nm^-2^ for the backbone bead pairs and 100 kJ mol^-1^ nm^-2^ for the other pairs. In addition, the side chain beads of proteins within 3.5 Å from the Mg^2+^ ions of BChls were connected with the Mg^2+^ ions by springs with a force constant of 500 kJ mol^-1^ nm^-2^ to maintain the coordination structure. Each of the whole (i.e., UQ-10-bound) and the UQ-10-free structures was embedded in a solvated lipid bilayer using the CHARMM-GUI server ^66,87^. Each solvated lipid bilayer was composed of approximately 2,000 POPC molecules, 58,000 CG waters, and 0.15 M Na^+^/Cl^-^ ions that neutralized the net charge of the system. After energy minimization, each system was equilibrated for 1 μs in the *NPT* ensemble. During the equilibration, the positional restraints were imposed on the protein backbone beads (for both the systems) and the UQ-10 beads (for the system with the UQ-10-bound structure). For the UQ-10-binding simulation, 60 POPC molecules in the system with the UQ-10-free structure farther than 140 Å from the center of mass of the dimer were randomly replaced by UQ-10 molecules. The system was energy-minimized and equilibrated for 100 ps with the positional restraints imposed on the protein backbone beads. For each system, a production run was performed for 5 μs in the *NPT* ensembles. The temperature was kept at 303.15 K using the velocity-rescaling method ^88^ with coupling constants of 1.0 ps. The pressure was kept at 1.0×10^5^ Pa using the Parrinello-Rahman method ^74,75^ with coupling constants of 12.0 ps. Electrostatic interactions were calculated using the reaction-field method ^89^ with a cutoff of 1.1 nm. Van der Waals interactions were calculated with a modified Lennard-Jones potential, where the potential was shifted to zero at the cut-off distance of 1.1 nm. Bond lengths specified in the MARTINI force field were constrained using the LINCS algorithm. The time step was 20 fs. All the CGMD simulations were performed using GROMACS 2021.

## ACKNOWLEDGEMENTS

We thank B. L. Zhu, X. J. Huang, L. H. Chen, X. J. Li, G. Ji, D. Y. Fan, B. X. Huangpu, T. X. Niu, F. Sun, and other staff members at the Center for Biological Imaging (IBP, CAS) for the technical support in EM data collection; X. Ding, Z. S. Xie, L. L. Niu, N. L. Zhu, M. M. Zhang and F. Q. Yang for mass spectrometer; X. D. Su, L. Si, X. L. Zhao, H. M. Zhang and L.F. Shi for the assistance in sample preparation and data collection. Cryo-EM data of native dimers were collected at the Center for Biological Imaging, Core Facilities for Protein Science at the Institute of Biophysics (IBP), Chinese Academy of Sciences (CAS). Cryo-EM of the other complexes were collected at the cryo-EM facility of the RIKEN Center for Biosystems Dynamics Research (Yokohama). This work was supported by the Strategic Priority Research Program of CAS (XDB27020106 to M.L. and XDB37020101 to P.C.), the Royal Society University Research Fellowship (URF\R\180030 to L.-N.L.), the Royal Society International Exchanges grant (IEC\NSFC\191600) and the National Natural Science Foundation of China (32011530168) to L.-N.L. and M.L., the Royal Society Enhancement Awards (RGF\EA\181061 and RGF\EA\180233 to L.-N.L.), the Biotechnology and Biological Sciences Research Council Grant (BB/R003890/1 and BB/V009729/1 to L.-N.L.). The authors further acknowledge the support from the National Natural Science Foundation of China (32070259 to P.C., 31930064 to M.L., 32070109 to L.-N.L.). L.B. was supported by a Liverpool-Riken International PhD Studentship. This work was partially supported by Platform Project for Supporting Drug Discovery and Life Science Research [Basis for Supporting Innovative Drug Discovery and Life Science Research (BINDS)] from AMED under grant numbers JP20am0101082 (to M.S.) and JP20am0101107 (to T.T.), and by JSPS/MEXT KAKENHI (JP19H03162 to A.Y.), RIKEN Pioneering Projects “Biology of Intracellular Environments” (to M.S.) and RIKEN BDR Structural Cell Biology Project (to M.S.).

## AUTHOR CONTRIBUTIONS

M.S., M.L. and L.-N.L. conceived the study. T.T., D.P.C., M.L. and L.-N.L. designed the experiments. P.C., L.B., A.Y., B.M.C., T.N., B.Z., T.T. and D.P.C. performed the experiments. P.C., L.B., A.Y., B.M.C., T.N., B.Z., T.T., D.P.C., M.L., and L.-N.L. analyzed the results. P.C., L.B., A.Y., M.L., and L.-N.L. generated structural models. P.C., L.B., M.L., and L.-N.L. wrote the paper with the contributions from all other authors.

## CONFLICT OF INTERESTS

The authors declare no competing interests.

## Notes

### Competing Interest Statement

The authors have declared no competing interest.

