## Supplementary Information for "Structural basis for the assembly and electron transport mechanisms of the dimeric photosynthetic RC–LH1 supercomplex"

### SUPPLEMENTARY TEXT

**Parameters of the coarse-grained (CG) models of the ligands:** The parameters of the coarse-grained (CG) models of bacteriochlorophyll *a* (BCL), bacteriopheophytin *a* (BPH), and tetrastearoyl cardiolipin (TSCL) were created by modifying those of chlorophyll A (CLA) [1], pheophytin A (PHA) [1], and tetraoleoyl cardiolipin (TOCL) [2], respectively. BCL and BPH have an acetyl group in place of the vinyl group of CLA and PHA. Therefore, the SP3 beads of CLA and PHA that represent the vinyl group were replaced by Na beads. The parameters of TSCL were created by replacing the five beads representing the unsaturated aliphatic tails of TOCL with the four beads representing saturated aliphatic tails, according to the parameters of other lipids [3]. The atoms of spheroidene (SPO) were mapped to the CG beads as shown in Fig. S34. The bonded parameters of SPO were determined based on the comparison of the results of coarse-grained molecular dynamics (CGMD) simulations with those of all-atom molecular dynamics (AAMD) simulations performed for a SPO molecule in water. The system for the AAMD simulation was composed of an SPO molecule and 6,988 TIP3P water molecules. The OPLS-AA force field [4] was used for the all-atom (AA) model of SPO and its parameters were determined using the LigParGen server [4,5]. The AAMD simulations were performed for 50 ns with the following methods. The temperature was kept at 303.15 K using the Nose-Hoover method [6,7] with coupling constants of 1.0 ps. The pressure was kept at $1.0 \times 10^5$ Pa using the Parrinello-Rahman method [8] with coupling constants of 5.0 ps. Electrostatic interactions were calculated using the reaction-field method [9] with a cutoff of 1.4 nm. Van der Waals interactions were calculated with a modified Lennard-Jones potential that was shifted so that the potential value is zero at the cut-off distance of 1.4 nm. Bond lengths involving hydrogen atoms were constrained using the linear constraint solver (LINCS) algorithm [10,11] to allow a time step of 2 fs. The system for the CGMD simulation was composed of an SPO CG model and 1,900 CG water models. The CGMD simulation was performed for 5 ns with the same methods as described in the Methods section of the text. We confirmed that the distributions of the bond lengths, the angles, and the dihedrals of the CG model of SPO were agreed well with those of the corresponding values calculated from the AAMD simulation (Fig. S34). Validity of the mapping of the CG bead types was confirmed by calculating water-octanol partition coefficient based on the CGMD simulations. The SPO-water system was composed of an SPO CG model and 1,700 CG water models. The SPO-octanol system was composed of an SPO CG model, 50 CG water models, and 760 1-octanol CG models. For each system, the CGMD simulations were performed for 2 ns each in 21 states with the λ values of 0, 0.05, 0.1, 0.15, 0.2, 0.25, 0.3, 0.35, 0.4, 0.45, 0.5, 0.55, 0.6, 0.65, 0.7, 0.75, 0.8, 0.85, 0.9, 0.95, and 1. Free-energy differences between the neighboring states were calculated using the Bennett acceptance ratio method [12,13]. Solvation free energies in water ($\Delta G_{\mathrm{wat}}$) and in 1-octanol ($\Delta G_{\mathrm{oct}}$) were calculated as the sum of the free-energy differences obtained from the CGMD simulations performed for the SPO-water and the SPO-octanol systems, respectively. The calculated $\Delta G_{\mathrm{wat}}$ and $\Delta G_{\mathrm{oct}}$ values were $-13.08$ kJ mol$^{-1}$ and $-97.26$ kJ mol$^{-1}$, respectively. The partition coefficient calculated from these values was 14.51, which was well agreed with the value predicted by XLOGP3 (14.7) [14].

### SUPPLEMENTARY FIGURES

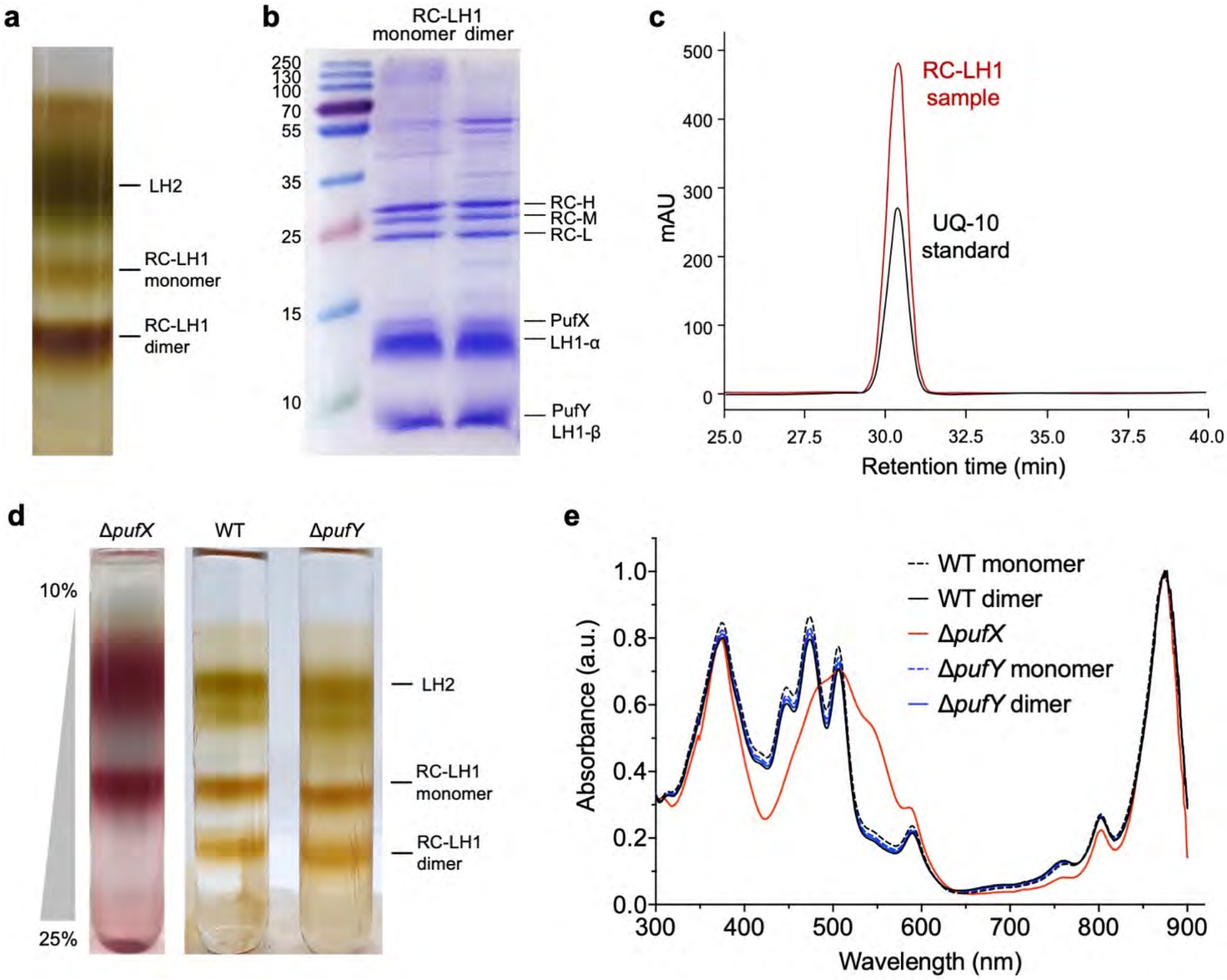


**Fig. S1. Isolation and characterization of the *Rba. sphaeroides* RC−LH1 core complexes.** (**a**) Separation of membrane proteins from WT *Rba. sphaeroides* using a 10-25 % continuous sucrose gradient. The pigmented fractions were identified as LH2, RC–LH1 monomer and RC–LH1 dimer from the top to the bottom. (**b**) SDS-PAGE of purified RC–LH1 monomers and dimers, stained by Coomassie Brilliant Blue. (**c**) HPLC analysis of quinones in the RC-LH dimers (red) using UQ-10 as a standard (black), monitored at 275 nm. (**d**) Separation of photosynthetic proteins from the WT, Δ*pufX*, and Δ*pufY Rba. sphaeroides* using 10-25 % continuous sucrose gradients. (**e**) Absorbance spectra of purified WT and mutant *Rba. sphaeroides* RC–LH1 complexes. Note that the Δ*pufX* strain was grown microoxically in the dark, given its inability to photosynthesize. The Δ*pufX* RC–LH1 complex has a broad absorption and maxima at 481, 509 and 545 nm, and thus a visibly different color, because of the conversion of the carotenoid spheroidene to spheroidenone under the microoxic culture conditions.

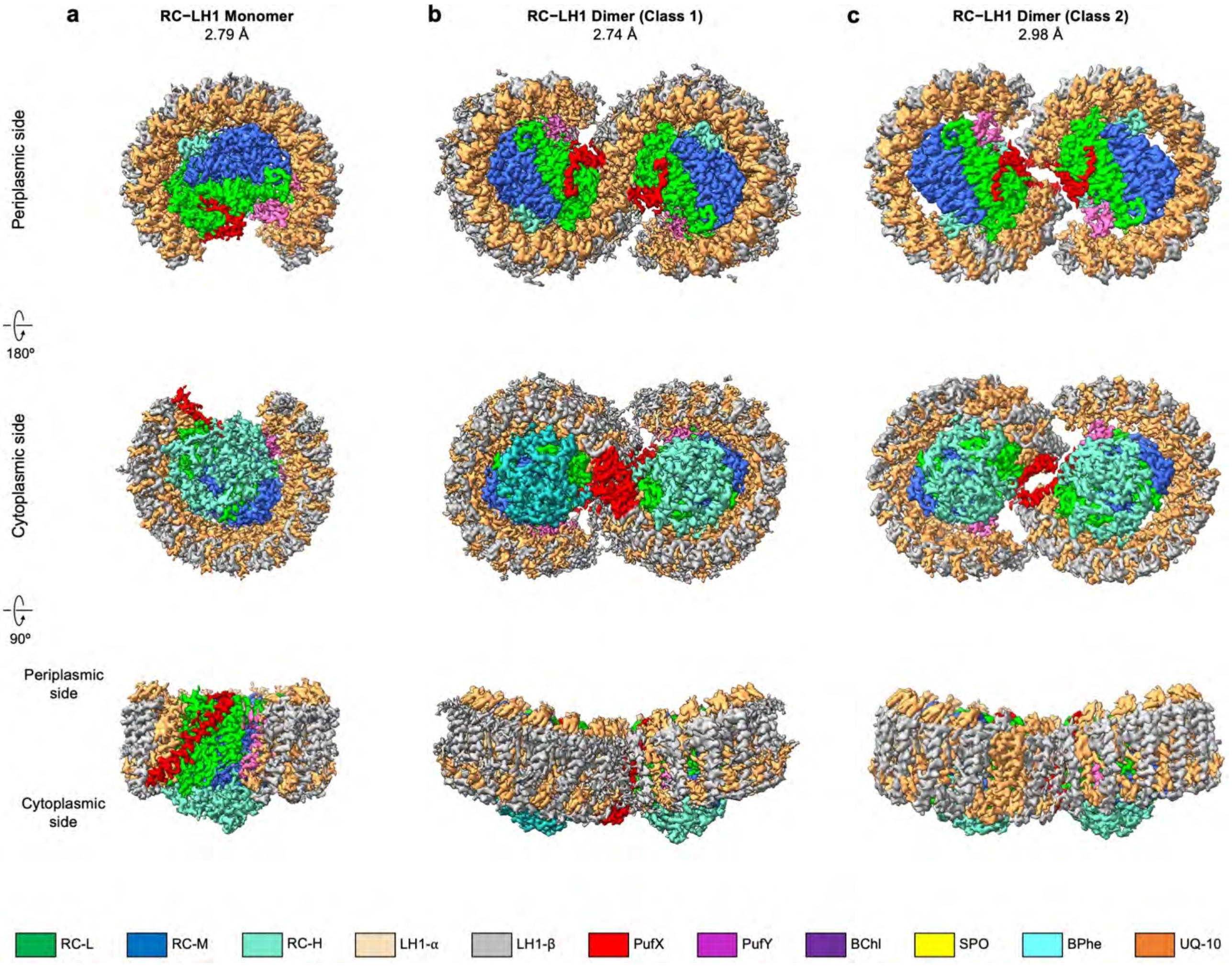


**Fig. S2. Color-coded electron density maps of the RC–LH1 core complexes from *Rba. sphaeroides*.** (**a**) The RC–LH1 monomer. (**b**) The Class-1 RC–LH1 dimer. (**c**) The Class-2 RC–LH1 dimer. Color scheme is presented as the same as shown in Fig. 1: LH1-α, wheat; LH1-β, grey; PufX, red; PufY, purple; RC-L, green; RC-M, marine; RC-H, teal; BChls, purple; BPhes, cyan; carotenoids, yellow; quinones, orange.

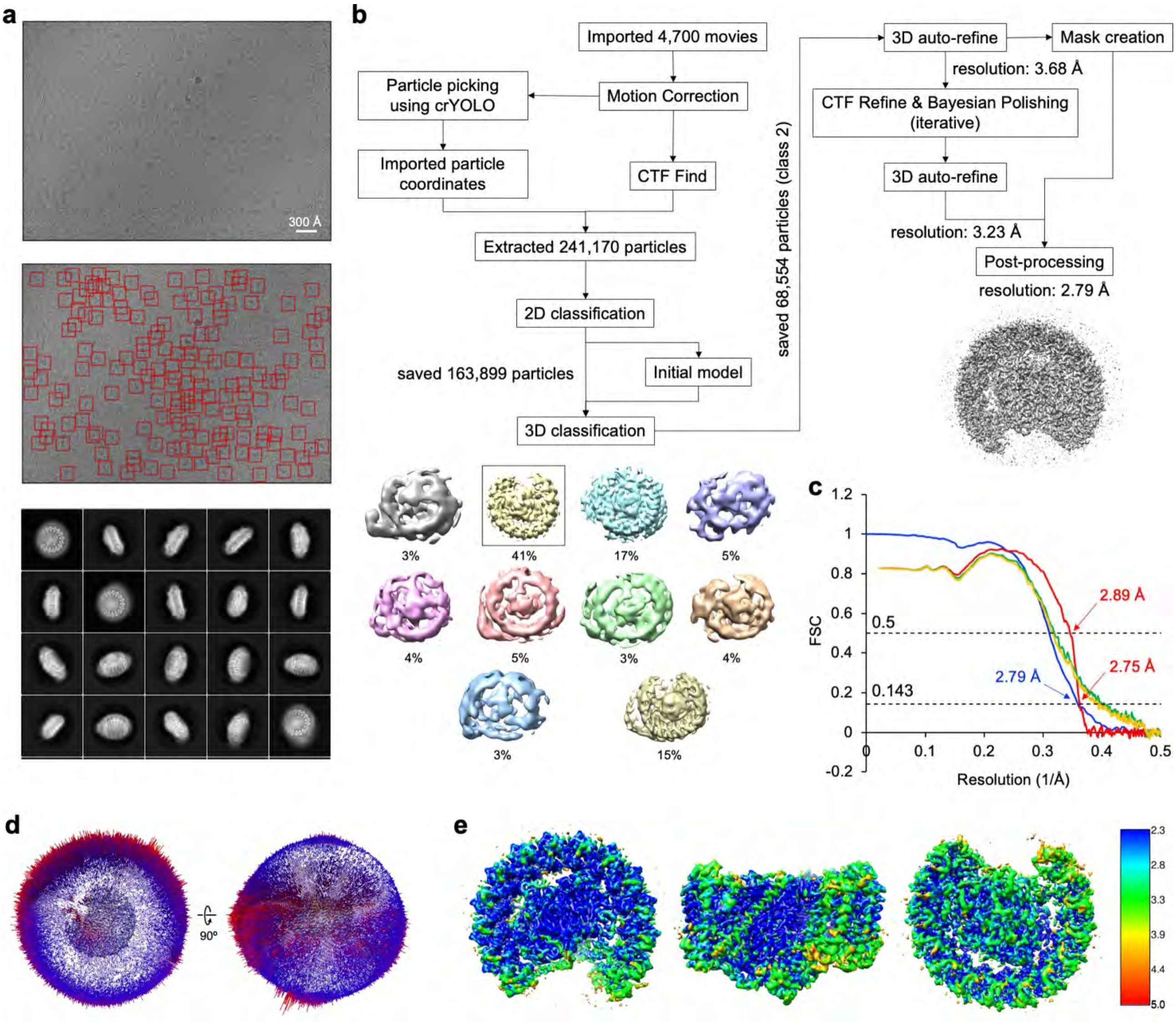


**Fig. S3. Cryo-EM data process of the *Rba. sphaeroides* WT RC–LH1 monomer.** (**a**) Motion-corrected example of a cryo-EM captured movie is shown on top and all particles picked from that micrograph are marked with red squares below. Representative reference-free 2D class averages are shown at the bottom. (**b**) Overview of cryo-EM data processing workflow for the WT RC–LH1 monomer dataset. Every major step is shown with the associated number of particles, percentage of particles per class, or estimated resolution. Selected 3D class that went into further processing is marked with a rectangle. (**c**) Fourier Shell Correlation (FSC) curves including FSC between two independently refined half-maps generated by RELION (blue) and model-to-map FSC generated by Phenix (red). Green line: FSC of the model refined against the 1st half map versus the 1st half map; Yellow line: FSC of the model refined against the 1st half map versus the 2nd half map. Global resolution values were calculated according to the gold-standard FSC = 0.5 and 0.143. (**d**) Angular distribution of the particles used to generate the final electron density map. The direction and length of each cylinder represent the view and the number of particles respectively. (**e**) Local resolution of the cryo-EM map as seen from the periplasmic view (left), side view (middle), and cytoplasmic view (right), estimated by RELION 3.1.

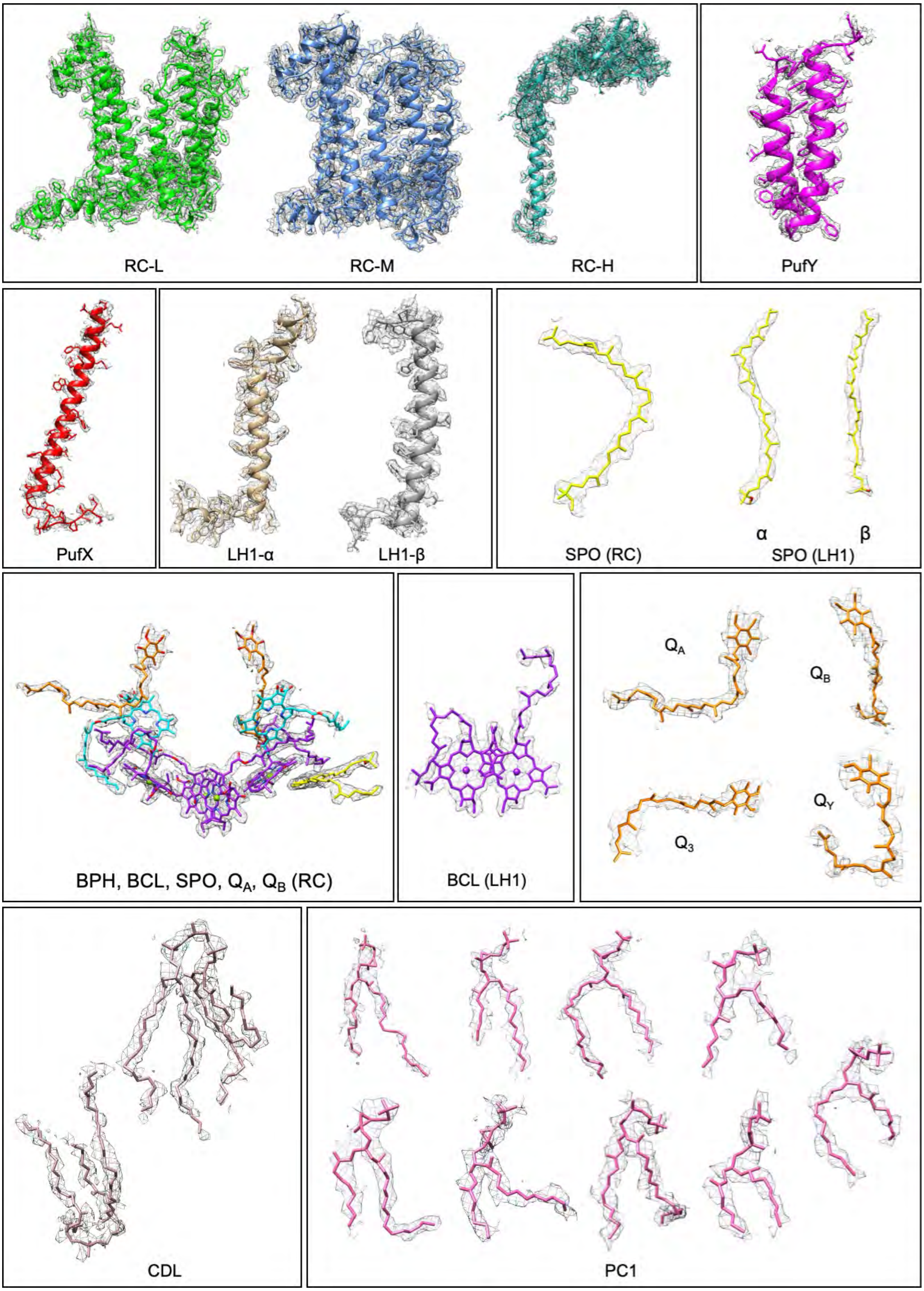


**Fig. S4. Cryo-EM map densities and structural models of protein peptides and cofactors in the WT RC–LH1 monomer.**

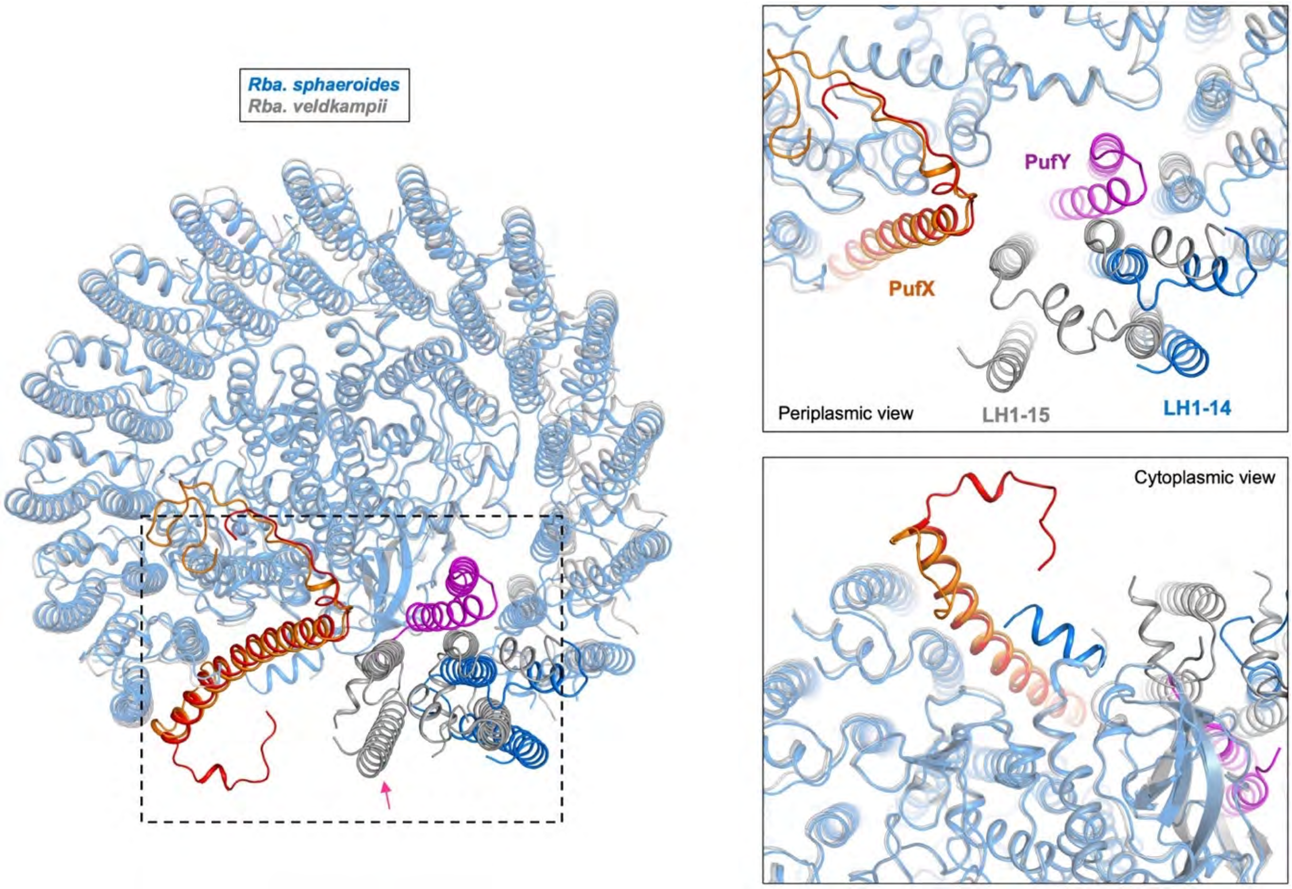


**Fig. S5. Comparison of the RC–LH1 monomers from *Rba. sphaeroides* and *Rba. veldkampii* (PDB ID: 7DDQ).** *Rba. sphaeroides* model is colored in blue, with PufX colored in red and PufY colored in magenta. *Rba. veldkampii* model is colored in grey, with PufX colored in orange. The 15$^{th}$ LH1 subunit (arrow) is present in the RC–LH1 monomer from *Rba. veldkampii* but is missing in *Rba. sphaeroides*. On the right, the top panel shows a close-up periplasmic view of the area around PufY at the periplasmic side; the bottom panel shows a focused cytoplasmic view of the PufX N-terminus at the cytoplasmic side.

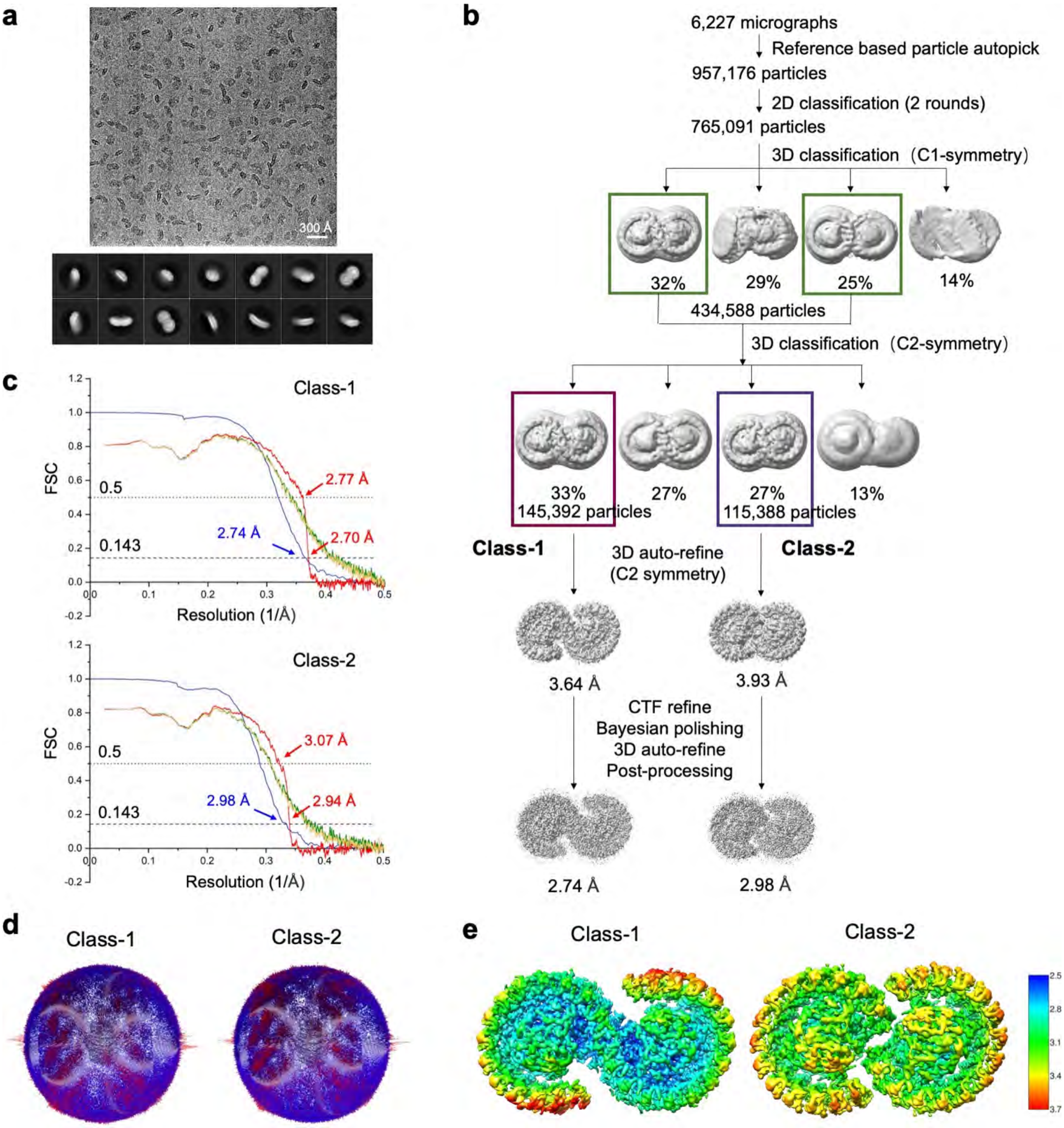


**Fig. S6. Cryo-EM data process of the *Rba. sphaeroides* RC–LH1 dimers.** (**a**) Representative cryo-EM micrograph of RC-LH dimeric supercomplex (upper) and representative 2D class averages (lower). The square size in 2D class average image is 300 Å × 300 Å. (**b**) Cryo-EM data processing workflow. (**c**) Gold-standard FSC curves of the Class-1 and Class-2 dimers, including FSC between two independently refined half-maps generated by RELION (blue) and model-to-map FSC generated by Phenix (red). Green line: FSC of the model refined against the 1$^{st}$ half map versus the 1$^{st}$ half map; Yellow line: FSC of the model refined against the 1$^{st}$ half map versus the 2$^{nd}$ half map. Global resolution values were calculated according to FSC = 0.5 and 0.143. (**d**) Angular distribution of Class-1 (left) and Class-2 (right) in the final 3D refinement with C2 symmetry. (**e**) Local resolution of Class-1 (left) and Class-2 (right) maps.

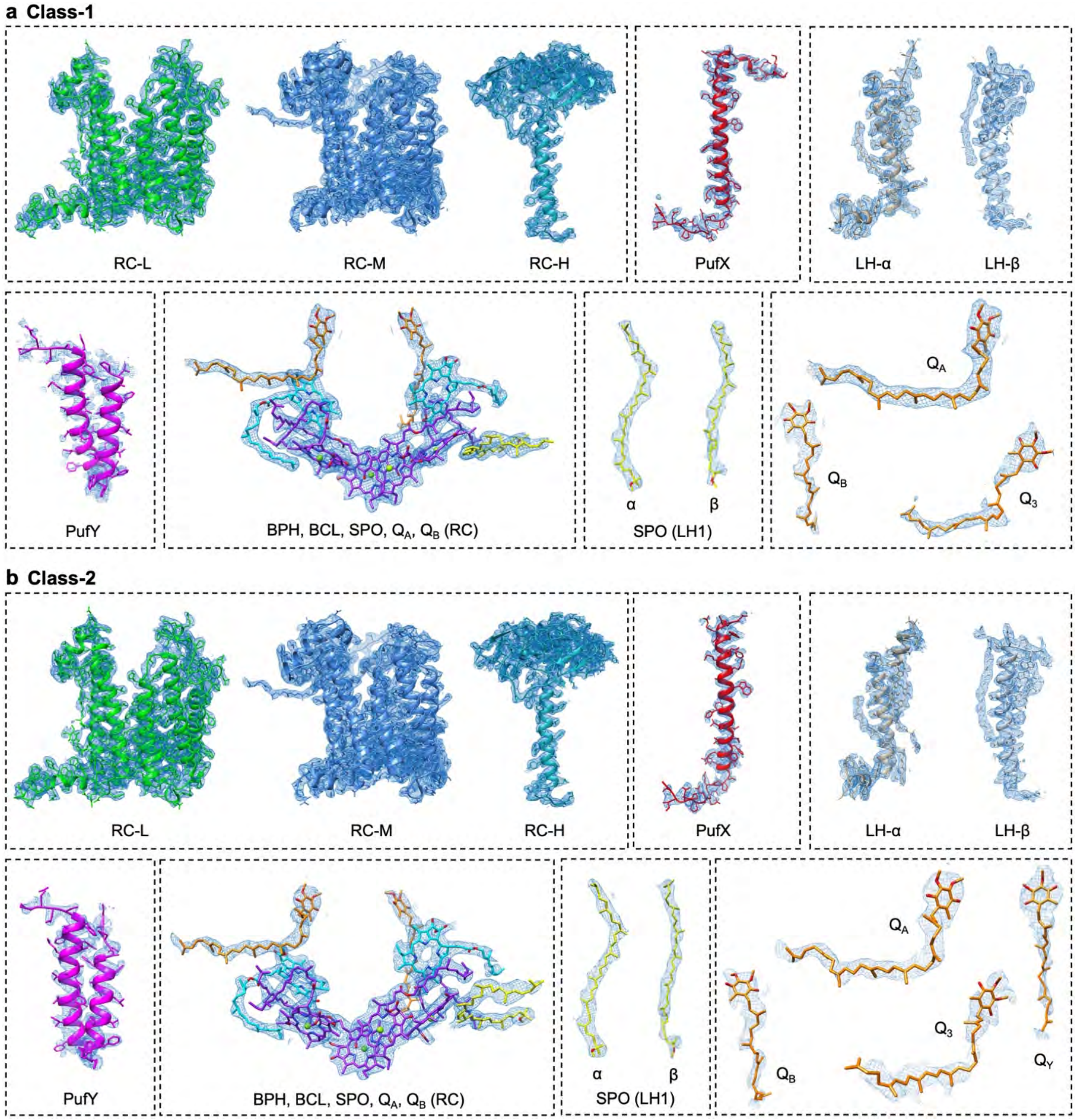


**Fig. S7. Cryo-EM map densities and structural models of protein peptides and cofactors in the RC–LH1 dimers.** (**a**) Cryo-EM densities of subunits and cofactors in Class-1. (**b**) Cryo-EM densities of subunits and cofactors in Class-2.

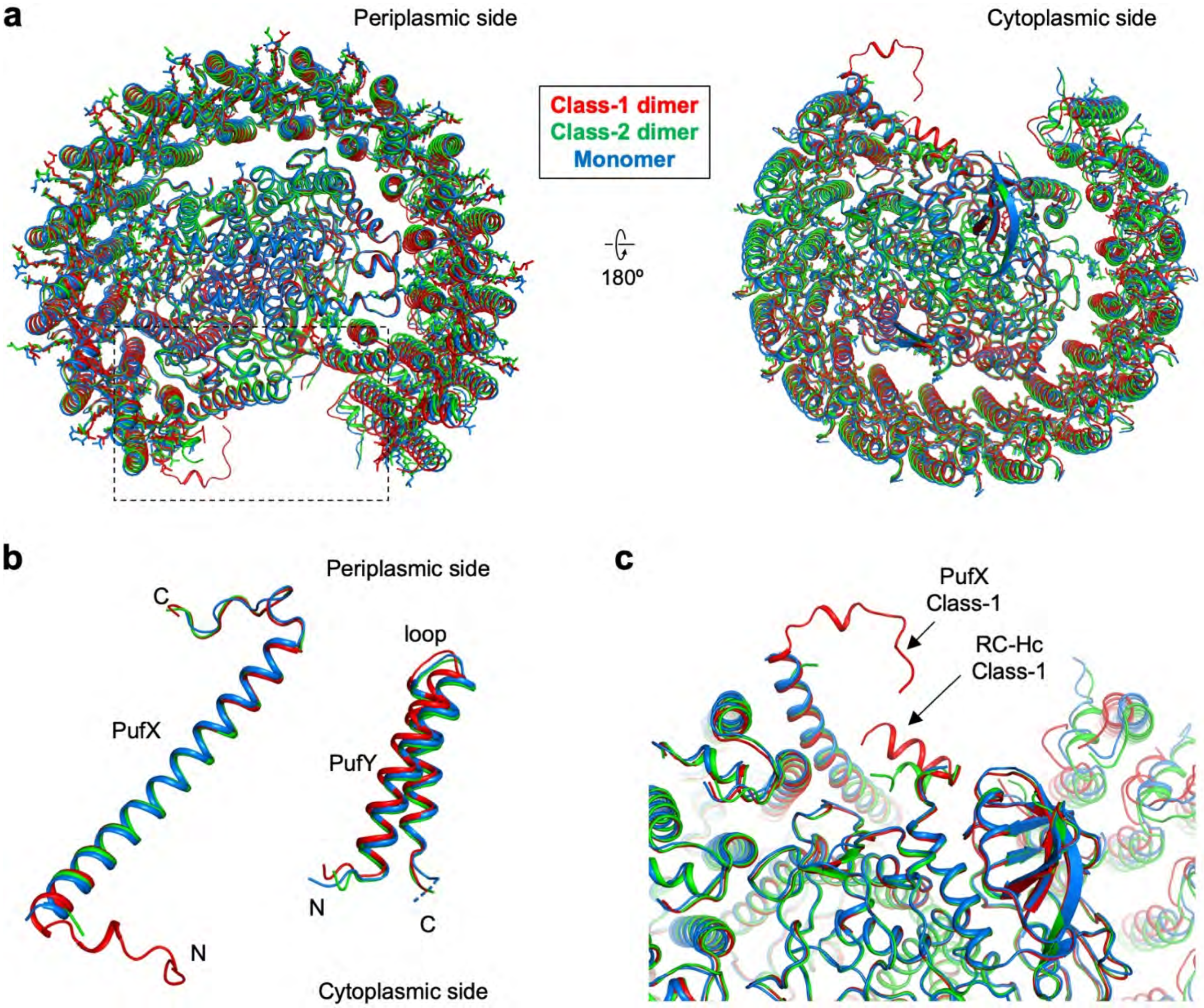


**Fig. S8. Comparison of the monomeric structure of the WT Class-1 and Class-2 dimers and monomer superimposing on their RC subunits.** (**a**) Superimposition of a monomer from the Class-1 dimer (red), Class-2 dimer (green), and RC−LH1 monomer (blue), as seen from the periplasmic side (left) and the cytoplasmic side (right). (**b**) Focused view of PufX and PufY in (**a**). (**c**) Focused view of PufX N-terminal region at the cytoplasmic side from the boxed region in (**a**). The extended tail of PufX and the C-terminus region of RC-H (RC-Hc) in the Class-1 dimer are marked with arrows.

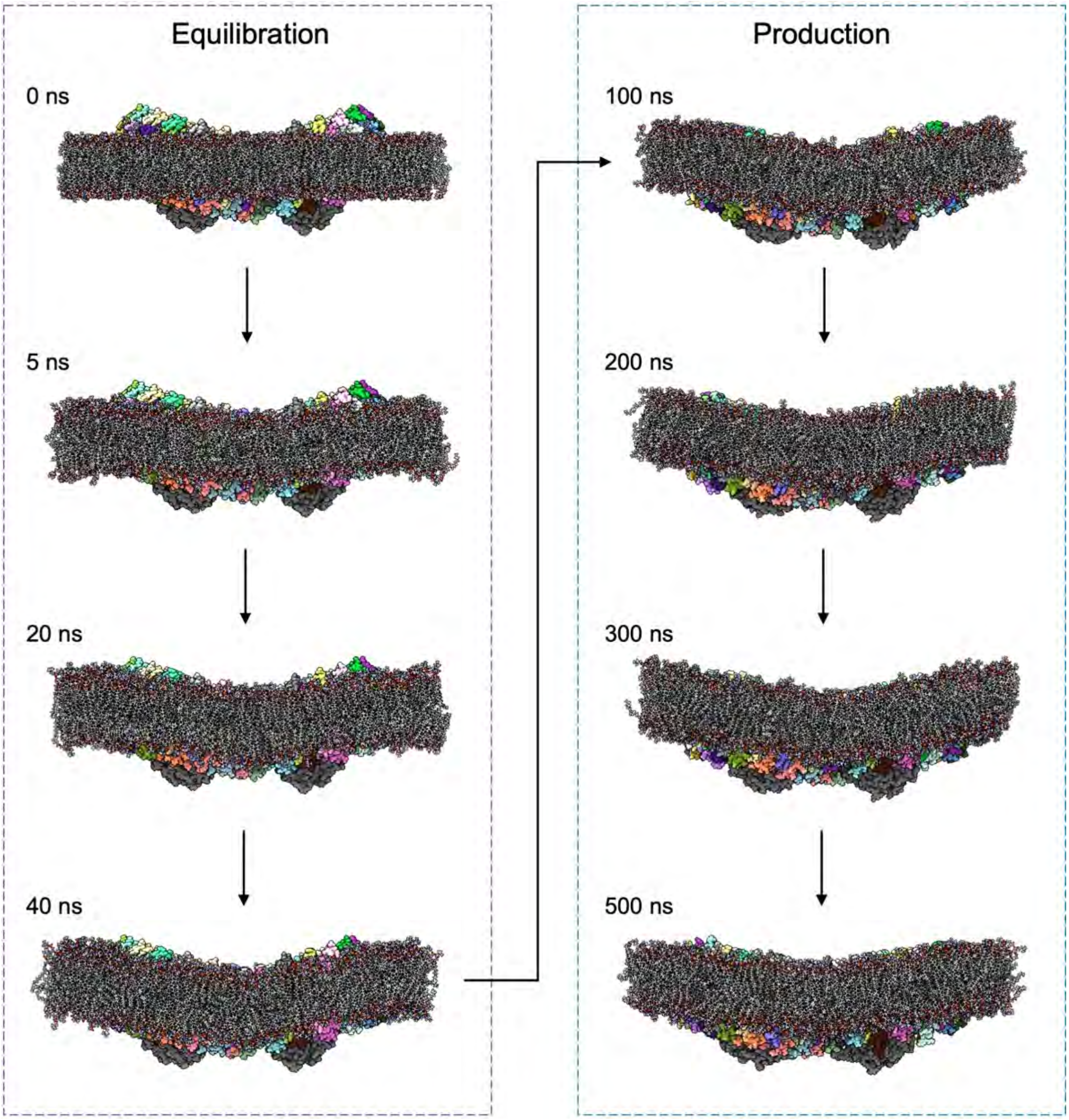


**Fig. S9. Computational simulations of dimer-induced membrane curvature.** Side views of the structures of the RC–LH1 dimer embedded in a lipid bilayer at 0, 20, and 40 ns of the equilibration and at 500 ns of the production all-atom molecular dynamics (AAMD) simulations are shown. Equilibration 0 ns to 40 ns represents the membrane curvature induced by the bent conformation of the RC–LH1 dimer (left). The curved structure of the lipid bilayer was maintained during the 500-ns unrestrained AAMD simulations. Protein chains are shown with molecular surfaces colored by chain. The atoms of ligands and lipids are shown in sticks. Hydrogen, carbon, nitrogen, oxygen, and phosphorous atoms are colored in white, gray, blue, red, and orange, respectively. Images were produced by UCSF ChimeraX.

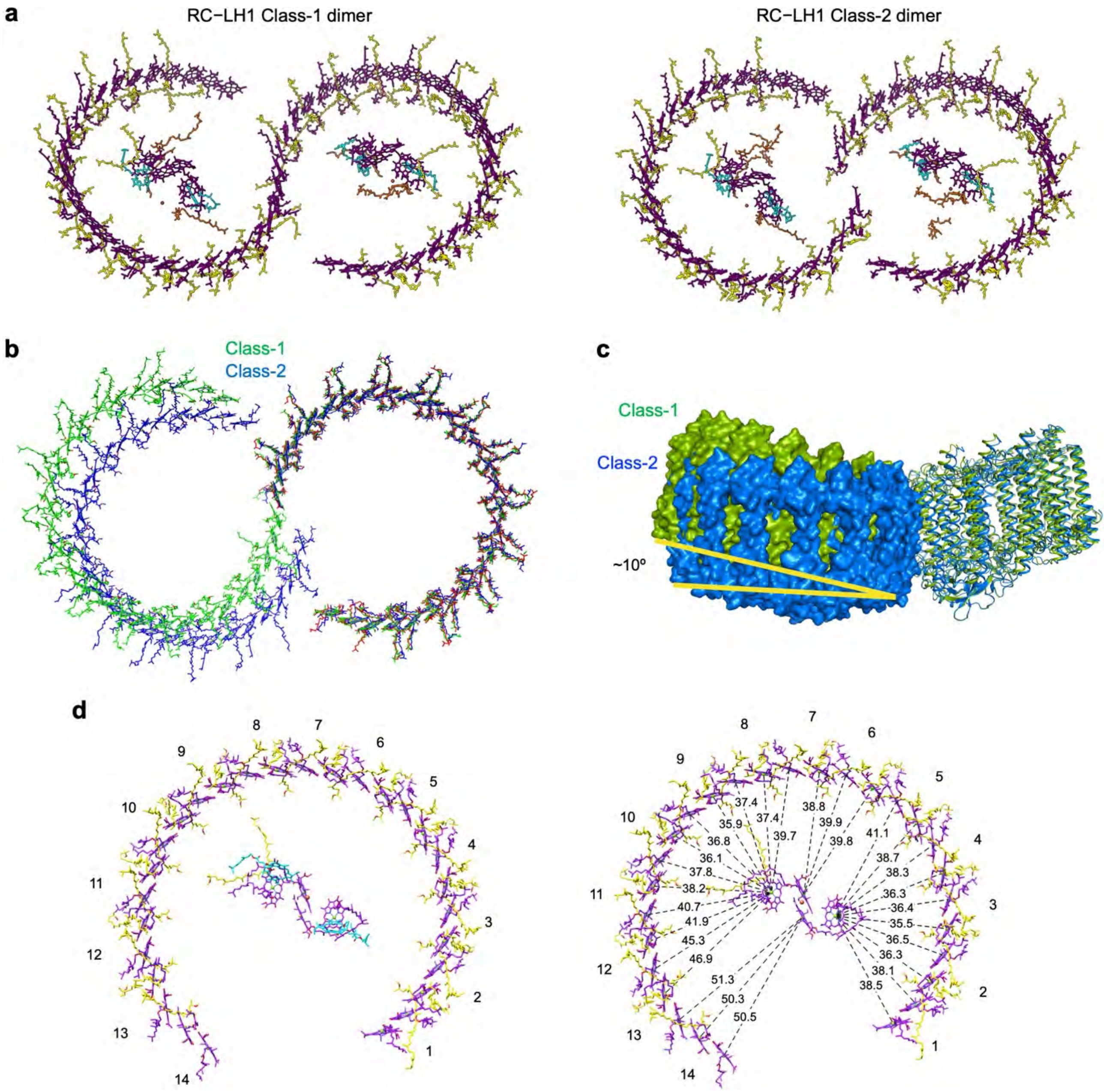


**Fig. S10. Pigment arrangement and comparison of the two classes of the WT RC–LH1 dimers from *Rba. sphaeroides*.** (**a**) Pigment arrangement within the WT RC–LH1 Class-1 dimer (left) and Class-2 dimer (right). BChls are colored in purple, SPOs in yellow, BPhes in cyan, and UQ-10s in orange. (**b**) LH1 ring associated pigment comparison between Class-1 (green) and Class-2 (blue). Monomers on the right of each model were superimposed and the other monomer followed in its relation to the first. (**c**) Comparison of the tilted conformations of Class-1 (green) and Class-2 (blue). Monomers on the right of each model were superimposed and the other monomer followed in its relation to the first. (**d**) Pigment arrangement within the RC–LH1 monomer. The LH1 subunit numbers are depicted on the outside. On the right, the Mg-Mg distance of each LH1 BChl to the nearest RC BChl is shown in Å. BChls are colored in purple, SPOs in yellow, BPhes in cyan.

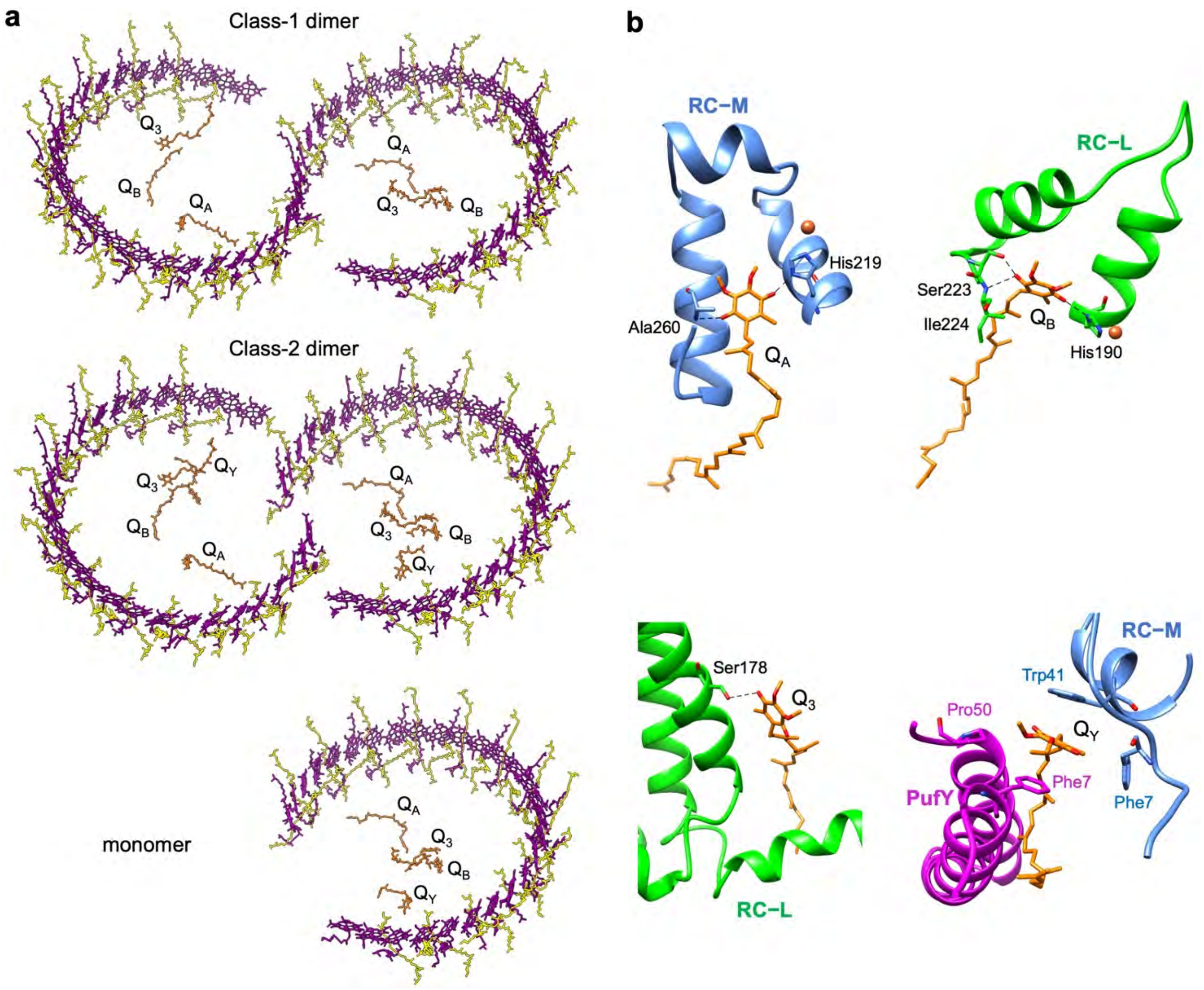


**Fig. S11. Quinones in the *Rba. sphaeroides* RC–LH1 complexes.** (**a**) Positions of identified quinone molecules in respect to the LH1 ring in the Class-1 dimer (top), the Class-2 dimer (middle), and the WT monomer (bottom). (**b**) Focused views of each identified quinone. The residues they interact with are marked and shown in sticks. The head group of $Q_Y$ is surrounded by a specific pocket formed by the aromatic rings of the Trp41 and Phe7 residues of RC-M and the Phe7 and Pro50 residues of PufY.

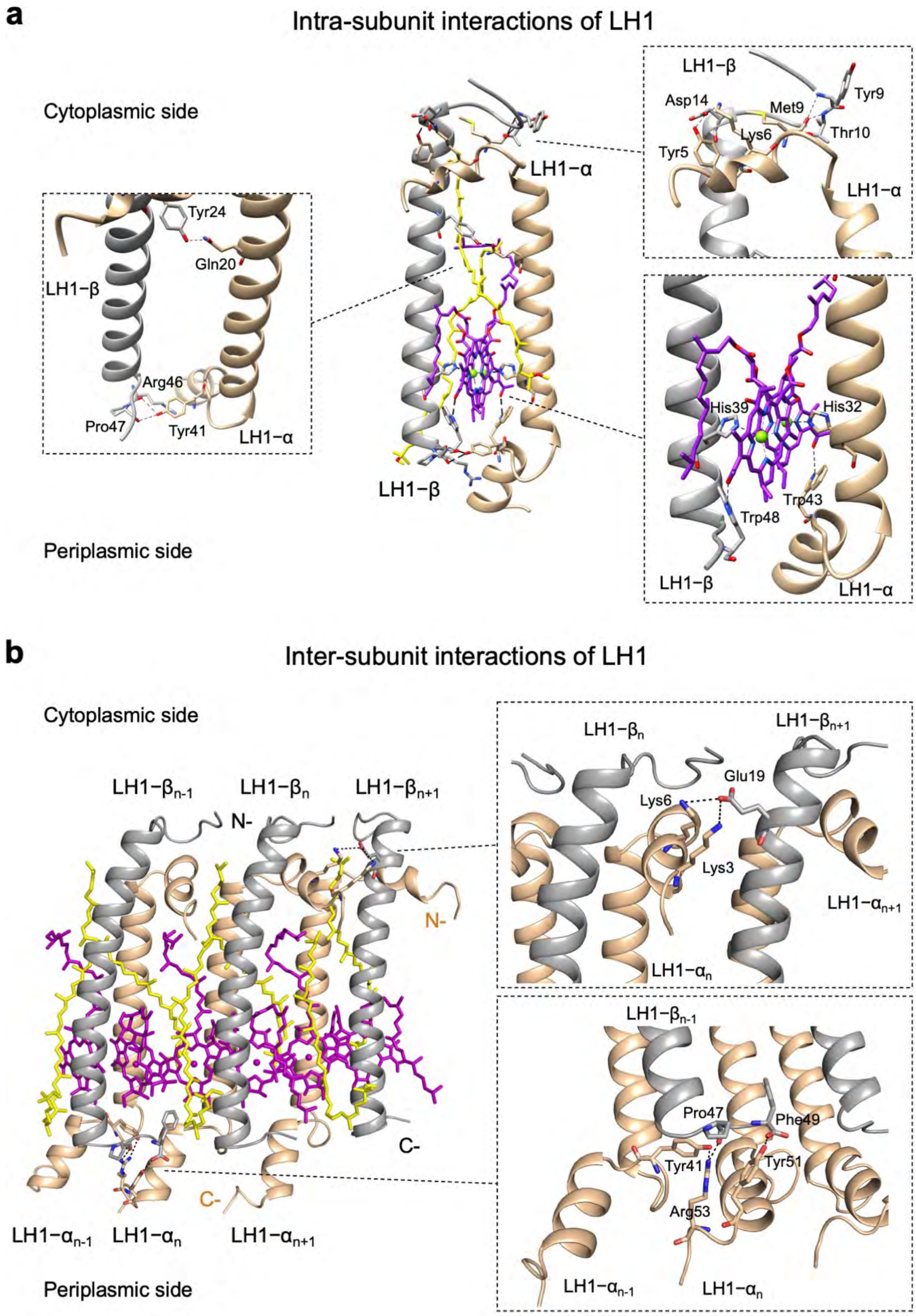


**Fig. S12. Interactions within the LH1 subunit and between neighboring LH1 subunits.** (**a**) Representative LH1 subunit organization. Associated pigments (BChl – purple, SPO – yellow) and interacting residues are shown in sticks. Close-up views of interactions within an LH1 subunit are shown in boxes on the side. (**b**) Representative LH1-LH1 interface. Associated pigments (BChl – purple, SPO – yellow) and interacting residues are shown in sticks. Close-up views of interactions between neighboring LH1 subunits are shown in boxes on the side. Interaction types and distances are displayed in Table S3.

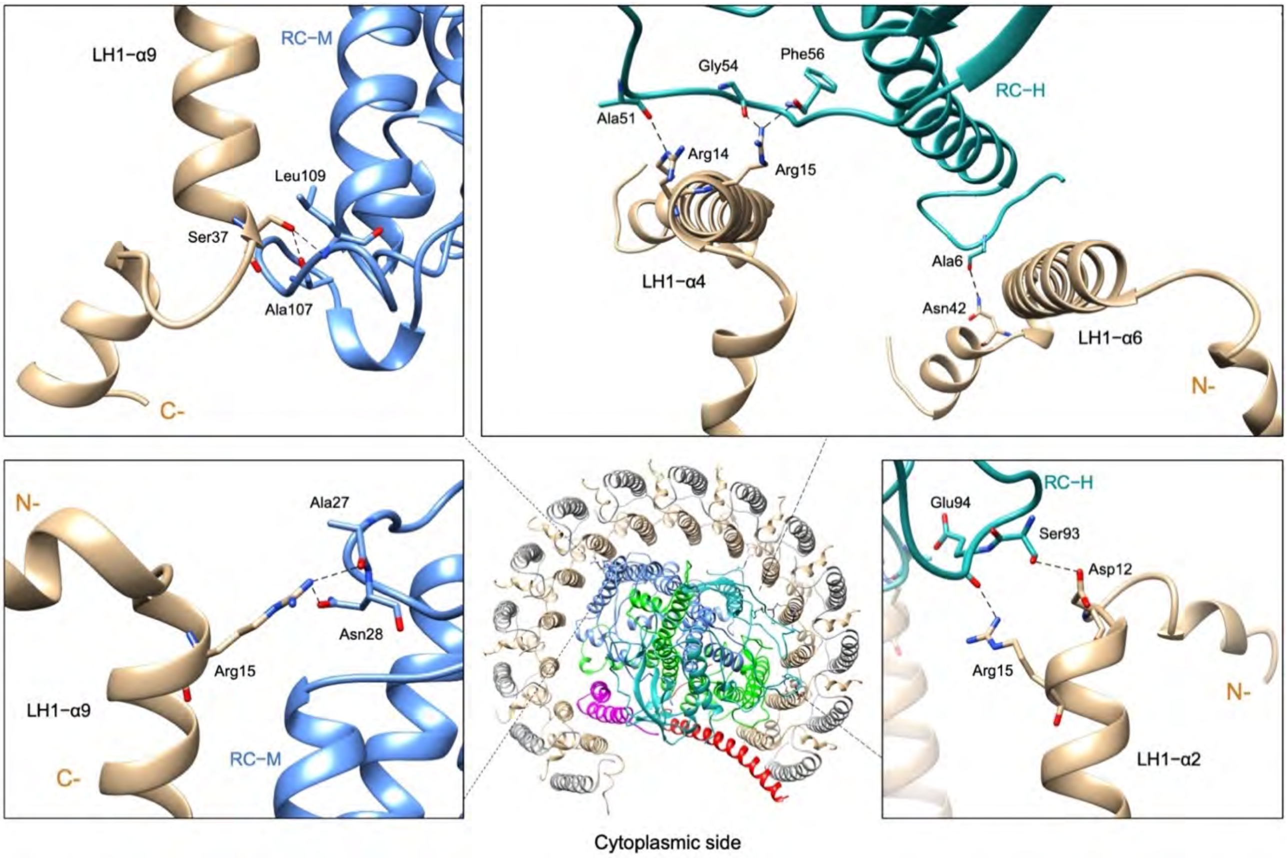


**Fig. S13. Interactions between the RC and LH1 in the *Rba. sphaeroides* RC–LH1 complex.** Interacting residues are shown in sticks. Detailed views of interactions are displayed in the boxes on the side while the middle view shows their general position within the complex. Types and distances of interactions between the RC and the LH1 are displayed in Table S3.

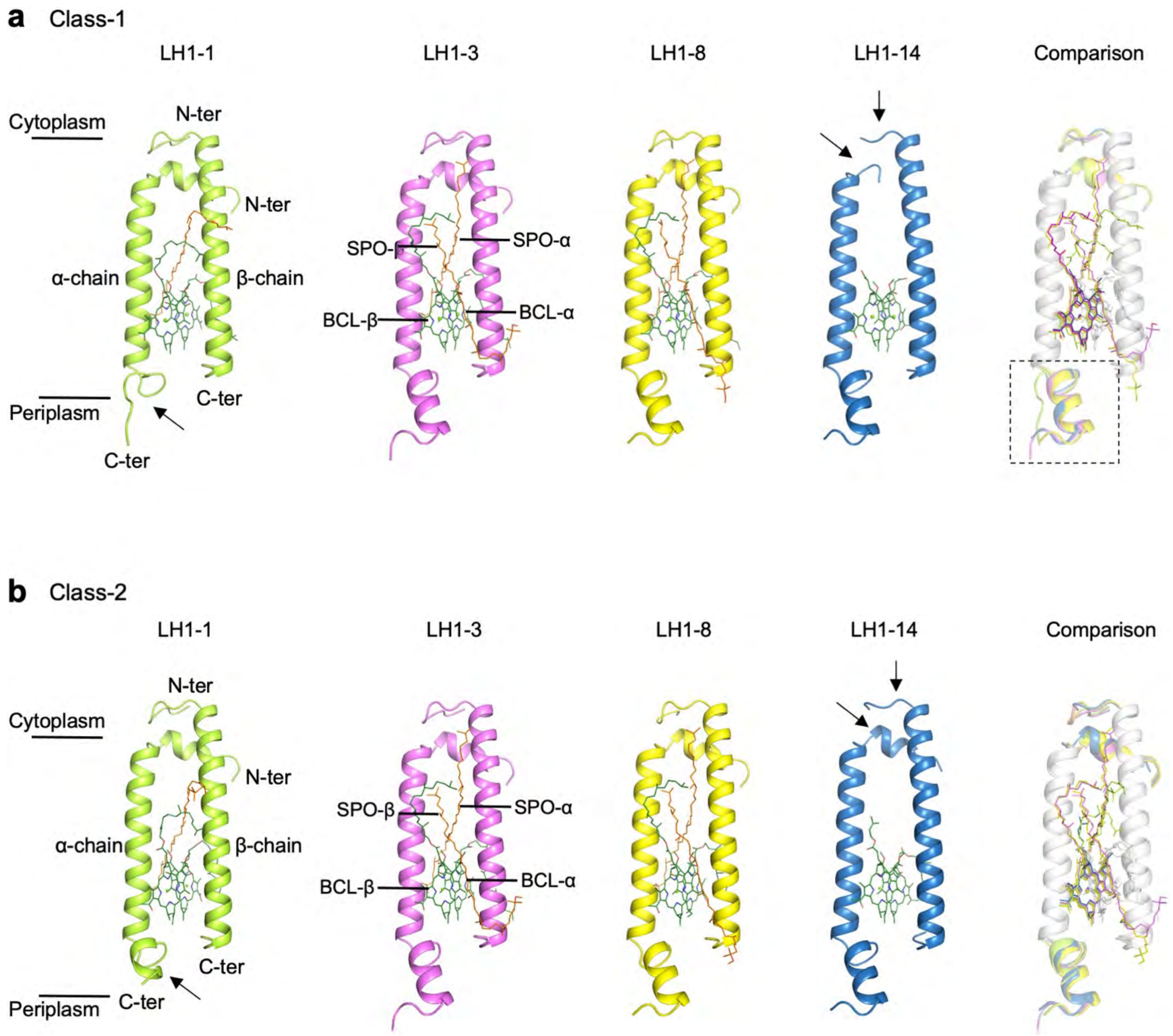


**Fig. S14. Structural comparison of the 14 LH1 αβ-heterodimers of *Rba. sphaeroides*.** (**a**) Overall structures and comparison of the LH1 subunits in the Class-1 dimer, including LH1-1 with 2 BChls and 1 SPO-α, LH1-14 with only 2 BChls, as well as LH1-3 and LH1-8 with 2 BChls and 2 SPOs representing other 12 similar LH1 heterodimers (LH1-2~LH1-13). The N-terminus and C-terminus are labeled in the LH1-1. SPO and BCL are labeled in the LH1-3. The C-terminus of LH1-1α and the N-termini of LH1-14α/β (arrows) exhibit distinct conformations compared with those of other LH1 polypeptides. The phytol tail of BCL-β and the SPO-α molecules in LH1-1 adopt different conformation than those in other LH1 subunits, enabling the association of PufX. Moreover, LH1-1 lacks SPO-β, which also facilitates its binding with PufX as otherwise the SPO molecule would clash with PufX. The SPO molecules in LH1-14 exhibit poor density, likely due to the terminal location and hence a higher mobility, and thus were not built in the structure. The tail regions of SPO-β molecules are highly flexible since they reach out into the solvent zone and adopt two major conformations as shown in LH1-3 and 1-8. Structural comparison of individual LH1 subunits is shown on the right. The similar part of LH1s is colored white and the different parts are highlighted in colors. (**b**) Structures and comparison of the LH1 subunits in the Class-2 dimer, including LH1-1 with 2 BChls and 1 SPO-α, LH1-14 with only 2 BChls, as well as LH1-3 and LH1-8 with 2 BChls and 2 SPOs representing other 12 similar LH1 heterodimers (LH1-2~LH1-13).

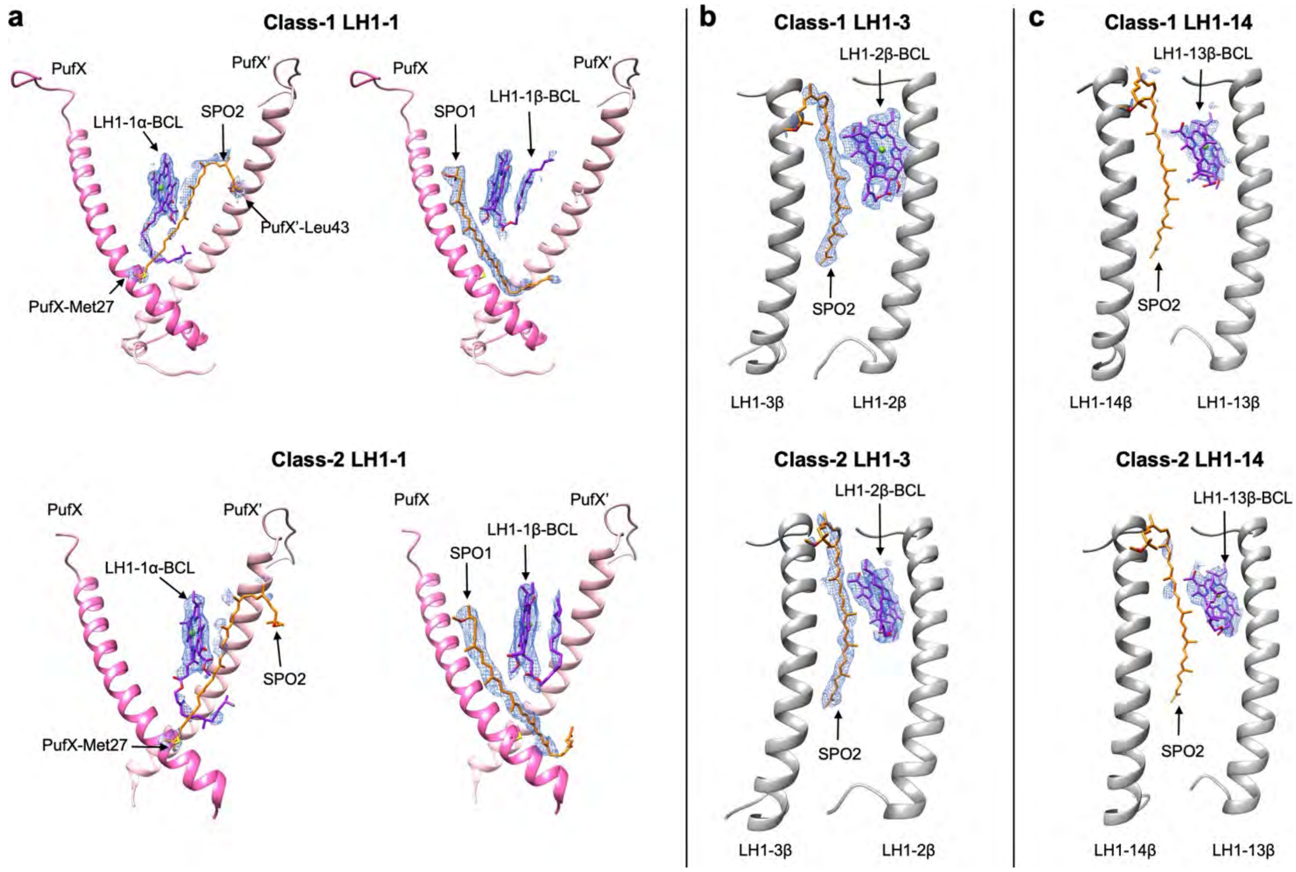


**Fig. S15. Densities around the position of potential SPO2 molecule in LH1-1 and LH1-14, and the SPO2 density in LH1-3 in both Class-1 and Class-2 dimers.** (**a**) Densities of the potential SPO2 and other cofactors in LH1-1 (shown in two panels for clarity). Left panel shows the densities of BChl of LH1-1α and the potential SPO2 molecule. Right panel shows the densities of BCL of LH1-1β and the SPO1 molecule. Compared with other cofactors, the potential SPO2 shows no density. In addition, the potential SPO2 molecule clashes with the phytol chain of BChl of LH1-1α, as well as Met27 of PufX and Leu43 of PufX', suggesting that LH1-1 does not possess SPO2 molecule. PufX and PufX' are shown as ribbons in pink and salmon, respectively. Residues are shown in sticks. (**b**) Densities of SPO2 of LH1-3 and BCL of LH1-2β. LH1-2β and LH1-3β are shown as ribbons. (**c**) Densities of potential SPO2 of LH1-14 and BCL of LH1-13β. LH1-13β and LH1-14β are shown as ribbons.

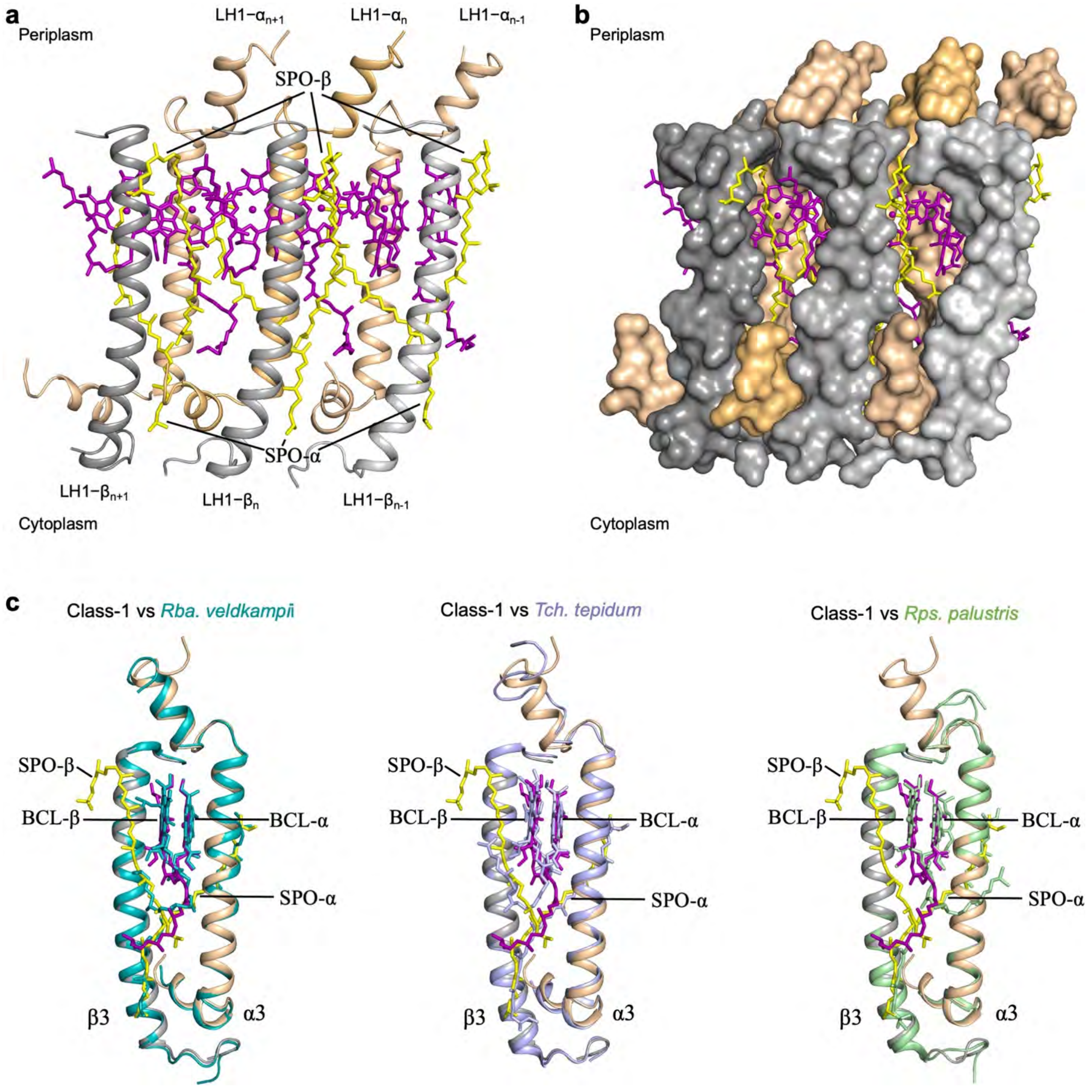


**Fig. S16. Analysis of the LH1 structure.** (**a**) Architecture of three adjacent LH1 αβ-apoproteins and their bound BChls and SPOs. Pigments are involved in the interaction with neighboring LH1s. (**b**) Dense arrangement of pigment molecules within the LH1 barrier, which potentially blocks the shuttle of quinones/quinols across the LH1 ring. (**c**) Structural comparison of LH1 αβ-heterodimers from the *Rba. sphaeroides* Class-1 dimer (represented by LH1-3) with LH1 from *Rba. veldkampii* (PDB ID: 7DDQ), *Tch. tepidum* (PDB ID: 575S), and *Rps. palustris* (PDB ID: 6Z5S). The presence of a second SPO (SPO-β) is unique in the LH1 of *Rba. sphaeroides*.

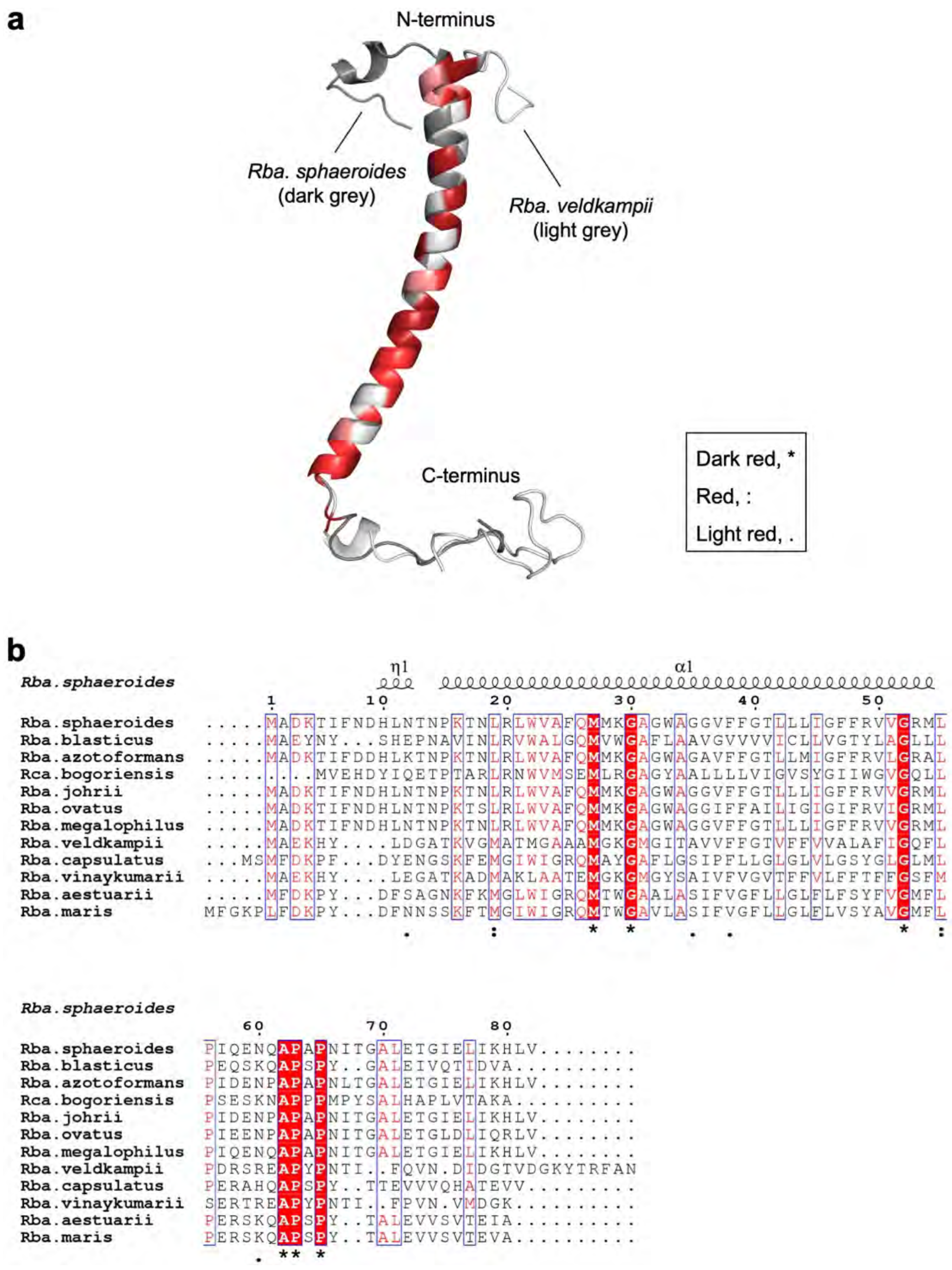


**Fig. S17. Comparison of the PufX polypeptides from *Rba. sphaeroides* and other species.** (**a**) Comparison of the cryo-EM structure of PufX in the RC−LH1 complexes from *Rba. sphaeroides* and *Rba. veldkampii* (PDB ID: 7DDQ). Fully conserved residues are colored in dark red; residues with strongly and weakly similar properties are colored in red and light red, respectively; non-conserved residues are colored in white. (**b**) Sequence alignment of the PufX subunits from *Rba. sphaeroides* and other *Rhodobacter* species. Conserved residues are highlighted.

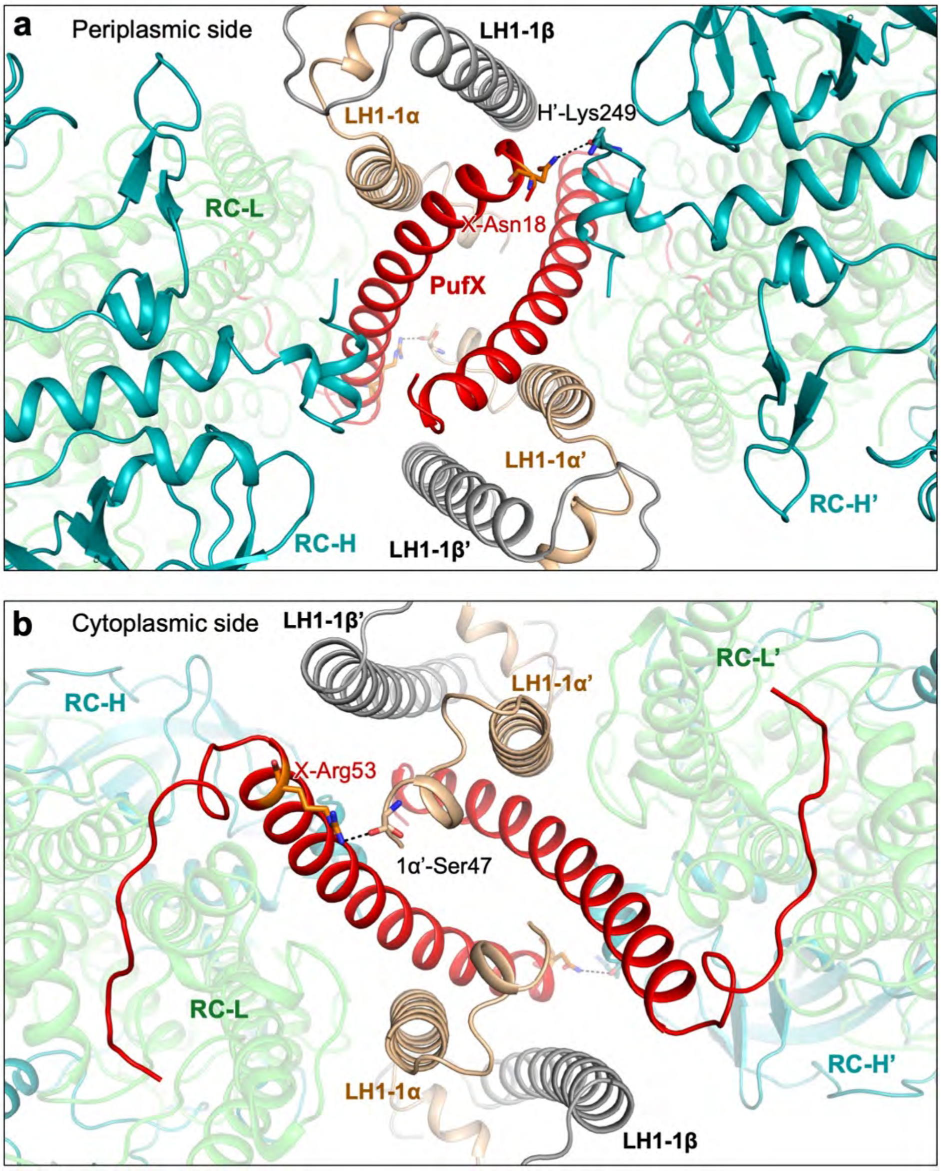


**Fig. S18. The interactions of PufX in the Class-2 dimer.** (**a**) At the periplasmic side, Asn18 of PufX is hydrogen bonded with the Lys249 residue of RC-H from the neighboring monomer. (**b**) At the cytoplasmic side, Arg53 of PufX forms a hydrogen bond with Ser47 of LH1-1α from the neighboring monomer.

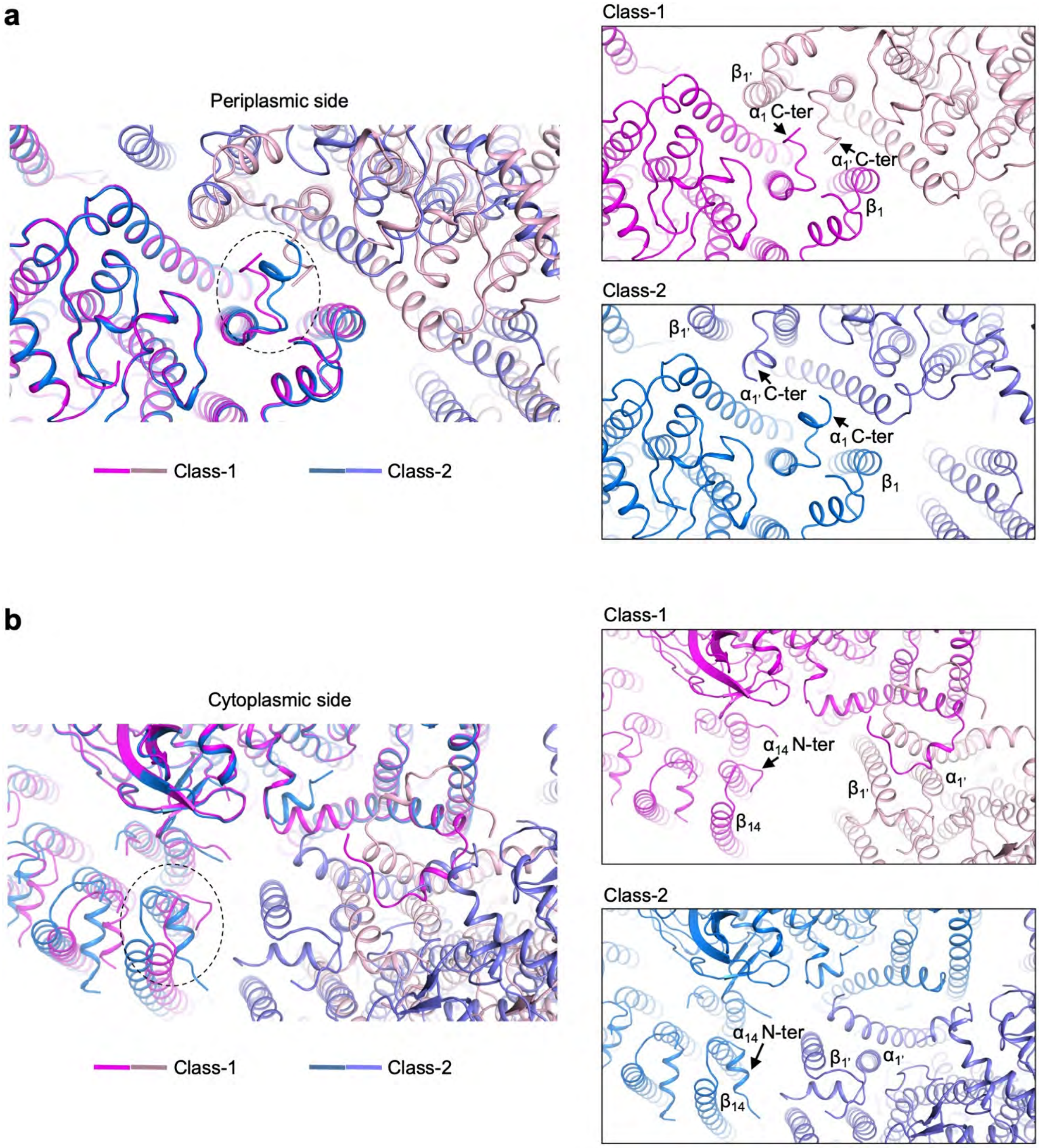


**Fig. S19. Structural comparisons of the LH-1 structures in the two classes of dimers.** (**a**) periplasmic view. (**b**) cytoplasmic view. The monomers within the two classes of dimers exhibit structural differences, mainly at the dimerization interface, in particular the last two LH1s and the C-terminal tails of LH1-$\alpha_1$ as indicated by arrows. At the periplasmic side, the C-terminal tail of LH1-$\alpha_1$ shows slightly different conformations between Class-1 and Class-2. It is hydrogen bonded with the C-terminal region of LH1-$\beta_1$ in Class-1, whereas the two LH1 units are separated without any contacts in Class-2. At the cytoplasmic side, the C-terminal helix of RC-H orientates differently in Class-1 and Class-2. It interacts with the N-terminal region of PufX' in Class-1, whereas the interaction is absent in Class-2. In contrast, the N-terminus of LH1-$\alpha_{14}$ is better identified in Class-2. It forms contacts with LH1-$\beta_{1'}$ in Class-2, whereas they are separated in Class-1. Moreover, the last pairs of LH1 slightly shift outward in Class-2 compared with those in Class-1, forming interactions with the neighboring monomer. The closer association of LH1-13/14 region with the neighboring monomer stabilizes the terminal LH1 units and PufY, thus resulting in clearer densities in Class-2.

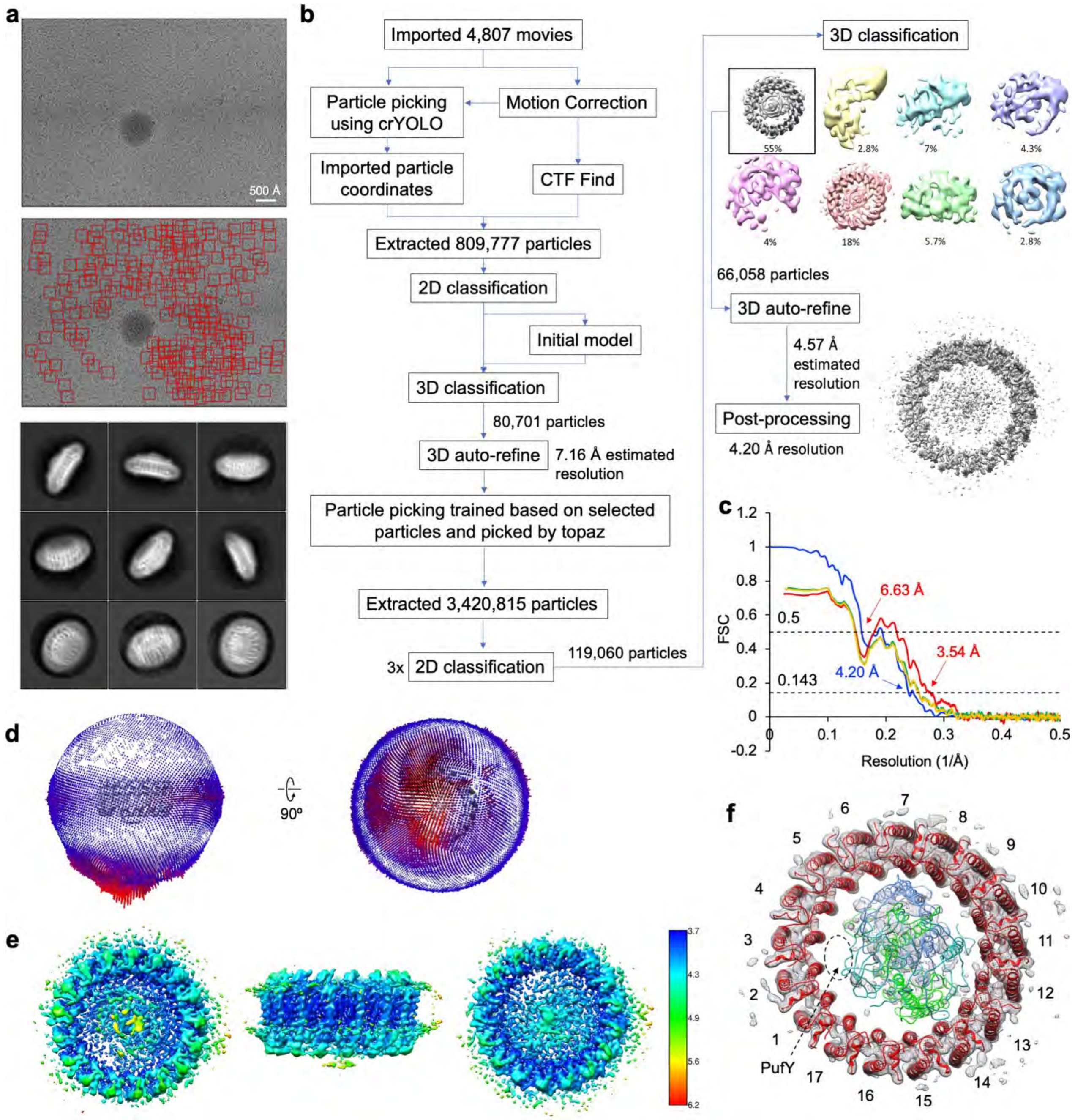


**Fig. S20. Cryo-EM data process of the Δ*pufX* RC–LH1 monomer.** (**a**) Motion-corrected example of a cryo-EM captured movie is shown on top and all particles picked from that micrograph are marked with red squares below. Representative reference-free 2D class averages are shown at the bottom. (**b**) Overview of cryo-EM data processing workflow for the Δ*pufX* RC–LH1 dataset. Every major step is shown with the associated number of particles, percentage of particles per class, or estimated resolution. Selected 3D class that went into further processing is marked with a rectangle. (**c**) FSC between two independently refined half-maps generated by RELION 3.1 (blue) and model-to-map FSC generated by Phenix (red). Green line: FSC of the model refined against the 1$^{st}$ half map versus the 1$^{st}$ half map; Yellow line: FSC of the model refined against the 1$^{st}$ half map versus the 2$^{nd}$ half map. Global resolutions of 6.62 Å and 6.56 Å were calculated using the FSC cut-off of 0.143. (**d**) Angular distribution of the particles used to generate the final electron density map. The direction and length of each cylinder represent the view and the number of particles respectively. (**e**) Local resolution of the cryo-EM map as seen from the periplasmic view (left), side view (middle), and the cytoplasmic view (right), estimated by RELION 3.1. (**f**) Cryo-EM map densities of the Δ*pufX* RC–LH1 monomer and the fitting of 17 LH1 subunits and RC subunits. The internal structures of the Δ*pufX* RC–LH1 monomer exhibit poor densities probably due to unstable association with LH1 in the absence of PufX. Hence, PufY was not built in the structure, and its possible location was indicated by a dashed circle and arrow.

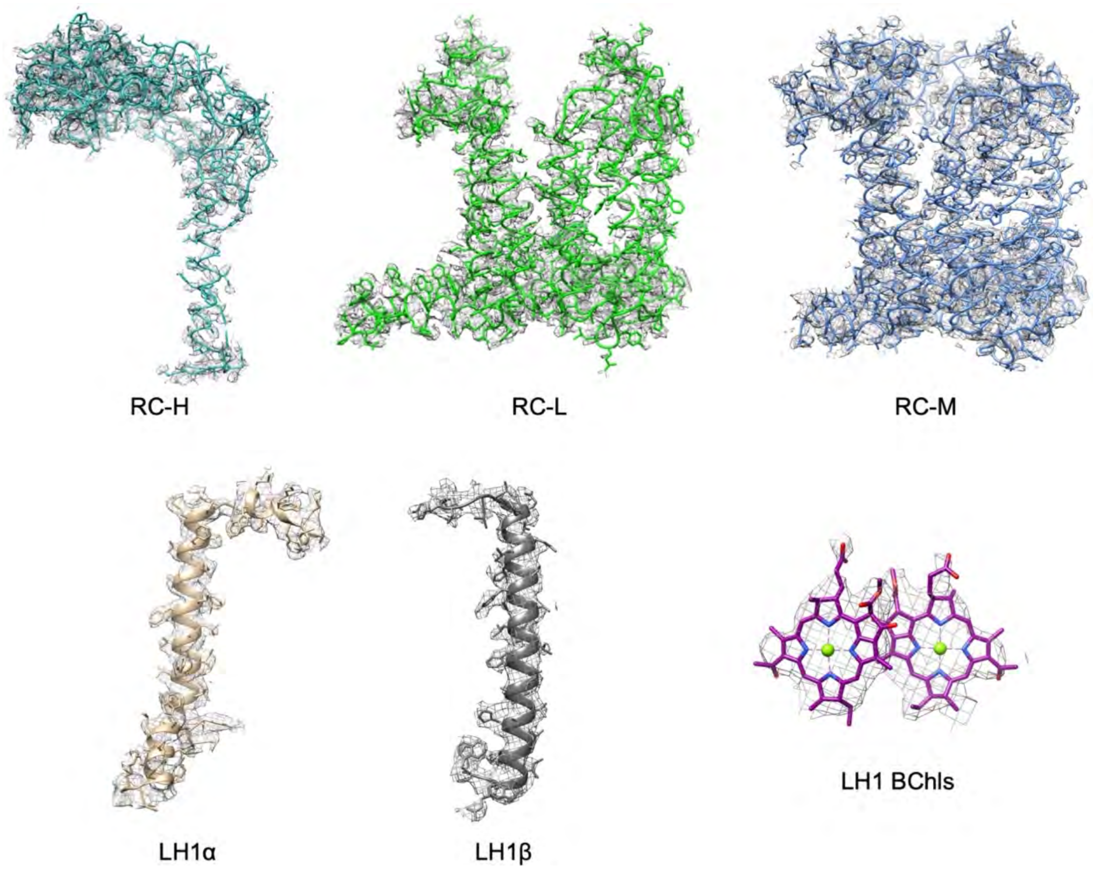


**Fig. S21. Cryo-EM map densities and structural models of protein peptides and cofactors in the Δ*pufX* RC–LH1 monomer.**

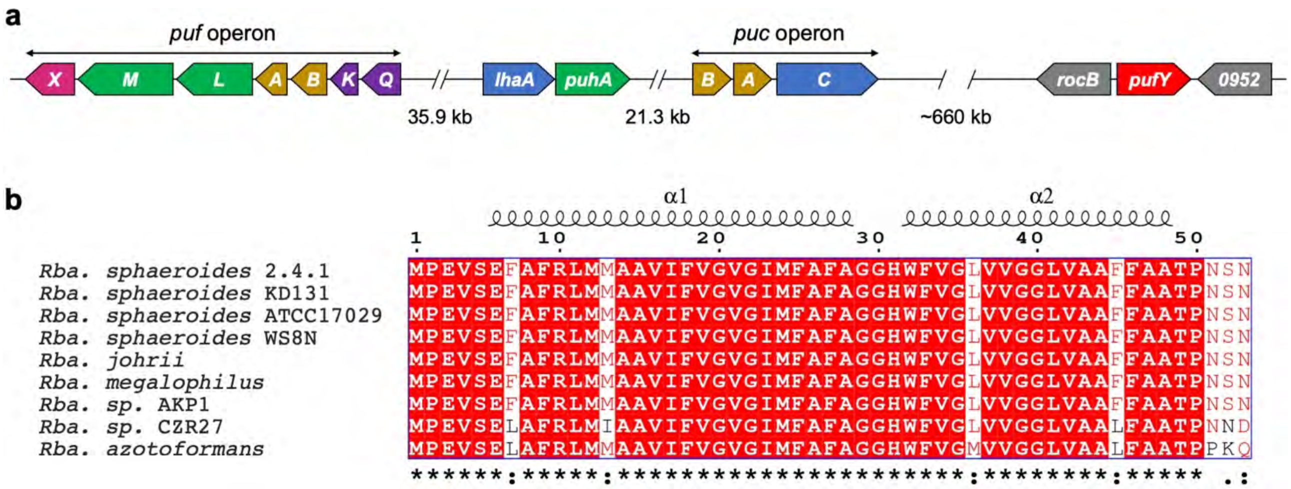


**Fig. S22. Bioinformatic analysis of the *pufY* gene in the genome of *Rba. sphaeroides* (a) and PufY protein sequence alignment (b).** PufY was previously annotated as a hypothetical protein in the genome of *Rba. sphaeroides*. Fully conserved residues are highlighted and indicated as "*". ":" indicates the residues with strongly similar properties. "." indicates the residues with weakly similar properties.

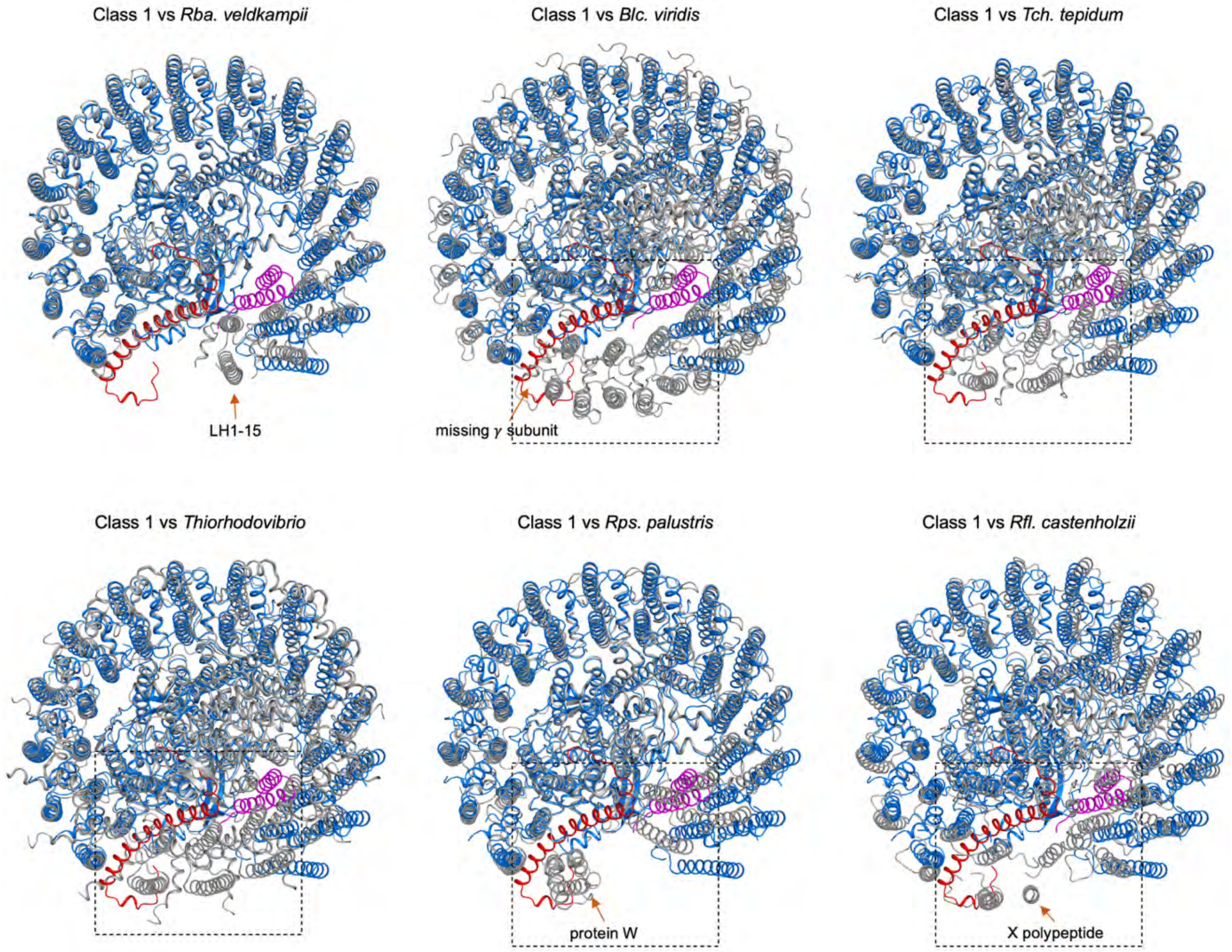


**Fig. S23. Structural diversity of the photosynthetic RC−LH1 core complexes from different species.** The *Rba. sphaeroides* RC−LH1 monomer in the Class-1 dimer is shown in blue and other RC−LH1 structures are shown in grey. In addition to the presence of PufY (purple), the main structural differences were observed at the large opening regions formed by PufX (red). Compared with the *Rba. sphaeroides* RC−LH1 monomer, the RC−LH1−PufX monomer (PDB ID: 7DDQ) of *Rba. veldkampii* has an extra LH1 pair (arrow, 15 LH1 subunits in total); the *Blc. viridis* RC−LH1 (PDB ID: 6ET5) has a small gap in the LH1 ring due to the absence of a γ subunit (arrow); the RC−LH1 structures from *Tch. tepidum* (PDB ID: 5Y5S) and *Thiorhodovibrio* strain 970 (PDB ID: 7C9R) have a closed LH1 ring; the *Rps. palustris* RC−LH1 (PDB ID: 1PYH) has a gap in the LH1 ring created by a protein W (arrow); the *Roseiflexus* (*Rfl.*) *castenholzii* RC−LH1 (PDB ID: 5YQ7) has a gap in the LH1 ring created by an X polypeptide (arrow).

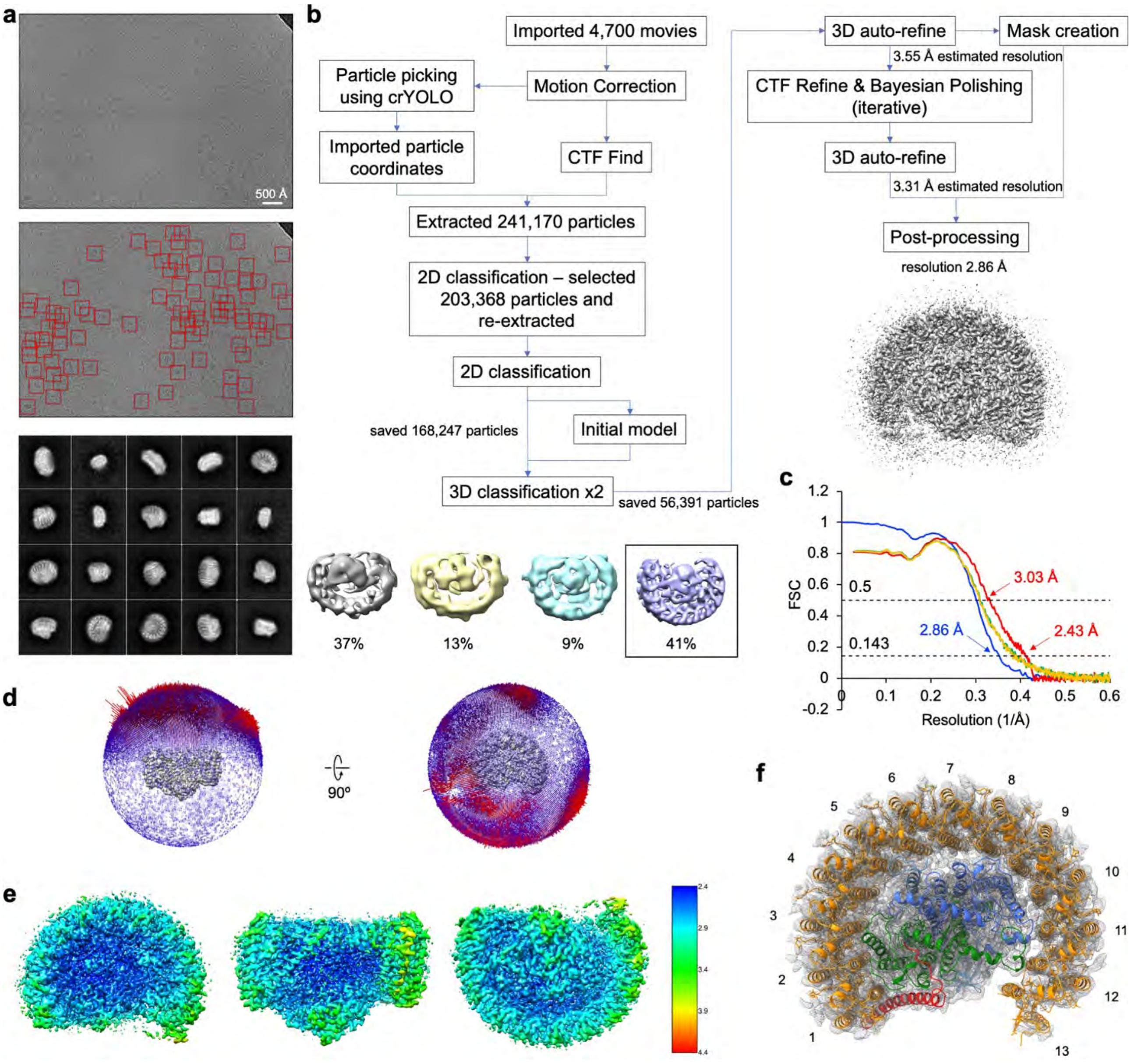


**Fig. S24. Cryo-EM data process of the Δ*pufY* RC–LH1 monomer.** (**a**) Motion-corrected example of a cryo-EM captured movie is shown on top and all particles picked from that micrograph are marked with red squares below. Representative reference-free 2D class averages are shown at the bottom. (**b**) Overview of cryo-EM data processing workflow for the Δ*pufY* RC–LH1 monomer dataset. Every major step is shown with the associated number of particles, percentage of particles per class, or estimated resolution. Selected 3D class that went into further processing is marked with a rectangle. (**c**) FSC curves including FSC between two independently refined half-maps generated by RELION 3.1 (blue) and model-to-map FSC generated by Phenix (red). Green line: FSC of the model refined against the 1st half map versus the 1st half map; Yellow line: FSC of the model refined against the 1st half map versus the 2nd half map. Global resolutions of 2.86 Å and 2.43 Å were calculated using the FSC cut-off of 0.143. (**d**) Angular distribution of the particles used to generate the final electron density map. The direction and length of each cylinder represent the view and the number of particles respectively. (**e**) Local resolution of the cryo-EM map as seen from the periplasmic view (left), side view (middle), and the cytoplasmic view (right), estimated by RELION 3.1. (**f**) Cryo-EM map densities and structural model of protein peptides in the Δ*pufY* RC–LH1 monomer, indicating the presence of PufX and 13 pairs of LH1 αβ-subunits.

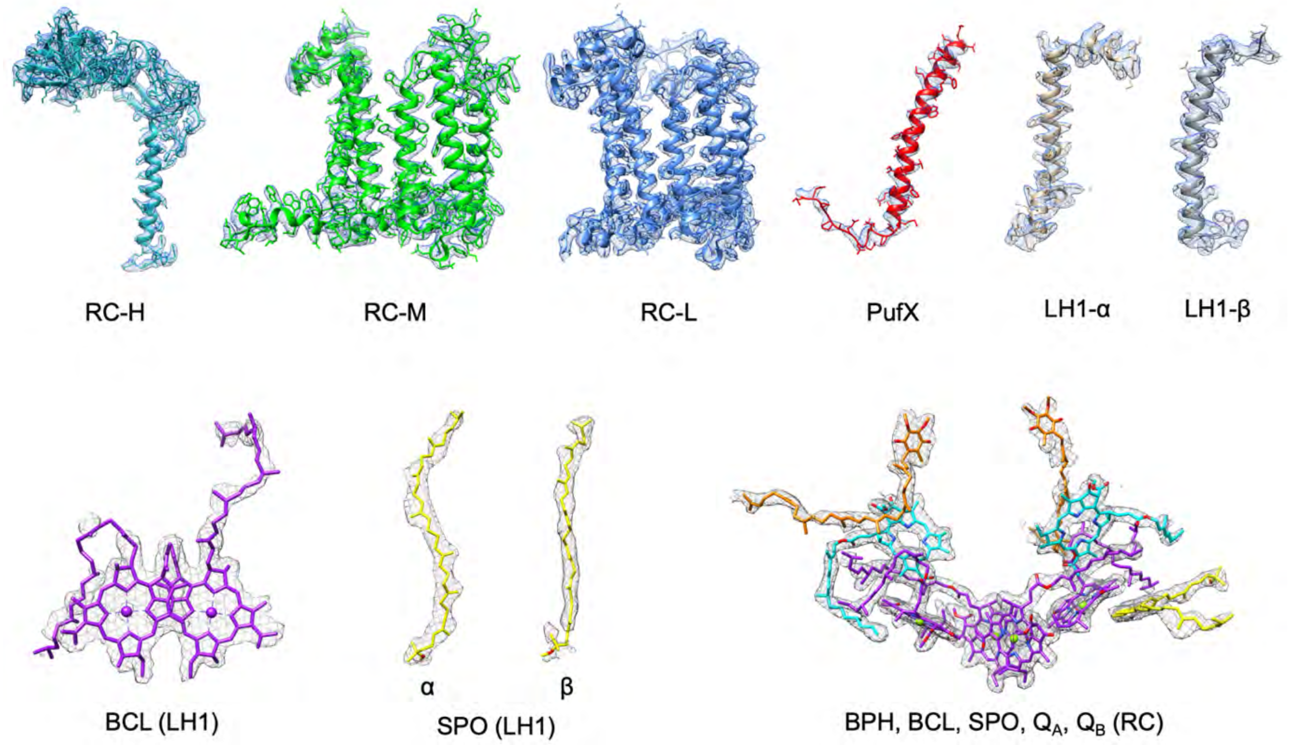


**Fig. S25. Cryo-EM map densities and structural models of protein peptides and cofactors in the Δ*pufY* RC–LH1 monomer.**

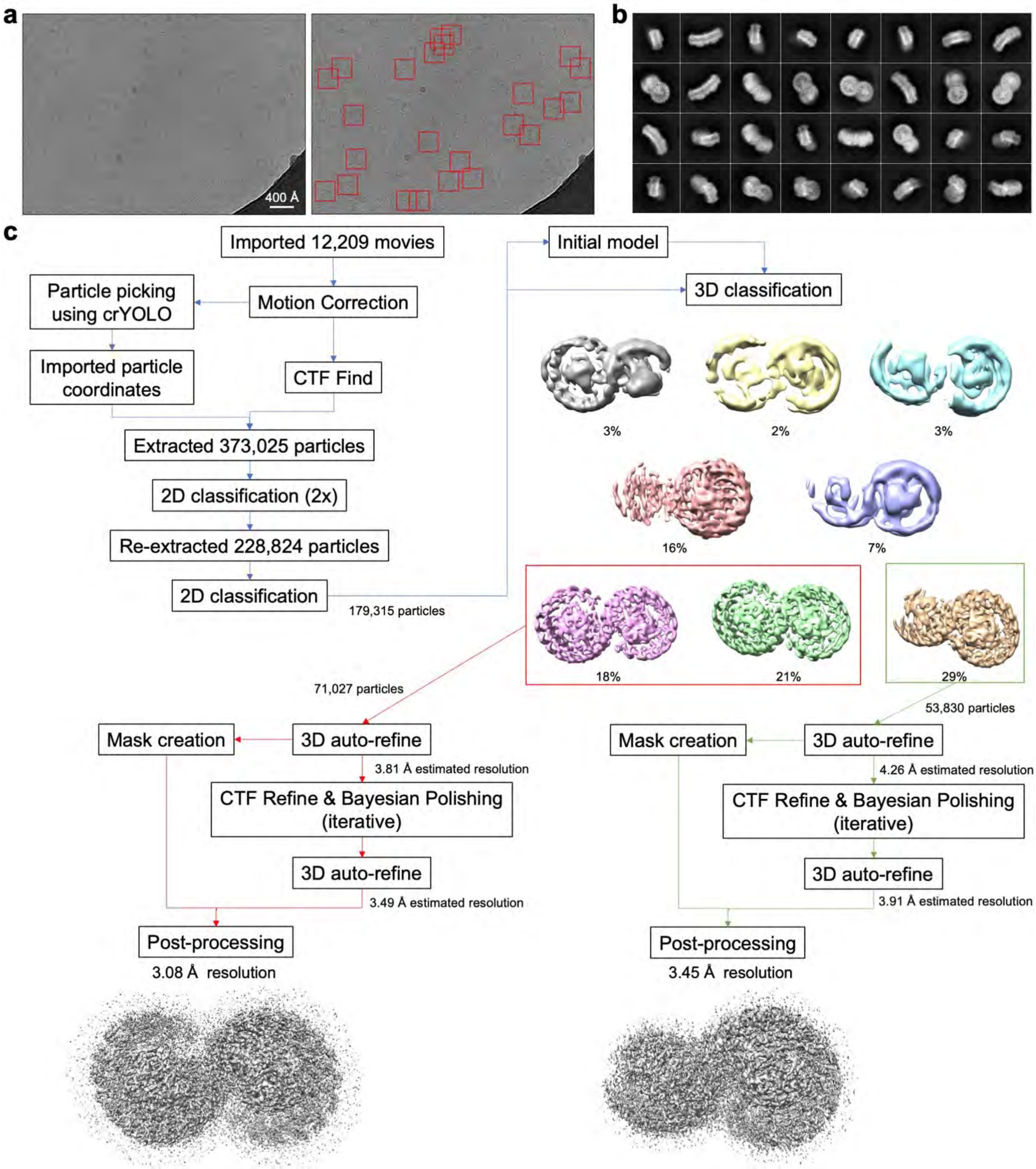


**Fig. S26. Cryo-EM analysis of Δ*pufY* RC–LH1 dimers.** (**a**) Motion-corrected example of a cryo-EM captured movie is shown on top and all particles picked from that micrograph are marked with red squares on the right. (**b**) Representative reference-free 2D class averages. (**c**) Overview of cryo-EM data processing workflow for the Δ*pufY* RC–LH1 dimer dataset. Every major step is shown with the associated number of particles, percentage of particles per class, or estimated resolution. Selected 3D classes that went into further processing are marked with rectangles (red – complete class, green – incomplete class). Two distinct classes were identified and separately refined, polished and sharpened.

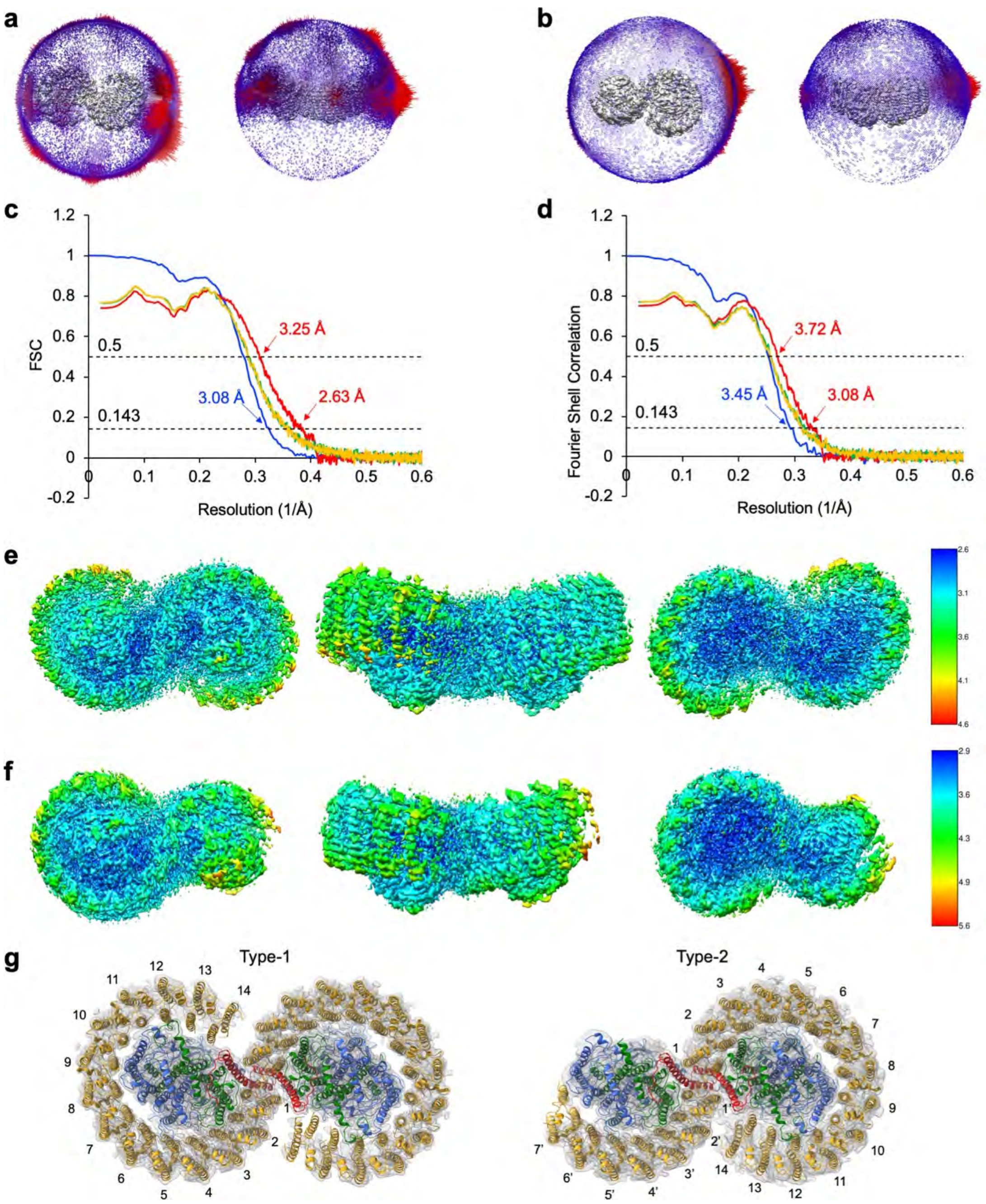


**Fig. S27. Cryo-EM data processing of Δ*pufY* RC–LH1 dimers.** (**a**-**b**) Angular distribution of the particles used to generate the final electron density maps. The direction and length of each cylinder represent the view and the number of particles respectively. (**a**) Complete (Type-1) dimer. (**b**) Incomplete (Type-2) dimer. (**c**) FSC curves of a complete (Class-1) dimer including FSC between two independently refined half-maps generated by RELION 3.1 (blue) and model-to-map FSC generated by Phenix (red). Global resolutions of 3.08 Å and 2.63 Å were calculated using the FSC cut-off of 0.143. (**d**) FSC curves of an incomplete (Type-2) dimer including FSC between two independently refined half-maps generated by RELION 3.1 (blue) and model-to-map FSC generated by Phenix (red). Green line: FSC of the model refined against the 1$^{st}$ half map versus the 1$^{st}$ half map; Yellow line: FSC of the model refined against the 1$^{st}$ half map versus the 2$^{nd}$ half map. Global resolutions of 3.45 Å and 3.08 Å were calculated using the FSC cut-off of 0.143. (**e**-**f**) Local resolution of the cryo-EM maps as seen from the cytoplasmic view (left), side view (middel), and the periplasmic view (right), estimated by RELION 3.1. (**e**) Complete (Type-1) dimer. (**f**) Incomplete (Type-2) dimer. (**g**) Cryo-EM map densities and structural model of protein peptides in the Δ*pufY* RC–LH1 dimers. Typ-1 Δ*pufY* RC–LH1 dimer (left) shows a 2-fold symmetry with 14 LH1 subunits in each monomer. Type-2 dimer adopts an asymmetric structure, with one monomer consisting of 14 LH1 subunits and the other having only 7-9 pairs of LH1 αβ-polypeptides. The last several pairs of LH1 in Type-2 dimer show weaker densities than other LH1s.

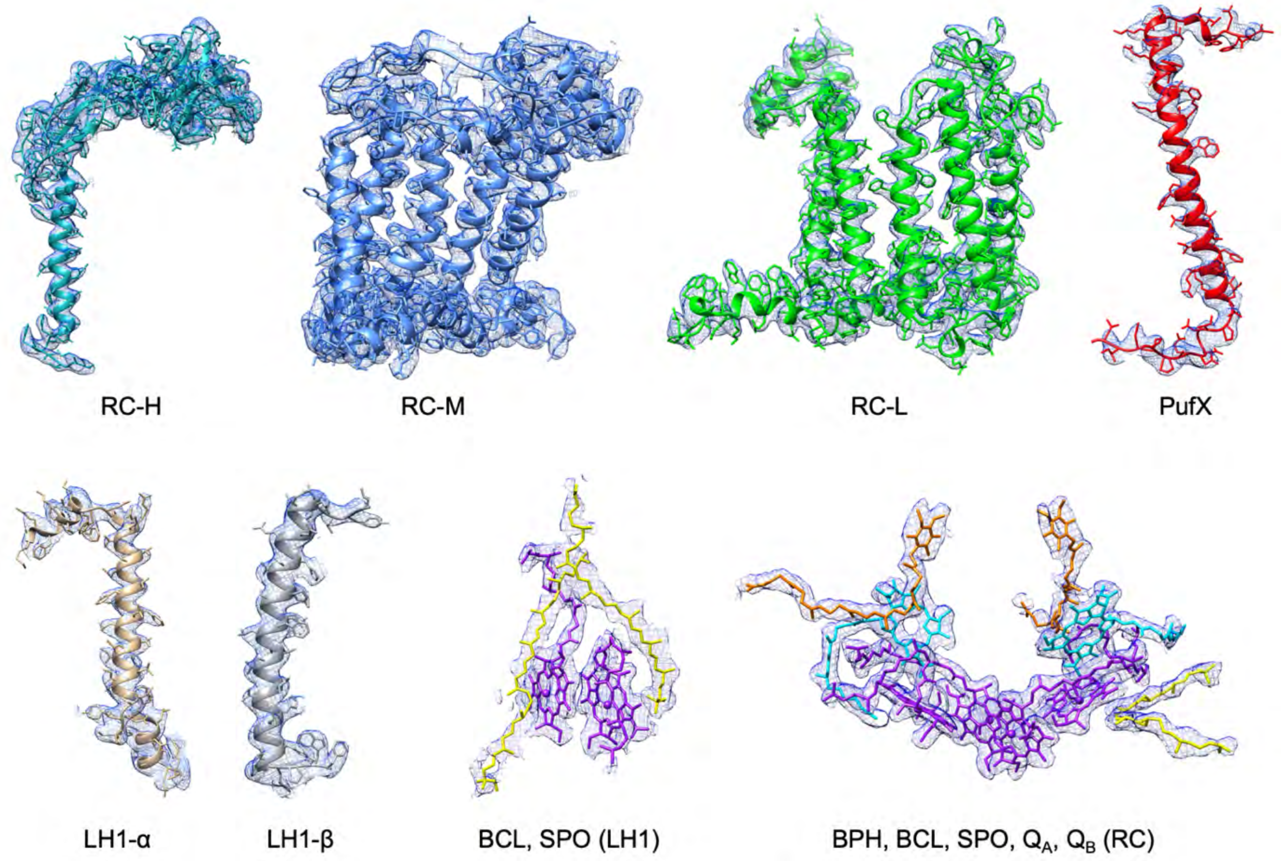


**Fig. S28. Cryo-EM map densities and structural models of protein peptides and cofactors in the Δ*pufY* RC–LH1 Type-1 dimer.**

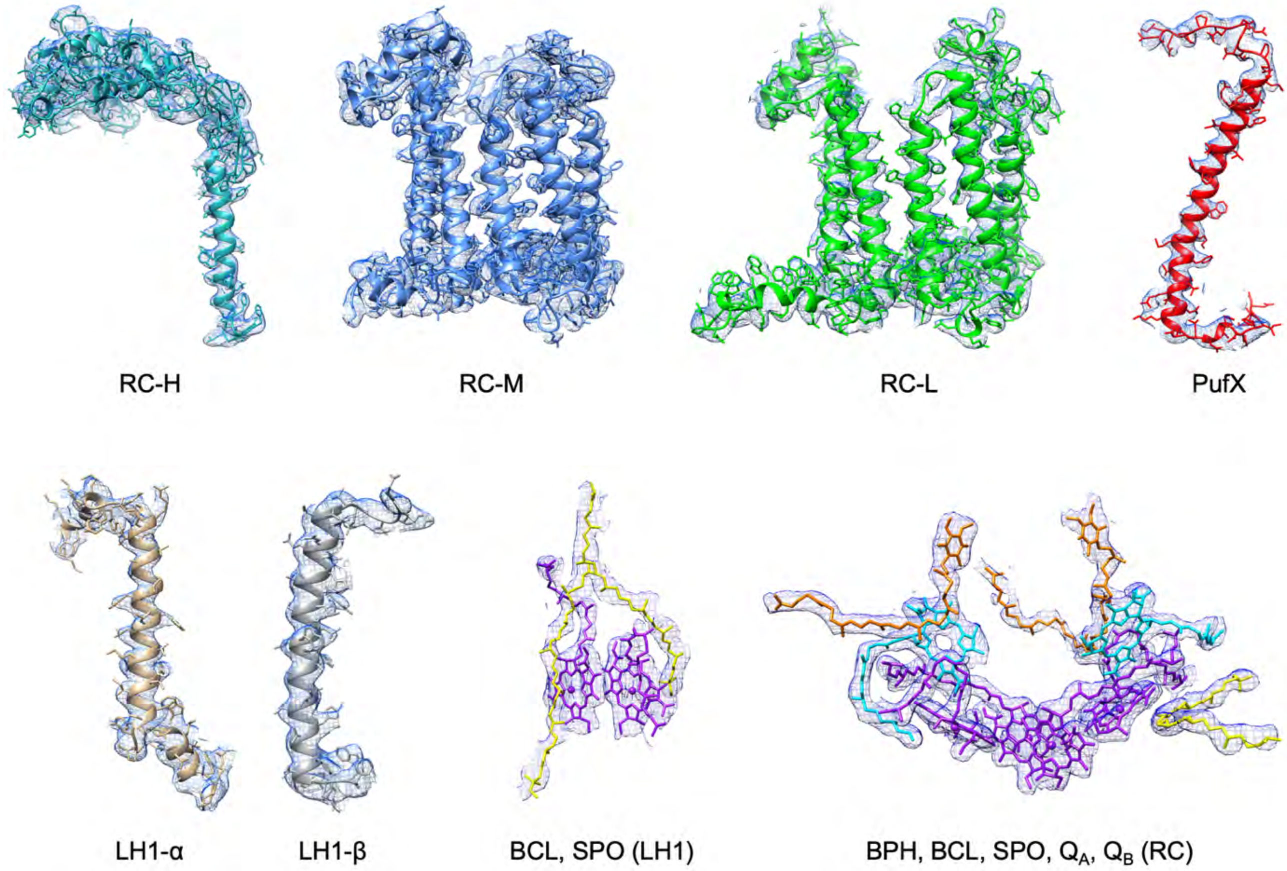


**Fig. S29. Cryo-EM map densities and structural models of protein peptides and cofactors in the Δ*pufY* RC–LH1 Type-2 dimer.**

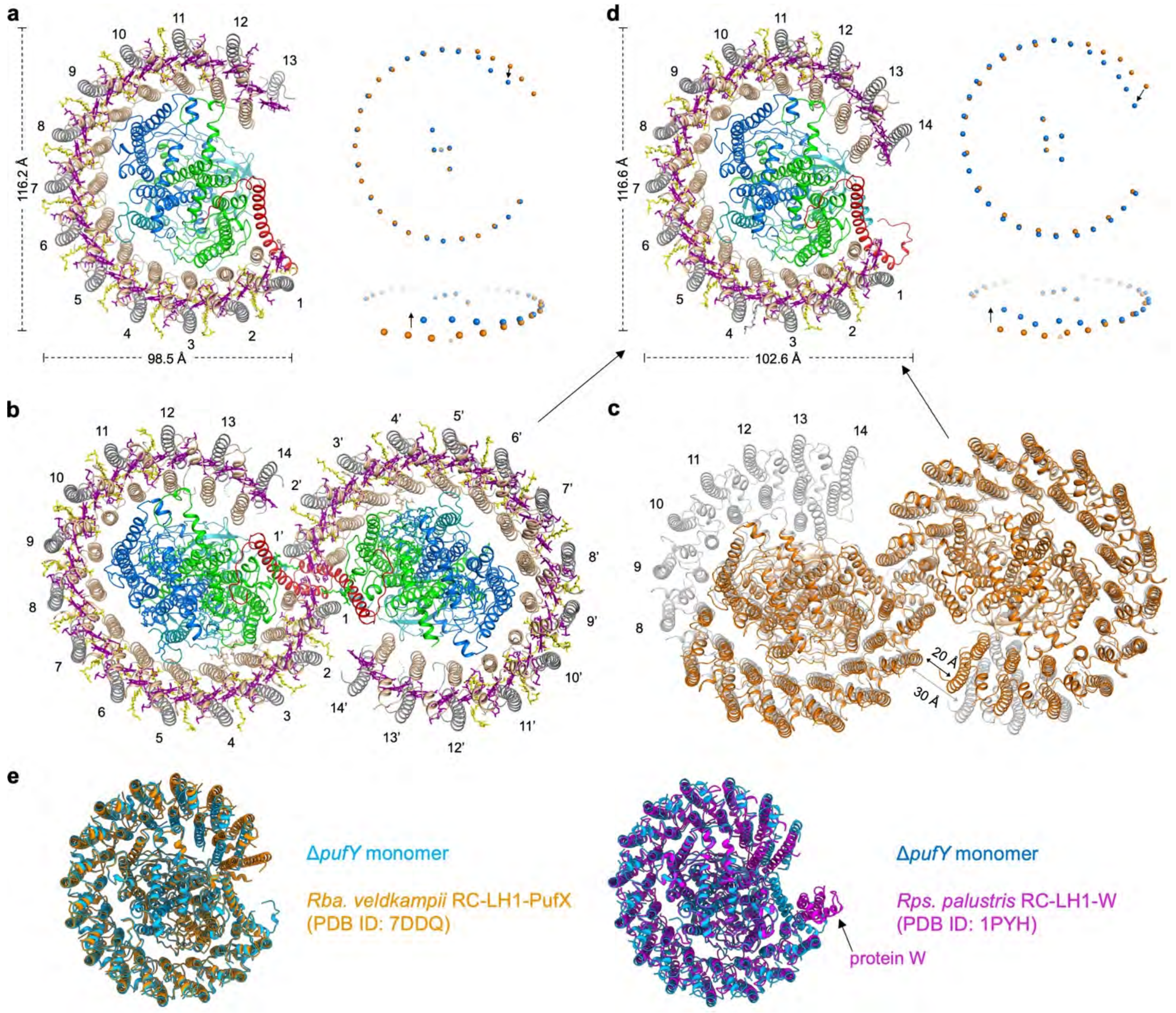


**Fig. S30. Δ*pufY* RC–LH1 monomeric and dimeric structures.** (**a**) Δ*pufY* RC–LH1 monomer that consists of RC associated with PufX and 13 LH1 subunits. The complex contains an elliptical LH1 ring (116 Å × 98.5 Å) with an enlarged opening in the LH1 array. Right, comparison of the Mg locations in the WT (orange) and Δ*pufY* (blue) RC–LH1 monomers. The absence of PufY caused the organizational shifts of LH1 terminal subunits and the lack of the $14^{th}$ LH1 subunits. (**b**) Δ*pufY* Class-1 RC–LH1 dimer, with a 2-fold symmetry of 14 LH1 subunits and two PufX located at the monomer-monomer interface. (**c**) Δ*pufY* Class-2 RC–LH1 dimer, with an asymmetric structure and two PufX located at the monomer-monomer interface. One monomer has 14 LH1 subunits and the other has an incomplete LH1 array with 7 LH1 subunits modeled in the structure. (**d**) The monomeric RC–LH1 structure from the Δ*pufY* RC–LH1 dimers, which consists of a more closed, elliptical LH1 ring of 14 subunits (117 Å × 103 Å). Right, comparison of the Mg locations in the WT (orange) and Δ*pufY* (blue) RC–LH1 monomers. The absence of PufY caused the organizational shifts of LH1 terminal subunits. (**e**) In the absence of PufY, the organization of LH1 subunits in the Δ*pufY* RC–$LH1_{14}$ monomer is more comparable to that of the *Rba. veldkampii* RC−LH1−PufX monomer (left, PDB ID: 7DDQ) than that of the *Rps. palustris* RC−LH1 (right, PDB ID: 1PYH) containing a protein W (arrow).

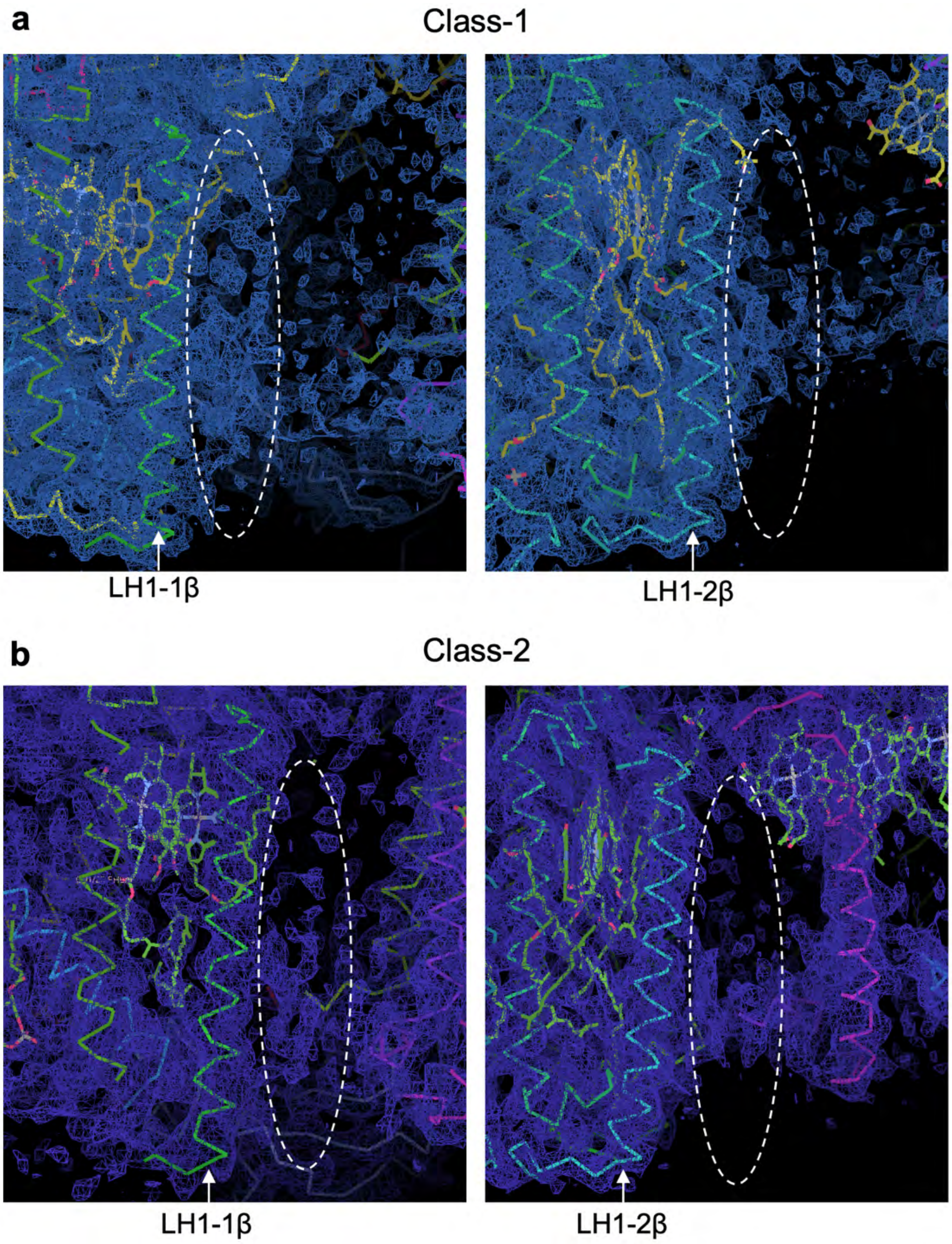


**Fig. S31. Cryo-EM density maps showing the densities at the potential locations of "protein-Z" [15] (indicated by dash circles) near LH1-1β (left) and LH1-2β (right).** (**a**) Local densities in the WT Class-1 dimer structure. (**b**) Local densities in the WT Class-2 dimer structure.

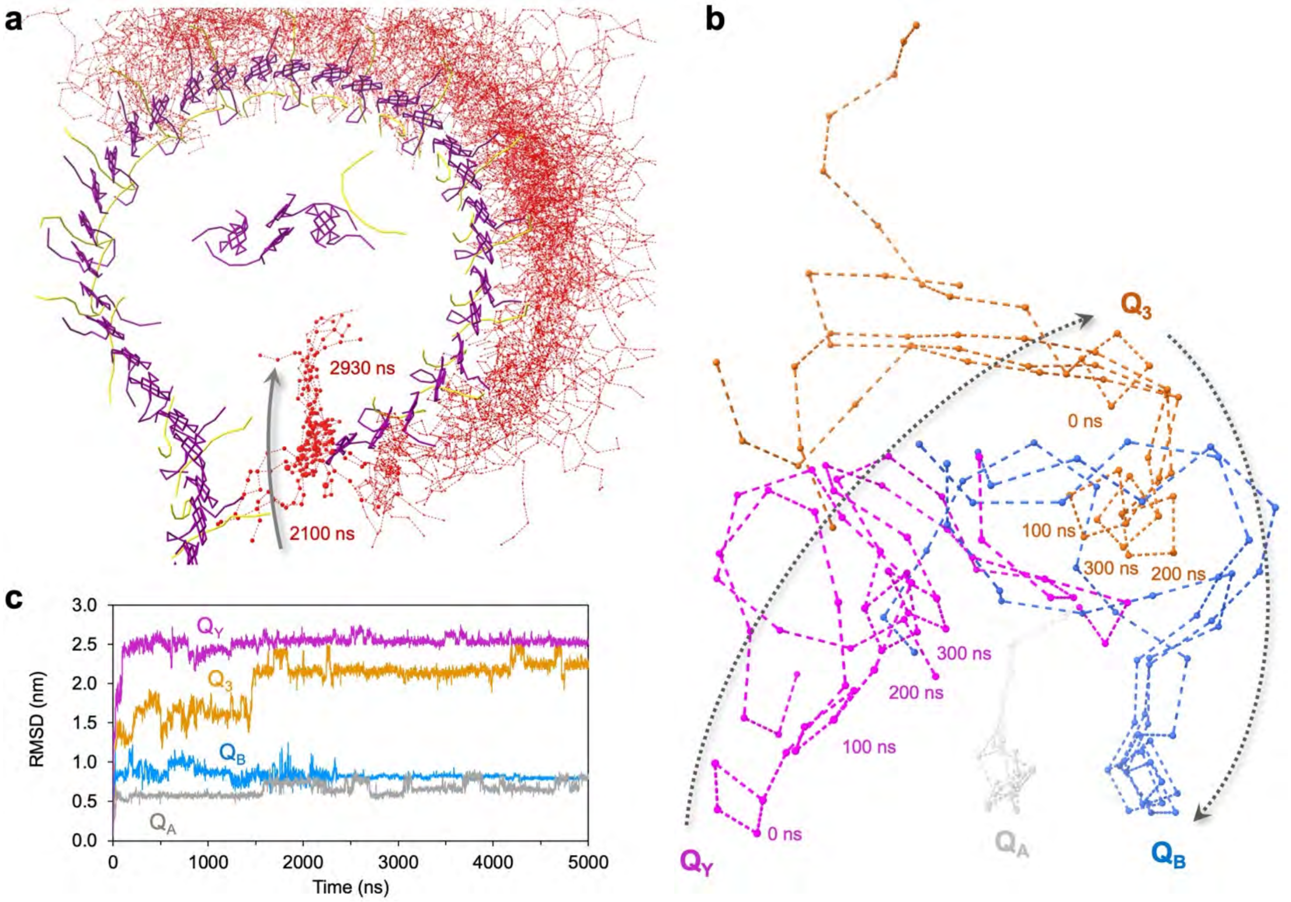


**Fig. S32. MD simulations of quinone movement within the dimeric RC−LH1 complex.** (**a**) CGMD simulations at the 5 μs timescale using a quinone-free RC−LH1 dimeric structure show quinones can enter the RC−LH1 complex through the large opening of the S-shaped LH1 ring as the predominant quinone transport pathway. (**b**) CGMD simulations using the RC−LH1 dimeric structure bound with $Q_A$ (grey), $Q_B$ (blue), $Q_3$ (orange), and $Q_Y$ (magenta) show the movement and conformational shifts of $Q_Y$ and $Q_3$. Dashed arrows imply the possible movement of $Q_Y$ towards $Q_3$ and $Q_3$ towards $Q_B$ within 300 ns. (**c**) Root-mean-square deviation (RMSD) analysis show the high conformational shifts of quinones in the RC−LH1 dimer, particularly at the early stage of 5000 ns simulations. $Q_A$ and $Q_B$ are relatively stable, whereas $Q_Y$ exhibits a greater conformational flexibility, likely explaining why $Q_Y$ is unresolved in the Class-1 RC−LH1 dimer.

```
RC-L            MALLSFERKYRVPGGTLVGGNLFDFWVGPFYVGFFGVATFFFAALGIILIAWSAVLQGTWNPQLISVYPPALEYGLGGAPLAKGGLW
                QIITICATGAFVSWALREVEICRKLGIGYHIPFAFAFAILAYLTLVLFRPVMMGAWGYAFPYGIWTHLDWVSNTGYTYGNFHYNPAH
                MIAISFFFTNALALALHGALVLSAANPEKGKEMRTPDHEDTFFRDLVGYSIGTLGIHRLGLLLSLSAVFFSALCMIITGTIWFDQWV
                DWWQWWVKLPWWANIPGGING

RC-M            MAEYQNIFSQVQVRGPADLGMTEDVNLANRSGVGPFSTLLGWFGNAQLGPIYLGSLGVLSLFSGLMWFFTIGIWFWYQAGWNPAVFL
                RDLFFFSLEPPAPEYGLSFAAPLKEGGLWLIASFFMFVAVWSWWGRTYLRAQALGMGKHTAWAFLSAIWLWMVLGFIRPILMGSWSE
                AVPYGIFSHLDWTNNFSLVHGNLFYNPFHGLSIAFLYGSALLFAMHGATILAVSRFGGERELEQIADRGTAAERAALFWRWTMGFNA
                TMEGIHRWAIWMAVLVTLTGGIGILLSGTVVDNWYVWGQNHGMAPLN

RC-H            MVGVTAFGNFDLASLAIYSFWIFLAGLIYYLQTENMREGYPLENEDGTPAANQGPFPLPKPKTFILPHGRGTLTVPGPESEDRPIAL
                ARTAVSEGFPHAPTGDPMKDGVGPASWVARRDLPELDGHGHNKIKPMKAAAGFHVSAGKNPIGLPVRGCDLEIAGKVVDIWVDIPEQ
                MARFLEVELKDGSTRLLPMQMVKVQSNRVHVNALSSDLFAGIPTIKSPTEVTLLEEDKICGYVAGGLMYAAPKRKSVVAAMLAEYA

Class-1 dimer   MVGVTAFGNFDLASLAIYSFWIFLAGLIYYLQTENMREGYPLENEDGTPAANQGPFPLPKPKTFILPHGRGTLTVPGPESEDRPIAL
                ARTAVSEGFPHAPTGDPMKDGVGPASWVARRDLPELDGHGHNKIKPMKAAAGFHVSAGKNPIGLPVRGCDLEIAGKVVDIWVDIPEQ
                MARFLEVELKDGSTRLLPMQMVKVQSNRVHVNALSSDLFAGIPTIKSPTEVTLLEEDKICGYVAGLMYAAPKRKSVVAAMLAEYA

Class-2 dimer   MVGVTAFGNFDLASLAIYSFWIFLAGLIYYLQTENMREGYPLENEDGTPAANQGPFPLPKPKTFILPHGRGTLTVPGPESEDRPIAL
                ARTAVSEGFPHAPTGDPMKDGVGPASWVARRDLPELDGHGHNKIKPMKAAAGFHVSAGKNPIGLPVRGCDLEIAGKVVDIWVDIPEQ
                MARFLEVELKDGSTRLLPMQMVKVQSNRVHVNALSSDLFAGIPTIKSPTEVTLLEEDKICGYVAGLMYAAPKRKSVVAAMLAEYA

WT monomer      MVGVTAFGNFDLASLAIYSFWIFLAGLIYYLQTENMREGYPLENEDGTPAANQGPFPLPKPKTFILPHGRGTLTVPGPESEDRPIAL
                ARTAVSEGFPHAPTGDPMKDGVGPASWVARRDLPELDGHGHNKIKPMKAAAGFHVSAGKNPIGLPVRGCDLEIAGKVVDIWVDIPEQ
                MARFLEVELKDGSTRLLPMQMVKVQSNRVHVNALSSDLFAGIPTIKSPTEVTLLEEDKICGYVAGLMYAAPKRKSVVAAMLAEYA

LH1α            MSKFYKIWMIFDPRRVFVAQGVFLFLLAVMIHLILLSTPSYNWLEISAAKYNRVAVAE
Class-1 dimer   MSKFYKIWMIFDPRRVFVAQGVFLFLLAVMIHLILLSTPSYNWLEISAAKYNRVAVAE
Class-2 dimer   MSKFYKIWMIFDPRRVFVAQGVFLFLLAVMIHLILLSTPSYNWLEISAAKYNRVAVAE
WT monomer      MSKFYKIWMIFDPRRVFVAQGVFLFLLAVMIHLILLSTPSYNWLEISAAKYNRVAVAE

LH1β            MADKSDLGYTGLTDEQAQELHSVYMSGLWLFSAVAIVAHLAVYIWRPWF
Class-1 dimer   MADKSDLGYTGLTDEQAQELHSVYMSGLWLFSAVAIVAHLAVYIWRPWF
Class-2 dimer   MADKSDLGYTGLTDEQAQELHSVYMSGLWLFSAVAIVAHLAVYIWRPWF
WT monomer      MADKSDLGYTGLTDEQAQELHSVYMSGLWLFSAVAIVAHLAVYIWRPWF

PufX            MADKTIFNDHLNTNPKTNLRLWVAFQMMKGAGWAGGVFFGTLLLIGFFRVVGRMLPIQENQAPAPNITGALETGIELIKHLV
Class-1 dimer   MADKTIFNDHLNTNPKTNLRLWVAFQMMKGAGWAGGVFFGTLLLIGFFRVVGRMLPIQENQAPAPNITGALETGIELIKHLV
Class-2 dimer   MADKTIFNDHLNTNPKTNLRLWVAFQMMKGAGWAGGVFFGTLLLIGFFRVVGRMLPIQENQAPAPNITGALETGIELIKHLV
WT monomer      MADKTIFNDHLNTNPKTNLRLWVAFQMMKGAGWAGGVFFGTLLLIGFFRVVGRMLPIQENQAPAPNITGALETGIELIKHLV

PufY            MPEVSEFAFRLMMAAVIFVGVGIMFAFAGGHWFVGLVVGGLVAAFFAATPNSN
Class-1 dimer   MPEVSEFAFRLMMAAVIFVGVGIMFAFAGGHWFVGLVVGGLVAAFFAATPNSN
Class-2 dimer   MPEVSEFAFRLMMAAVIFVGVGIMFAFAGGHWFVGLVVGGLVAAFFAATPNSN
WT monomer      MPEVSEFAFRLMMAAVIFVGVGIMFAFAGGHWFVGLVVGGLVAAFFAATPNSN
```

**Fig. S33. Amino acid sequences of protein polypeptides in the RC−LH1 complex from *Rba. sphaeroides*, and resolved residues in the dimeric and monomeric structures.** Protein residues that were not resolved in the structures of Class-1 and Class-2 dimers and the WT monomer are highlighted in red. Protein Uniprot ID: RC-L, Q3J1A5; RC-M, Q3J1A6; RC-H, Q3J170; LH1-α, Q3J1A4; LH1-β, Q3J1A3; PufX, P13402; PufY, U5NME9.

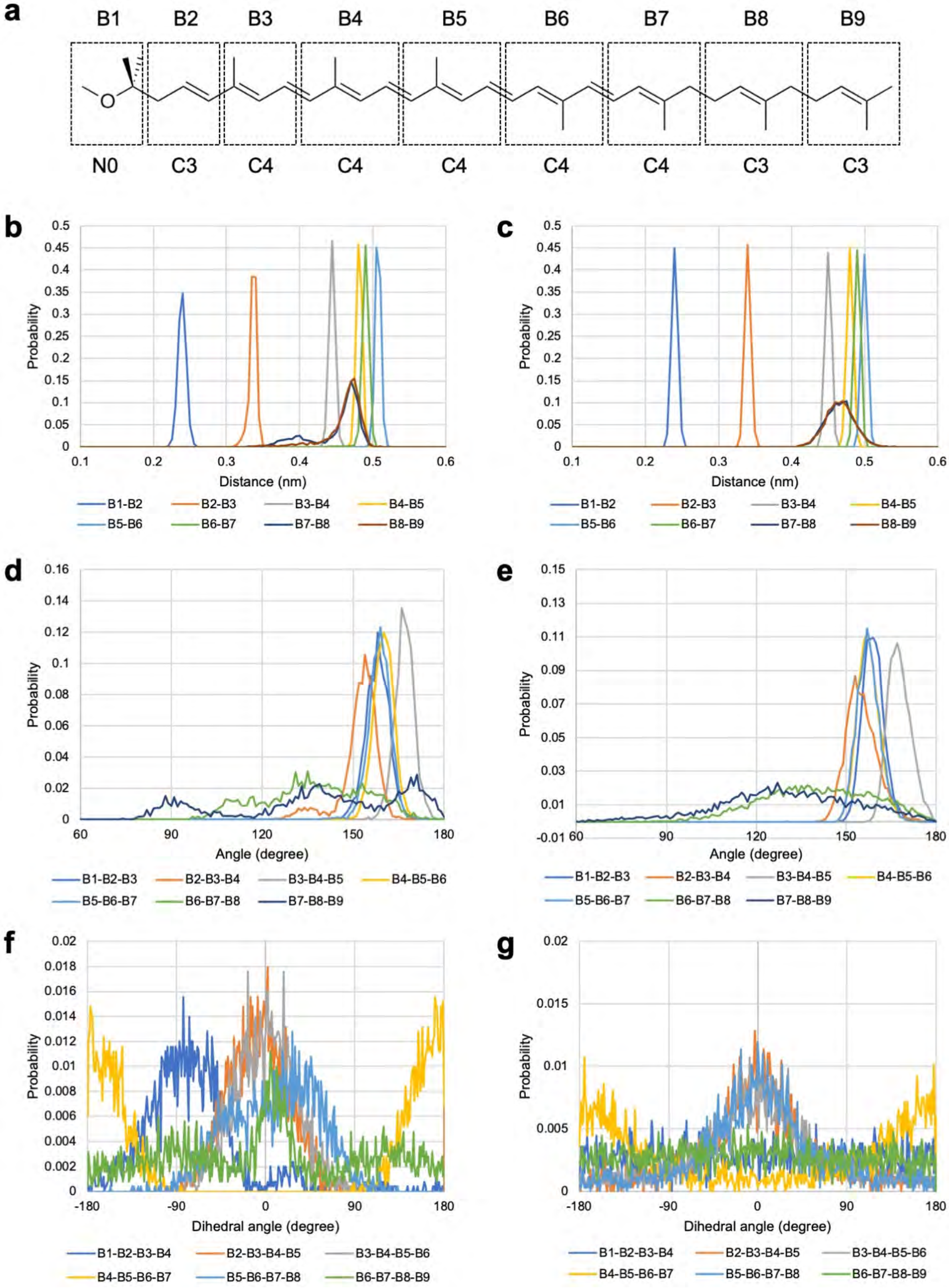


**Fig. S34. Setting the coarse-grained models of cofactors.** (**a**) Mapping of SPO atoms to coarse-grained (CG) beads. A CG bead was placed at the center of mass of the atoms within a dashed square. The name and the type of each CG bead are shown above and below the structure, respectively. (**b-g**) Comparison of the probability distributions of bond lengths (**a**, **c**), angles (**d**, **e**), and dihedral angles (**f**, **g**) between the AAMD (**b**, **d**, **f**) and CGMD (**c**, **e**, **f**) simulations.

### SUPPLEMENTARY TABLES

**Table S1. Cryo-EM data collection, refinement and validation statistics (to be continued).**

| | WT dimer Class-1 | WT dimer Class-2 | WT monomer |
|---|---|---|---|
| **Data collection and processing** | | | |
| Magnification | 130,000 x | - | 81,000 x |
| Voltage (kV) | 300 | - | 300 |
| Electron exposure ($e^{–}$/Å$^2$) | 60 | - | 45.026 |
| Defocus range (μm) | 1.8-2.2 | - | 0.8-1.8 |
| Pixel size (Å) | 1.04 | - | 1.06 |
| Symmetry imposed | C2 | C2 | C1 |
| Initial particle images (no.) | 957,176 | - | 241,170 |
| Final particle images (no.) | 145,392 | 115,388 | 68,554 |
| Map resolution (Å) | 2.74 | 2.98 | 2.79 |
| FSC threshold | 0.143 | 0.143 | 0.143 |
| Map resolution range (Å) | 2.5-5.3 | 2.7-4.8 | 2.3-4.5 |
| **Refinement** | | | |
| Initial model used (PDB code) | 5Y5S | Class-1 dimer | Class-2 dimer |
| Model resolution (Å) | 2.74 | 2.98 | 2.76 |
| FSC threshold | 0.143 | 0.143 | 0.143 |
| Map resolution range (Å) | - | - | - |
| Map sharpening *B* factor (Å$^2$) | -41.0 | -50.6 | -86.3 |
| Model composition | | | |
| Non-hydrogen atoms | 44,980 | 44,992 | 22,506 |
| Protein residues | 4,614 | 4,596 | 2,284 |
| Ligands | 148 | 152 | 76 |
| *B* factors (Å$^2$) | | | |
| Protein | 38.6 | 53.3 | 68.98 |
| Ligand | 39.3 | 45.4 | 63.88 |
| R.m.s. deviations | | | |
| Bond lengths (Å) | 0.006 | 0.006 | 0.012 |
| Bond angles (°) | 0.917 | 0.922 | 1.159 |
| **Validation** | | | |
| MolProbity score | 1.45 | 1.51 | 1.64 |
| Clashscore | 8.3 | 9.8 | 10.46 |
| Poor rotamers (%) | 0.37 | 0.37 | 0.05 |
| Ramachandran plot | | | |
| Favored (%) | 98.2 | 98.3 | 97.47 |
| Allowed (%) | 1.8 | 1.7 | 2.53 |
| Disallowed (%) | 0.00 | 0.00 | 0.00 |

**Table S1. Cryo-EM data collection, refinement and validation statistics (continued).**

| | **Δ*pufX* monomer** | **Δ*pufY* monomer** | **Δ*pufY* dimer Type-1** | **Δ*pufY* dimer Type-2** |
|---|---|---|---|---|
| **Data collection and processing** | | | | |
| Magnification | 81,000 x | 105,000 x | 105,000 x | - |
| Voltage (kV) | 300 | 300 | 300 | - |
| Electron exposure ($e^{-}$/Å$^2$) | 46.549 | 51.527 | 50.868 | - |
| Defocus range (μm) | 0.8-1.8 | 0.8-2.0 | 0.8-2.0 | - |
| Pixel size (Å) | 1.06 | 0.8285 | 0.8285 | - |
| Symmetry imposed | C1 | C1 | C1 | C1 |
| Initial particle images (no.) | 3,420,815 | 241,170 | 373,025 | - |
| Final particle images (no.) | 66,058 | 56,391 | 71,027 | 53,830 |
| Map resolution (Å) | 4.20 | 2.86 | 3.08 | 3.45 |
| FSC threshold | 0.143 | 0.143 | 0.143 | 0.143 |
| Map resolution range (Å) | 3.8-6.2 | 2.5-4.5 | 2.70-4.63 | 2.99-5.66 |
| **Refinement** | | | | |
| Initial model used (PDB code) | WT monomer | WT monomer | WT Class-1 dimer | Δ*pufY* RC−LH1 Type-1 dimer |
| Model resolution (Å) | 3.55 | 2.43 | 2.62 | 3.08 |
| FSC threshold | 0.143 | 0.143 | 0.143 | 0.143 |
| Map resolution range (Å) | - | - | - | - |
| Map sharpening *B* factor (Å$^2$) | -163.9 | -94.3 | -122.5 | -122.7 |
| Model composition | | | | |
| Non-hydrogen atoms | 22,119 | 20,589 | 42,176 | 36,469 |
| Protein residues | 2,479 | 2,112 | 4,350 | 3,807 |
| Ligands | 44 | 65 | 131 | 107 |
| *B* factors (Å$^2$) | | | | |
| Protein | 77.32 | 25.32 | 23.27 | 33.61 |
| Ligand | 68.62 | 22.74 | 26.66 | 34.48 |
| R.m.s. deviations | | | | |
| Bond lengths (Å) | 0.008 | 0.007 | 0.007 | 0.008 |
| Bond angles (°) | 0.929 | 1.006 | 1.069 | 0.912 |
| **Validation** | | | | |
| MolProbity score | 2.10 | 1.57 | 2.09 | 2.23 |
| Clashscore | 15.97 | 10.03 | 15.42 | 15.18 |
| Poor rotamers (%) | 1.25 | 0.62 | 2.78 | 3.26 |
| Ramachandran plot | | | | |
| Favored (%) | 95.38 | 97.81 | 97.70 | 97.11 |
| Allowed (%) | 4.58 | 2.19 | 2.30 | 2.86 |
| Disallowed (%) | 0.04 | 0.00 | 0.00 | 0.03 |

**Table S2. Components of the WT RC−LH1 monomer and dimers from *Rba. sphaeroides*.**

| Proteins | | Numbers | | | Cofactors | Numbers | | |
|---|---|---|---|---|---|---|---|---|
| | | **Monomer** | **Dimer Class-1** | **Dimer Class-2** | | **Monomer** | **Dimer Class-1** | **Dimer Class-2** |
| LH1 | α subunit | 14 | 28 | 28 | BChl *a* | 28 | 56 | 56 |
| | β subunit | 14 | 28 | 28 | Spheroidene (SPO) | 25 | 50 | 50 |
| RC | H | 1 | 2 | 2 | BChl *a* | 4 | 8 | 8 |
| | L | 1 | 2 | 2 | Spheroidene (SPO) | 1 | 2 | 2 |
| | M | 1 | 2 | 2 | BPhe | 2 | 4 | 4 |
| | | | | | $Fe^{2+}$ | 1 | 2 | 2 |
| | | | | | UQ-10 | 4 | 6 | 8 |
| | | | | | Cardiolipin (CDL) | 2 | 4 | 4 |
| | | | | | Phosphatidylcholine (PC1) | 9 | 16 | 18 |
| PufX | | 1 | 2 | 2 | | | | |
| PufY | | 1 | 2 | 2 | | | | |
| **Total** | | **33** | **66** | **66** | **Total** | **76** | **148** | **152** |

**Table S3. Interactions within the RC−LH assembly.**

RC-M:LH1

| | Structure 1 | Structure 2 | Dist. [Å] | Interaction |
|---|---|---|---|---|
| 1 | M:ALA27[O] | 9α:ARG15[NH2] | 2.8 | Hydrogen bond |
| 2 | M:SER54[OG] | 8α:ARG15[NH1] | 3.3 | Hydrogen bond |
| 3 | M:ASN81[ND2] | 11α:SER37[OG] | 2.7 | Hydrogen bond |
| 4 | M:TYR134[OH] | 7α:ARG15[NH1] | 3.4 | Hydrogen bond |

RC-L:LH1

| | Structure 1 | Structure 2 | Dist. [Å] | Interaction |
|---|---|---|---|---|
| 1 | L:TRP25[O] | 2α:ARG15[NH2] | 3.0 | Hydrogen bond |
| 2 | L:TRP25[O] | 2α:ARG15[NE] | 3.4 | Hydrogen bond |
| 3 | L:TRP51[NE1] | 2α:SER37[OG] | 3.0 | Hydrogen bond |
| 4 | L:TRP59[NE1] | 3α:SER37[OG] | 3.1 | Hydrogen bond |
| 5 | L:ALA78[O] | 1α:SER37[OG] | 2.9 | Hydrogen bond |

RC-H:LH1

| | Structure 1 | Structure 2 | Dist. [Å] | Interaction |
|---|---|---|---|---|
| 1 | H:ALA6[O] | 6α:ASN42[ND2] | 3.5 | Hydrogen bond |
| 2 | H:GLY8[N] | 5α:SER37[OG] | 3.4 | Hydrogen bond |
| 3 | H:GLY54[O] | 4α:ARG15[NH1] | 2.2 | Hydrogen bond |
| 4 | H:PHE56[O] | 4α:ARG15[NH1] | 3.2 | Hydrogen bond |
| 5 | H:SER93[OG] | 2α:ARG15[NH1] | 3.4 | Hydrogen bond |
| 6 | H:GLU94[O] | 2α:ARG15[NH1] | 2.6 | Hydrogen bond |
| 7 | H:ALA260[OXT] | 1α:ARG14[NH2] | 3.3 | Hydrogen bond |

LH1-1α:LH1-1β

| | Structure 1 | Structure 2 | Dist. [Å] | Interaction |
|---|---|---|---|---|
| 1 | 1α:TYR5[OH] | 1β:ASP14[OD1] | 3.2 | Hydrogen bond |
| 2 | 1α:TYR5[OH] | 1β:ASP14[OD2] | 3.1 | Hydrogen bond |
| 3 | 1α:TRP8[O] | 1β:THR10[OG1] | 3.0 | Hydrogen bond |
| 4 | 1α:MET9[O] | 1β:TYR9[N] | 3.0 | Hydrogen bond |
| 5 | 1α:GLN20[OE1] | 1β:TYR24[OH] | 2.3 | Hydrogen bond |
| 6 | 1α:GLN20[NE2] | 1β:TYR24[OH] | 3.2 | Hydrogen bond |
| 7 | 1α:TYR41[OH] | 1β:ARG46[O] | 3.0 | Hydrogen bond |
| 8 | 1α:TYR41[OH] | 1β:PRO47[O] | 2.9 | Hydrogen bond |

LH1-1α:LH1-2α

| | Structure 1 | Structure 2 | Dist. [Å] | Interaction |
|---|---|---|---|---|
| 1 | 1α:TYR41[OH] | 2α:ARG53[NH1] | 2.9 | Hydrogen bond |

**Table S4. PufX interactions in the Class-1 RC−LH1 dimer.** Note: no direct PufX-PufX interactions were identified.

| | Structure 1 | Structure 2 | Dist. [Å] | Interaction |
|---|---|---|---|---|
| PufX:L | X:ARG49[NH2] | L:VAL137[O] | 2.9 | Hydrogen bond |
| | X:ARG49[NH2] | L:MET138[O] | 3.2 | Hydrogen bond |
| | X:ALA62[O] | L:TYR144[OH] | 2.6 | Hydrogen bond |
| | X:PRO65[O] | L:ASN159[ND2] | 2.5 | Hydrogen bond |
| | X:ASN66[OD1] | L:GLY143[O] | 3.2 | Hydrogen bond |
| | X:ASN66[ND2] | L:ALA145[O] | 2.6 | Hydrogen bond |
| PufX:H | X:LYS29[NZ] | H:TYR259[O] | 2.3 | Hydrogen bond |
| PufX:H (neighbouring monomer) | X:ASN14[OD1] | h:GLU258[O] | 2.8 | Hydrogen bond |
| | X:THR17[OG1] | h:GLU258[O] | 2.4 | Hydrogen bond |
| | X:ARG20[NH1] | h:TYR259[OH] | 2.5 | Hydrogen bond |
| | X:ARG20[NH2] | h:GLU258[OE2] | 3.3 | Salt bridge |
| PufX:LH1-1α (neighbouring monomer) | X:ASP9[O] | 1α:ARG14[NH2] | 3.6 | Hydrogen bond |
| | X:ASP9[OD2] | 1α:ARG14[NE] | 3.9 | Salt bridge |
| | X:ASP9[OD1] | 1α:ARG14[NH2] | 3.8 | Salt bridge |
| PufX:LH1-1β (neighbouring monomer) | X:ILE6[O] | 1β:GLN16[NE2] | 2.8 | Hydrogen bond |
| | X:ILE6[N] | 1β:GLU19[OE1] | 2.8 | Hydrogen bond |
| | X:PHE7[O] | 1β:GLN16[NE2] | 3.1 | Hydrogen bond |
| | X:PHE7[N] | 1β:GLU19[OE1] | 3.1 | Hydrogen bond |
| | X:ARG53[NH2] | 1β:ILE44[O] | 2.9 | Hydrogen bond |
| | X:ARG53[NH1] | 1β:ILE44[O] | 2.8 | Hydrogen bond |

**Table S5. PufY interactions in the RC−LH1 monomer.**

| | Structure 1 | Structure 2 | Dist. [Å] | Interaction |
|---|---|---|---|---|
| PufY:LH1−14α | Y:Val4[O] | 14α:Arg15[NH1] | 3.4 | Hydrogen bond |
| | Y:Val4[O] | 14α:Arg15[NH2] | 3.3 | Hydrogen bond |
| | Y:Ser5[O] | 14α:Arg15[NH2] | 3.4 | Hydrogen bond |
| | Y:Glu6[OE1] | 14α:Arg15[NH2] | 3.9 | Salt bridge |
| PufY:LH1−13α | Y:Ala28[O] | 13α:Ser37[OG] | 3.3 | Hydrogen bond |
| | Y:Thr49[O] | 13α:Arg15[NH1] | 2.7 | Hydrogen bond |
| | Y:Pro50[O] | 13α:Arg15[NH1] | 3.0 | Hydrogen bond |
| | Y:Asn51[OD1] | 13α:Arg15[NE] | 3.0 | Hydrogen bond |
| PufY:RC-H | Y:Glu3[OE1] | H:Arg202[NH1] | 3.9 | Salt bridge |
